# Functional Immune Deficiency Syndrome via Intestinal Infection in COVID-19

**DOI:** 10.1101/2020.04.06.028712

**Authors:** Erica T. Prates, Michael R. Garvin, Mirko Pavicic, Piet Jones, Manesh Shah, Christiane Alvarez, David Kainer, Omar Demerdash, B Kirtley Amos, Armin Geiger, John Pestian, Kang Jin, Alexis Mitelpunkt, Eric Bardes, Bruce Aronow, Daniel Jacobson

## Abstract

Using a Systems Biology approach, we integrated genomic, transcriptomic, proteomic, and molecular structure information to provide a holistic understanding of the COVID-19 pandemic. The expression data analysis of the Renin Angiotensin System indicates mild nasal, oral or throat infections are likely and that the gastrointestinal tissues are a common primary target of SARS-CoV-2. Extreme symptoms in the lower respiratory system likely result from a secondary-infection possibly by a comorbidity-driven upregulation of ACE2 in the lung. The remarkable differences in expression of other RAS elements, the elimination of macrophages and the activation of cytokines in COVID-19 bronchoalveolar samples suggest that a functional immune deficiency is a critical outcome of COVID-19. We posit that using a non-respiratory system as a major pathway of infection is likely determining the unprecedented global spread of this coronavirus.

**One Sentence Summary:** A Systems Approach Indicates Non-respiratory Pathways of Infection as Key for the COVID-19 Pandemic

## Main Text

The COVID-19 beta-coronavirus epidemic that originated in Wuhan, China in December of 2019 is now a global pandemic and is having devastating societal and economic impacts. The increasing frequency of the emergence of zoonotics such as Ebola, Severe Acute Respiratory Syndrome (SARS), and Middle East Respiratory Syndrome (MERS) (among others) are of grave concern because of their high mortality rate (15% - 90%). Fortunately, successful containment of those pathogens prevented global-scale deaths. In contrast, the current estimates of mortality for COVID-19 are much lower (~4%), but it is now clear that the virus has not been contained, which may be due to higher rates of asymptomatic transmission of those infected with SARS-CoV-2 (*1*, *2*). Given that the SARS-CoV-2 virus has been detected worldwide, rapid measures are needed to address the expanding epidemic.

Paradoxically, an opportunity that was unavailable with SARS, MERS or Ebola has arisen because of the intense, globally distributed focus of medical and scientific professionals on COVID-19 that is providing a wealth of highly diverse information and data types. A Systems Biology approach done in combination with supercomputing can integrate these diverse sources of information to provide improved clarity and insights into the mechanisms of disease.

From the analysis of multi-omics data from the COVID-19 pandemic, we find that the critical outcomes of the disease involve the imbalance of several components of the Renin Angiotensin System (RAS) beyond the direct binding to angiotensin-converting enzyme 2 (ACE2) for cell entry. In addition, expression data from diverse sources and tissues suggests that the virus evolved to target the digestive system. We propose that a non-respiratory route of infection is a major contributor to the wide geographic spread of SARS-CoV-2 via “asymptomatic” carriers. Furthermore, the data suggest specific cells in the immune system are being repressed or destroyed in the lung, leading to a **functional immune deficiency syndrome (COVID-19-FIDS)**. By integrating extensive proteome structural analyses with the multi-omics data layers, the molecular differences between SARS-CoV-1 and SARS-CoV-2 were identified, and provide other elements that can explain the divergence in pathogenicity between SARS and COVID-19.

### Coronaviruses and Renin-Angiotensin-System (RAS)

Several species of coronavirus (with highly variable host mortality rates) have evolved to use proteases in the Renin Angiotensin System (RAS) to gain entry to host cells (*3*) including the current SARS-CoV-2 virus (Fig. 1A). Notably, in addition to ACE2, other RAS proteases, such as those that produce angiotensin IV (ANPEP) and angiotensin 1-7 (ACE), may be important in the context of COVID-19, given that they activate proinflammatory genes, which are components of the innate antiviral response and may be a key factor in mortality from coronaviruses (*4*). The expression profile of RAS genes of cells from bronchoalveolar lavage samples (BAL) taken from individuals with COVID-19 indicate upregulation of renin, angiotensin, ACE2, and MAS (Fig. 1B). Induction of this pathway from viral infection also indirectly implicates ACE, as it is needed to activate MAS via angiotensin 1-7. In addition, during the current pandemic, it may be informative to track several factors that are known to alter RAS, including drugs (R_X_ in Fig. 1A), outcomes of infection (*e.g.* dehydration from diarrhea), and genetic polymorphisms. For example, an *Alu* deletion in ACE is responsible for 40% of the variance in circulating levels of that enzyme (*5*) and the frequency of the allele varies considerably among populations (Table S1).

**Fig. 1.**
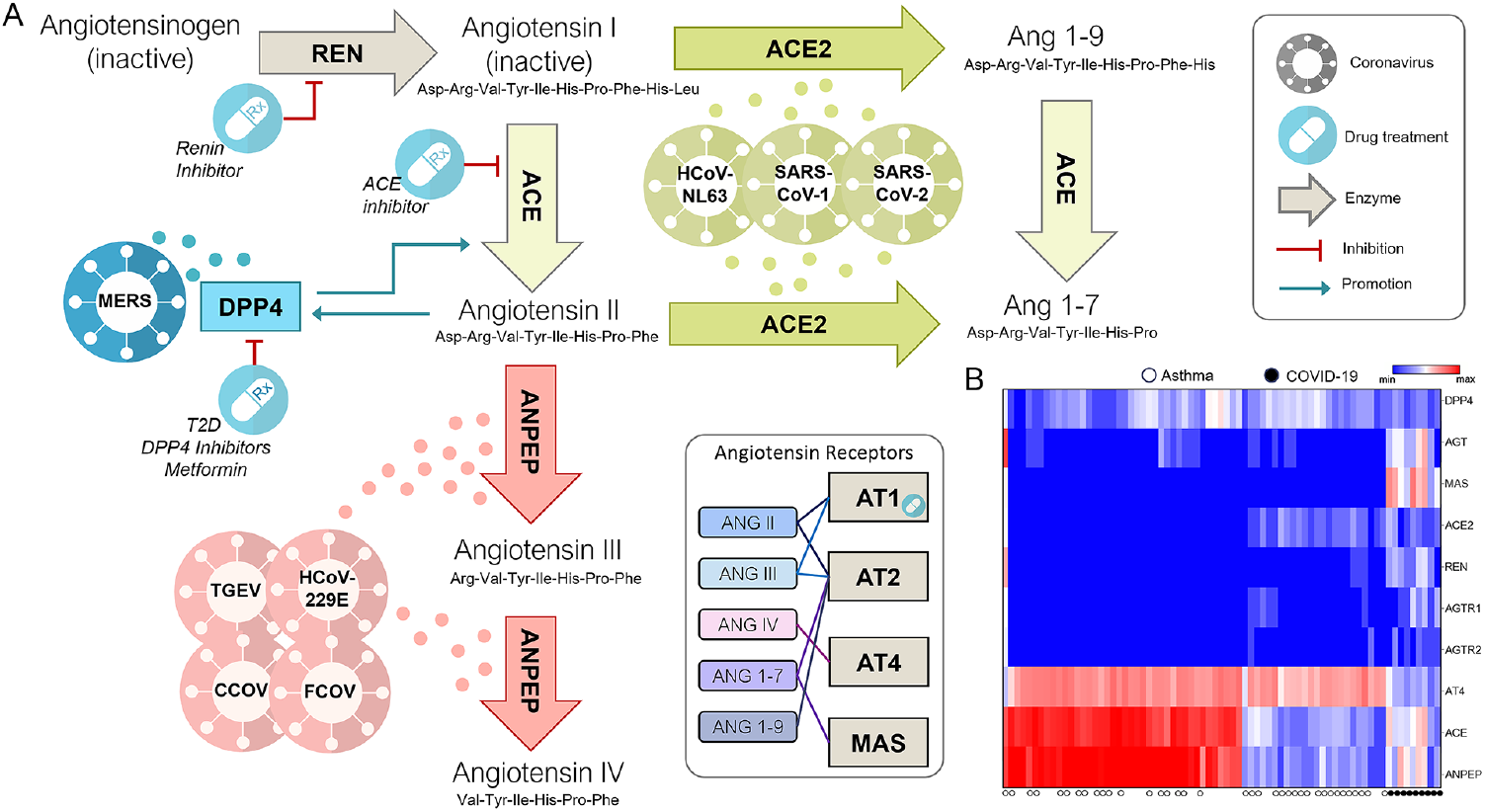
Brief summary of the Renin Angiotensin System (RAS). A) Angiotensinogen (AGT) is converted to angiotensin I by renin (REN), then to angiotensin II by Angiotensin Converting Enzyme (ACE), which is further cleaved to Angiotensin III and IV by Alanyl Amino Peptidase N (ANPEP). ACE2 converts angiotensin I and II to smaller peptides, angiotensin 1-9 or angiotensin 1-7. Four receptors are widely reported, namely, AT1, AT2, AT4, and MAS and different ligand/receptor pairs have different effects. Generally speaking, Ang 1-7/ACE2/MAS is an anti-inflammatory axis that counters ACE/AngII/AT_1_ by inhibiting NF-kappaB (*6*). Drugs used to treat hypertension and diabetes act on RAS, such as ACE inhibitors (e.g. captopril), RAS Inhibitors, and angiotensin receptor blockers. Inhibitors of the MERS receptor DPP4 can block ACE (*7*) and it can be regulated by angiotensin II (*8*). Several coronaviruses have evolved to use RAS proteins to enter cells (color coded according to known entry point) but mortality rates vary widely (MERS, 35%; SARS, 10%; COVID-19, 4%; HCoVNL63, rare). B) Cells from bronchoalveolar lavage samples of COVID-19 infected individuals show increased expression of REN, AGT, and MAS and downregulation of ANPEP and AT4 (heatmap lower right). Dehydration due to diarrhea, vomiting and perspiration could further alter RAS and affect virus pathogenicity as could several classes of widely prescribed drugs. Common side effects of these drugs are diarrhea and dry cough, similar to COVID-19. Deletion of an *Alu* sequence in ACE is correlated with higher blood levels of the enzyme and therefore this allele may be an informative predictor for outcomes of COVID-19. Focus on the entire RAS system is warranted in the continuing pandemic. Abbreviations: MERS (Middle East Respiratory Syndrome), TGEV (Transmissible Gastroenteritis Virus), FCoV (Feline Coronavirus), CCoV (Canine Coronavirus), HCoV229E (Human Coronavirus 229E), HCoV-NL63 (Human Coronavirus NL63), SARS (Severe Acute Respiratory Syndrome), and SARS-CoV-2 (cause of current COVID-19 pandemic).

### Is SARS-CoV-2 entering cells via ACE2 in the lungs?

A thorough analysis of the data from the Genotype-Tissue Expression (GTEx) Portal (https://www.gtexportal.org), the Proteomics DataBase (Proteomics DB) (https://www.proteomicsdb.org/) and the Human Cell Landscape (*9*) indicates ACE2 is either expressed at very low levels or is not detectable in tissues that are currently thought to be major entry points for SARS-CoV-1 and SARS-CoV-2 such as lung (Fig. 1-2, 4, Fig. S1, Supplementary Text). However, ACE2 is highly expressed in digestive tissue (gut, colon, rectum, ileum), reproductive tissue (testis, ovary), kidney, thyroid, adipose, breast and heart, possibly providing different viable routes for viral infection. Indeed, SARS-CoV-2 has been detected in feces in some individuals (*10*) and gastrointestinal disorders are now being reported as an early sign of COVID-19 infection (*11*). Additionally, the intestine has been shown to be a viable route for infection for MERS-CoV (*12*), suggesting that it survives the low pH of the stomach and is potentially replicating in digestive tissue.

Here, we revisit the hypothesis that infection occurs via the respiratory tract and posit that ACE may be the receptor for SARS-CoV-1 (and possibly SARS-CoV-2) in the lungs, given that the median mRNA levels of ACE are 39-fold higher than ACE2 which is often less than 1 TPM in lung tissue (Fig. 2 and Fig. S1). Similarly, the ACE protein is highly expressed in lungs, tonsils, salivary glands, and oral epithelium, whereas ACE2 is not detected in these tissues and yet, most citations identify ACE2 as the only receptor for SARS-CoV-1, often stating that it is highly expressed in the lung, when GTEx expression data, our analysis of the Human Cell Landscape and a careful review of the literature reveals that it is not. Furthermore, there is experimental evidence that ACE can mediate SARS-CoV-1 infection (Supplementary Text).

**Fig. 2.**
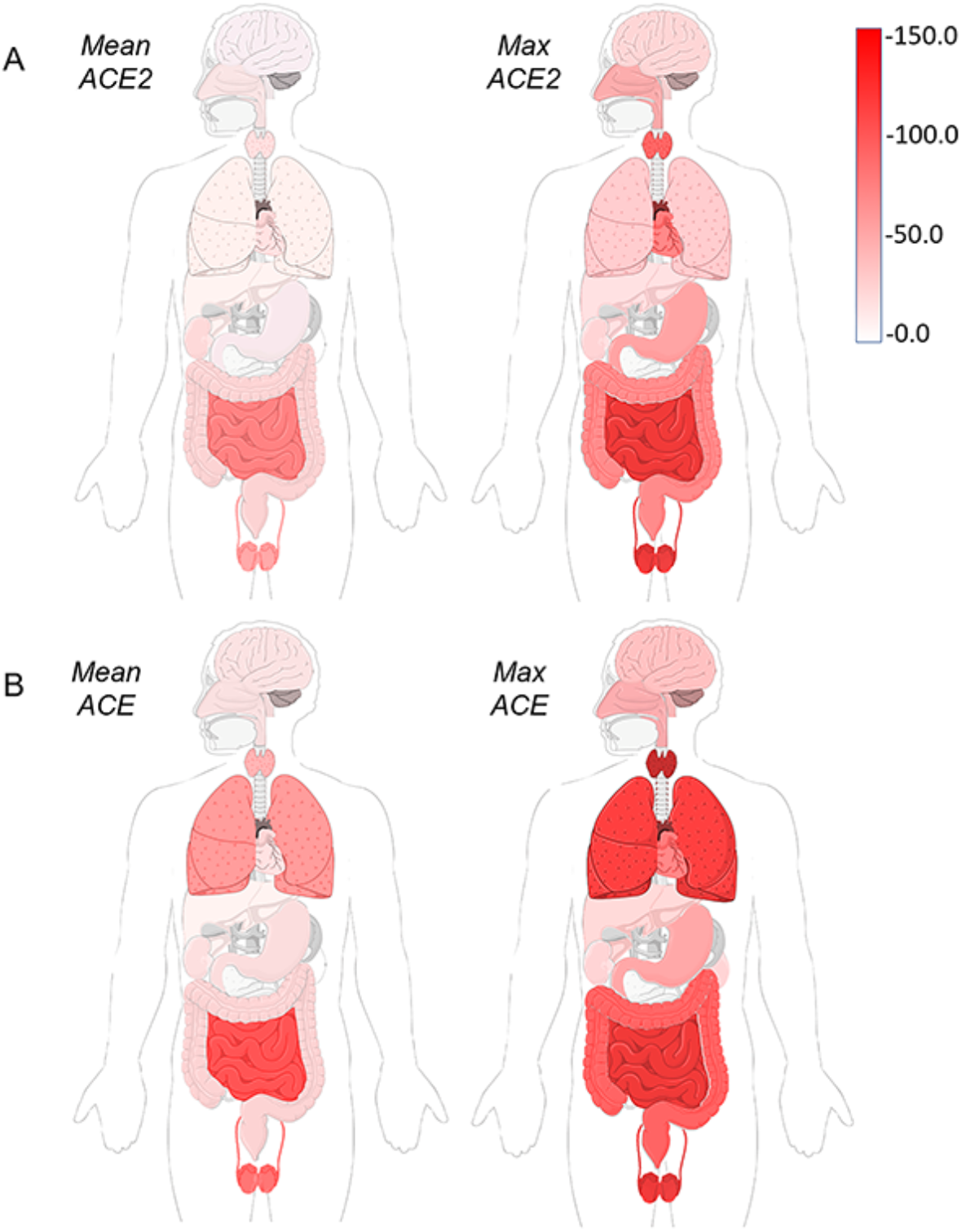
Mean and maximum expression levels (TPM) of ACE2 and ACE in different organs. Both ACE2 (A) and ACE (B) are highly expressed in digestive tissue and testis. Only ACE is highly expressed in the lung. Both ACE and ACE2 expression levels vary across individuals in the GTEx population.

**Fig. 3.**
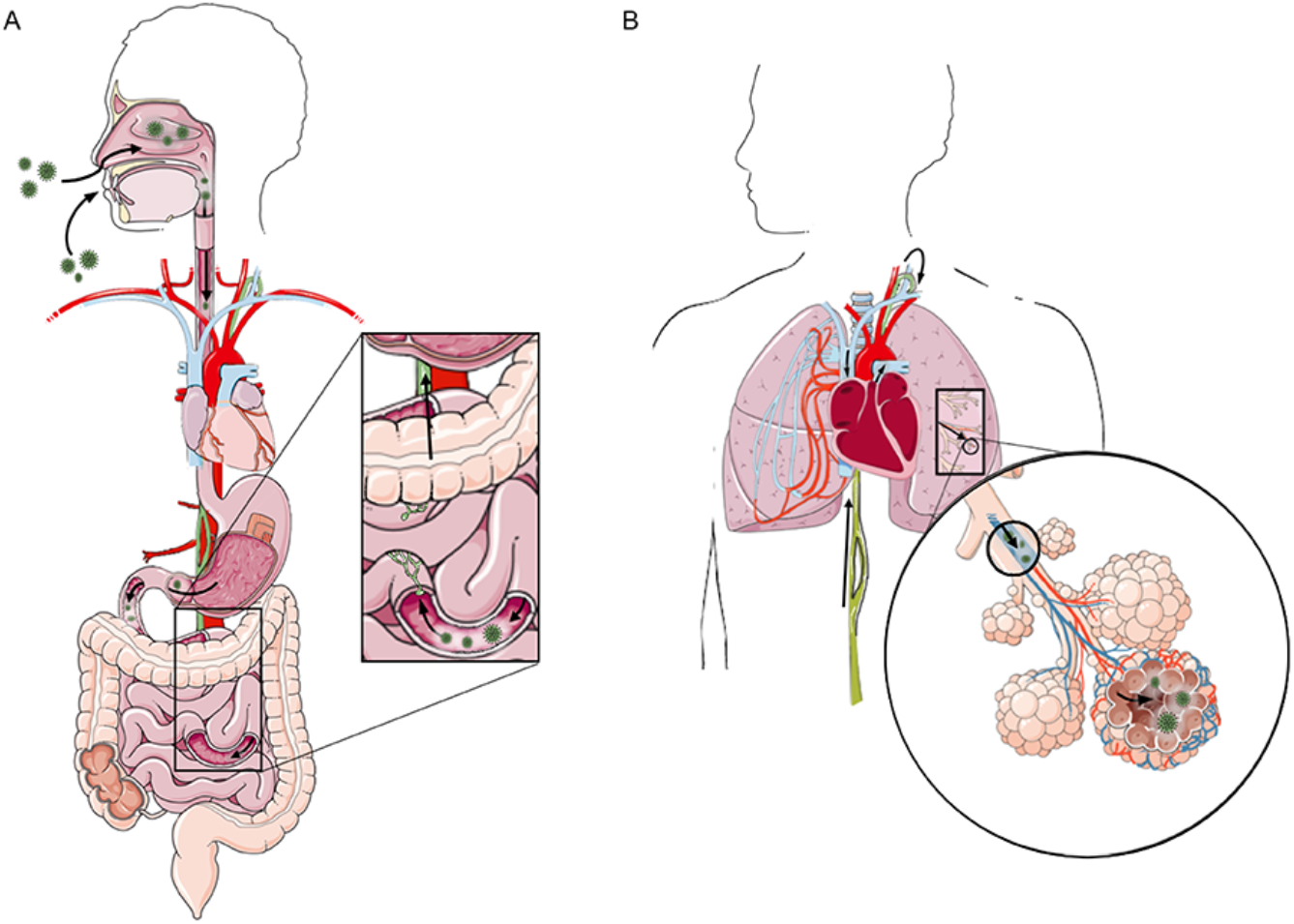
Possible routes of infection. Oral, nasal and throat tissues have low to moderate ACE2 expression that could support mild viral colonization. Intestinal tissue has high levels of ACE and ACE2 and can therefore become a major reservoir of SARS-CoV-2 virus, that can then migrate through the lymphatic system up to the thoracic duct, followed by entry into the venous system at the venous angle (Pirgoff’s angle) between the left subclavian vein and left internal jugular vein (A). From there, it can pass through the heart to reach the lungs, contacting the large surface area of the lung microvasculature, becoming a secondary site of infection. Respiratory infection may be facilitated in individuals with unusually high levels of ACE2 expressed in the lung, possibly due to comorbidity (B). From there the virus can spread through the circulatory system to other tissues with high ACE/ACE2 levels that correspond to symptoms being reported in COVID-19, *e.g.* testes (*13*), thyroid (fatigue (*14*,*15*)), brain epithelial cells (headache (*14*)), etc. This path of infection to the lung is supported by both bulk tissue and single cell gene expression patterns.

### What does cell-specific data tell us about ACE and ACE2 distribution?

We found considerable support of these differences in tissue expression of ACE vs. ACE2 in scRNA-Seq datasets (*e.g.* The Human Cell Landscape (*9*) and The Mouse Cell Atlas (*16*), Fig. S2). A single-cell atlas of human respiratory tissues available on the ToppCell server (toppcell.cchmc.org) reveals that, in contrast to high levels of expression of ACE in lung endothelial cells, ACE2 is very sparsely expressed in several different lung cell types (Fig. 4). Single cell data from the mouse atlas revealed that ACE expression is high in lung endothelial and mast cells, trachea, pancreatic endothelial and stellate cells, and heart fibroblasts (Fig. S2). High ACE2 expression occurred in the large intestine epithelium, pancreatic endocrine cells, heart myofibroblasts and the tongue keratinocytes (Fig. 4). Taken together, these data indicate that ACE2 is more sparsely expressed than ACE and is not detected in the endothelial or fibroblast lineages that express ACE (Fig. 4). However, moderate levels of ACE2 were found in differentiating keratinocytes of both nasal passage and the tongue (Fig. 4), with higher levels in enterocytes of the small and large bowel, and several important endocrine cells of the pancreas. Finally, nasal epithelial brushings have been an important site for detection of SARS-CoV-2 and we therefore sought to determine if an ACE2-positive signal in nasal brushings could indicate the presence of virus-susceptible cell type(s). We analyzed expression patterns of 937 genes that are highly correlated/anti-correlated with ACE2 in lung RNAseq data from GTEx and from RNA measured in nasal epithelium cells (from nasal brushings) in individuals of different ages. Although ACE2 is not highly expressed, there appear to be sub clusters of individuals defined by differentially expressed genes that participate in the development of the trigeminal nerve and microvilli in teens and adults. Infants do not express ACE2 to any appreciable extent (Fig. 5).

**Fig. 4.**
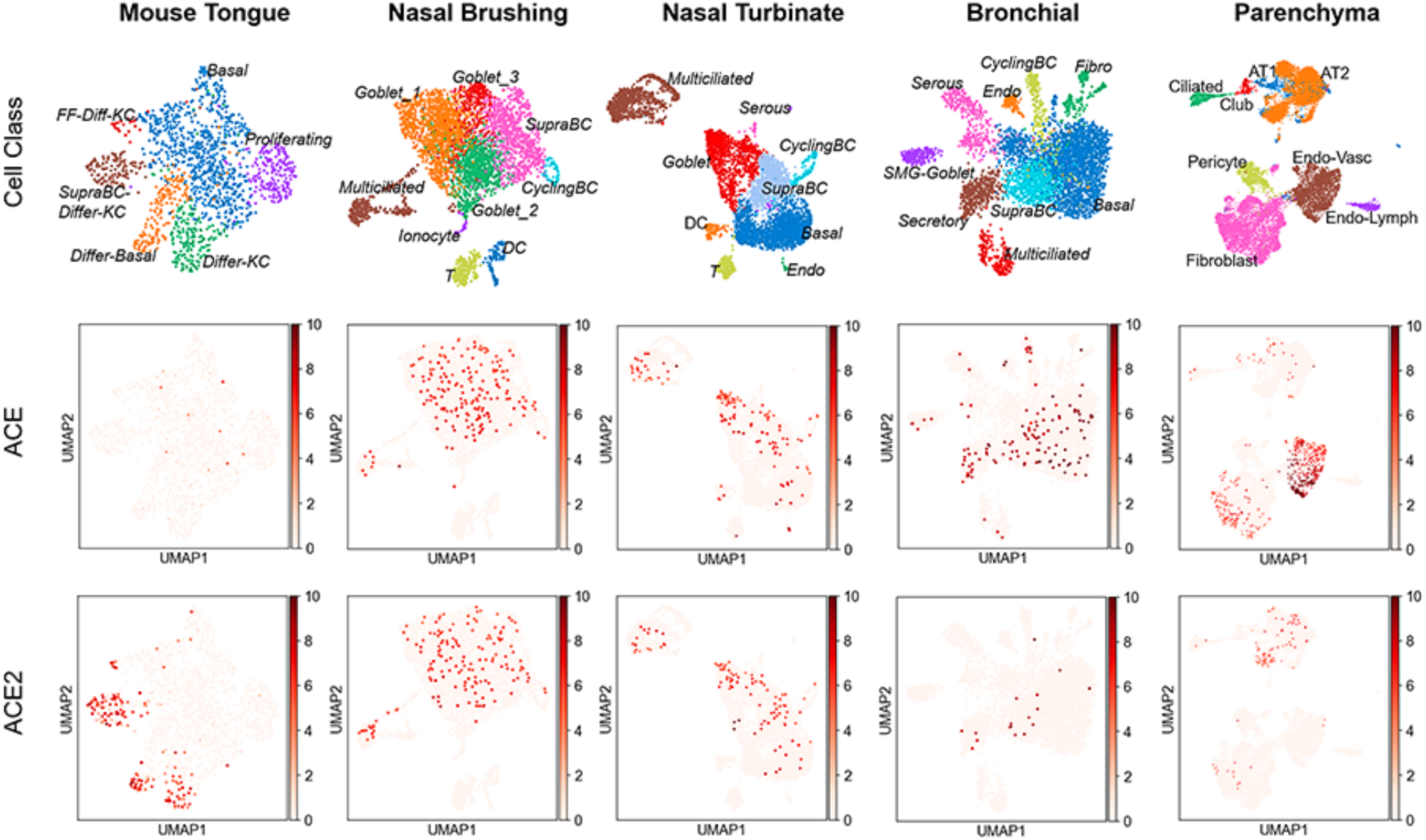
A variety of human single cell datasets from nose (*17*) to lower lung (*18*) were used to explore gene expression in previously identified cell lineages. The most consistent site of ACE2 expression was in differentiating keratinocytes of nasal brushing and nasal turbinates. Similarly, mouse ACE2 was most consistently expressed in differentiating keratinocytes of the tongue. Color scales are Log2(TPM+1). FF-Diff-KC, filiform differentiated keratinocytes; SupraBC-Differ-KC, suprabasal differentiating keratinocytes; Differ-Basal, differentiating basal cells; Differ-KC, differentiated keratinocytes; SupraBC, suprabasal cells; cyclingBC, cycling basal cells; T, T cells; DC, dendritic cells; Endo, endothelial cells; SMG-Goblet, submucosal gland Goblet cells; Fibro, Fibroblasts; AT1, Alveolar Type 1 cells; AT2, Alveolar Type 2 cells; Endo-Vasc, vascular endothelial cells; Endo_Lymph, lymph vessel endothelial cells.

**Fig. 5.**
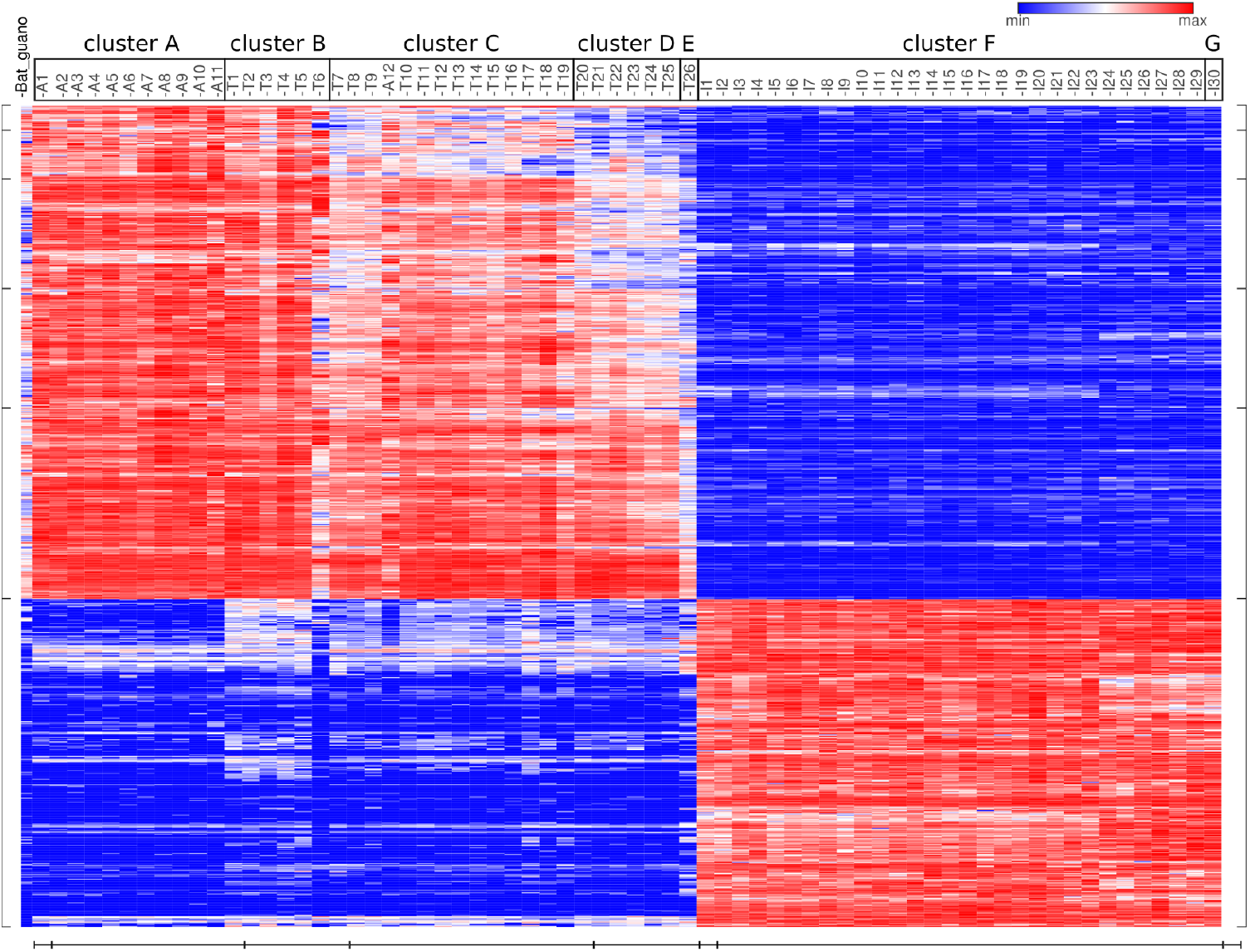
Clustering of 937 differentially (vertical axis) expressed genes that were also highly correlated with ACE2 in GTEx from lung (+/−0.8) in nasal brushings in different age classes (horizontal axis). ACE2 was not detected in the majority of infant (I) nasal epithelium and was moderately expressed in adults (A) and teens (T) (range 1.0-5.2 TPM). Hierarchical clustering identified sub-patterns of expression within those expressing ACE2 (e.g. cluster C is specific to one adult and a small number of younger individuals). A GO analysis of the genes upregulated with ACE2 (top left quadrant) identified a 30-fold enrichment for trigeminal ganglion development and 26-fold enrichment for microvilli assembly Those that were down-regulated compared to ACE2 (*blue*, lower left quadrant) did not show significant GO enrichment but were mainly non-coding RNAs. A sample from a bat guano collector in Thailand infected with a HKU1 coronavirus appears on the far left.

### ACE vs. ACE2 as the entry point for SARS-CoV-1 and SARS-CoV-2

Closer scrutiny of the initial report identifying ACE2 as the receptor for SARS-CoV-1 suggests it may be specific to kidney-derived cell lines and the potential for ACE as the receptor has not been adequately evaluated (Supplementary Text). Applying computational methods to study the molecular biophysics of viral and host machinery has been an effective way of checking this hypothesis and identifying key molecular elements of interactions with these putative receptors that can then be further explored in depth.

Currently, high-resolution structures of the receptor binding domains (RBD) of the SARS-CoV-1 and SARS-CoV-2 spike glycoproteins in complex with the peptidase domain of ACE2 are available, and enable a detailed description of the interfacial interactions (*19*). In order to explore the involvement of ACE and ACE2 in SARS-CoV-1 and -CoV-2 infection, we used these structures as a starting point for a comparative atomistic molecular dynamics (MD) study of the RBDs of the two viruses in complex with ACE2, here referred as RBD1-ACE2 and RBD2-ACE2 for SARS-CoV-1 and SARS-CoV-2, respectively. Additional simulations of the putative complexes with ACE were performed for the preliminary study of the stability of these complexes (RBD1-ACE and RBD2-ACE).

The amino acid sequences of SARS-CoV-1 and SARS-CoV-2 spike glycoproteins are 77% identical. The RBD is a region that harbors a high concentration of non-conservative substitutions, remarkably at regions that are known to directly bind to the host receptor (Fig. S3). These differences are expected to considerably influence the affinity of spike for ACE2. Tian *et al*. measured the binding of RBD2 to ACE2 with a biolayer interferometry binding assay and reported similar affinity of RBD1 and RBD2 to ACE2 (*K*_*d*_=15·0 nM and 15·2 nM, respectively) (*20*). In contrast, Wrapp *et al*. reported a 10- to 20-fold higher affinity of RBD2 to ACE2, compared to RBD1 (*21*). Despite the disagreement on the relative affinity for ACE2, both reports establish that RBD2 forms a stable complex with ACE2, which was shown to be a host receptor for SARS-CoV-2 cell entry (*22*).

Accordingly, RBD1-ACE2 and RBD2-ACE2 complexes are stable along all of the conducted MD simulations. The computed average number of contacts (residues with Cα less than 8 Å distant) is slightly smaller between RBD2 and ACE2 (22 ± 4) than in the complex with RBD1 (25 ± 3). This suggests that, if RBD2 has higher affinity to ACE2 than RBD1, as reported by Wrapp *et al*., this is resulting from stronger rather than additional interactions in RBD2-ACE2. As shown in Fig. 6A, the profile of ACE2 residues involved in persistent interactions with the RBDs is consistent in triplicate simulations. The contact profile, Fig. 6A-B, shows a slightly higher density of stable contacts in zone 2 for RBD1-ACE2 compared to RBD2-ACE2, that is likely due to the additional salt bridge formed by RBD1 Arg^426^ and ACE2 Glu^329^, which is lost with the substitution Arg^438^Asn in RBD2, as well as due to the presence of RBD1 Tyr^484^ (Gln^498^ in RBD2), packing with the hydrophobic tail of ACE2 Lys^353^. The weaker interactions in zone 2 are at least partially compensated in RBD2-ACE2 in zone 1, where hydrophobic packing is enhanced by the bulky RBD2 Phe^486^ and Phe^456^ (Leu^472^ and Leu^443^ in RBD1, respectively).

**Fig. 6.**
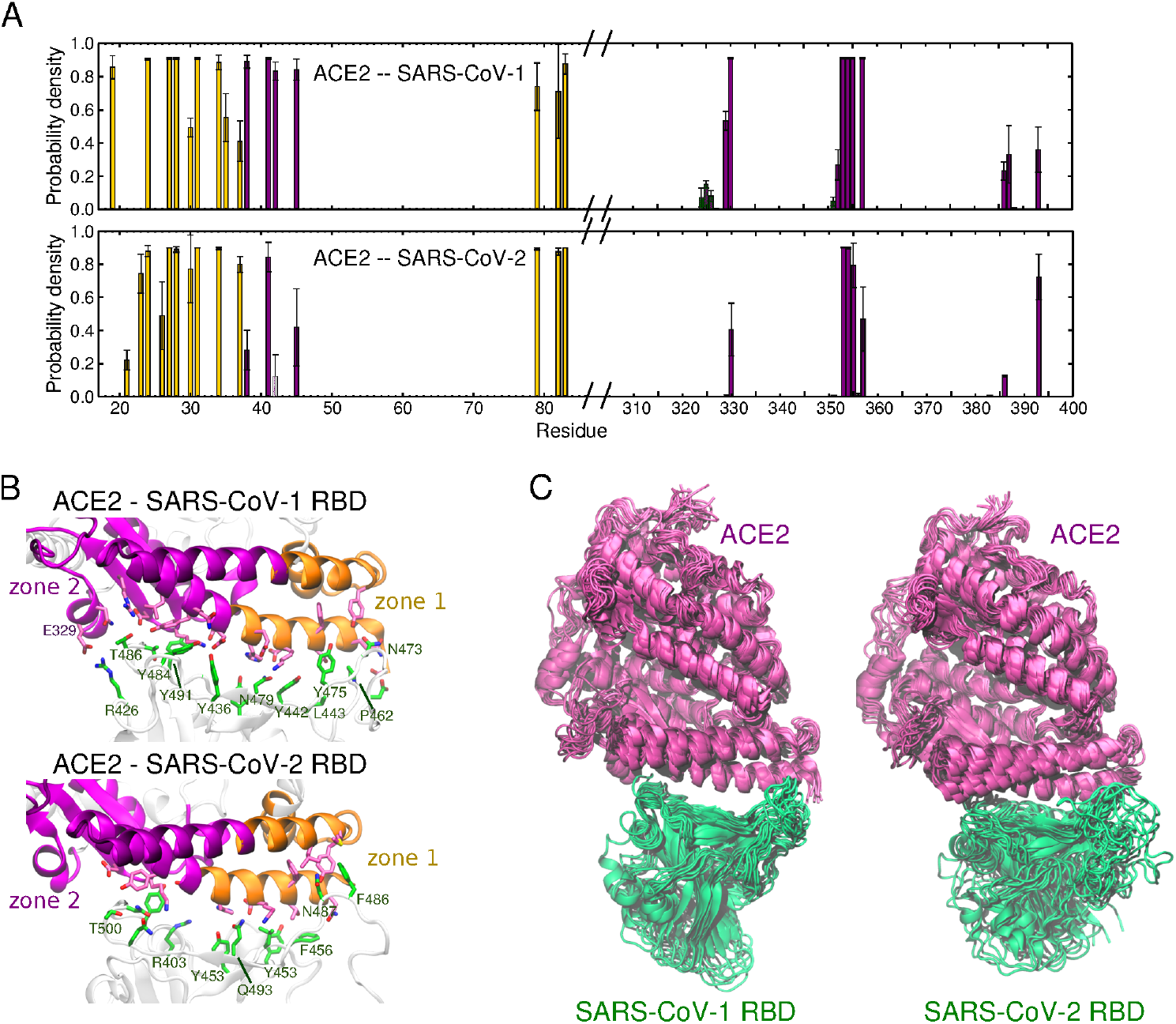
Analysis of simulations of SARS-CoV-1 and SARS-CoV-2 RBDs in complex with ACE2. A) Probability density of residues from ACE2 forming contacts with the RBDs. A maximum distance of 4 Å between any atom in a pair of residues was established. Bars with a standard deviation greater than 50% of the probability density are considered transient contacts in the simulations and not included in these plots. The colors of the bars correspond to zone 1 and zone 2 of ACE2, defined in (B), which shows the residues involved in contacts formed during more than 70% of the simulation time. RBD and ACE2 residues are represented as licorices, in *green* and *pink*, respectively. C) Superimposition of frames in a representative simulation of RBD1-ACE2 (*left*) and RBD2-ACE2 (*right*), using the initial position of ACE2 as reference for alignment. RBDs and ACE2 are represented in *green* and *pink*, respectively.

The analysis of the conformational dynamics of the two complexes revealed an important structural difference that was not captured in previous analyses of static structures. The RBD from SARS-CoV-2 exhibits considerably higher flexibility compared to the RBD from SARS-CoV-1, as evident in the superimposition of the simulation frames (Fig. 6C). A close inspection of the structure strongly suggests that this is caused by the substitution of Lys^447^ by Asn^460^ in RBD2, resulting in the loss of a salt bridge with Asp^407^, or Asp^420^ in RBD2 (Fig. S5). The weaker interaction with the α3 helix “unlocks” loop β4-5, that mostly interacts with zone 1 of ACE2. The elongation of the loop with the additional glycine, Gly^482^, further contributes to the higher conformational flexibility of the SARS-CoV-2 RBD. The effectiveness of drugs targeting the S-ACE2 interface will likely be correlated with the differences in this local mobility.

In order to perform preliminary tests of the hypothesis that ACE is an alternative receptor for SARS-CoV-1 and SARS-CoV-2, we also conducted MD simulations of RBD1-ACE and RBD2-ACE. The peptidase domains of ACE and ACE2 are 40% identical and have a very similar fold (RMSD 6.6 Å) and therefore we assumed that the interaction with RBDs would occur in the same region in the fold. We built the initial structures by alignment and replacement of ACE2 by ACE in the complexes described above. On the putative complex interface, only 35% of the residues are similar or identical to residues identified as stable in the interaction of ACE2 with RBD1 or RBD2. Therefore, local structural rearrangements are expected to happen in the built RBD1/2-ACE complexes along the MD simulations. In order to allow structural adjustments to happen, we conducted long equilibration simulations involving multiple steps for gradual relaxation of the system.

Our analysis verifies that, for both systems, the RBDs remain bound to ACE during the simulations. In all independent simulations of RBD1-ACE, a significant reorientation of the RBD1 is observed, so that the loop ß5-6 slides towards the center of the α1 helix of ACE. Fig. 7A shows the superimposed last frames of the three simulations of RBD1-ACE. Persistent interactions are established involving the formation of three salt bridges, namely, Asp^407^-Arg^53^, Lys^447^-Glu^49^, Asp^493^-Lys^94^, from RBD1 and ACE, respectively (Fig. 7B).

**Fig. 7.**
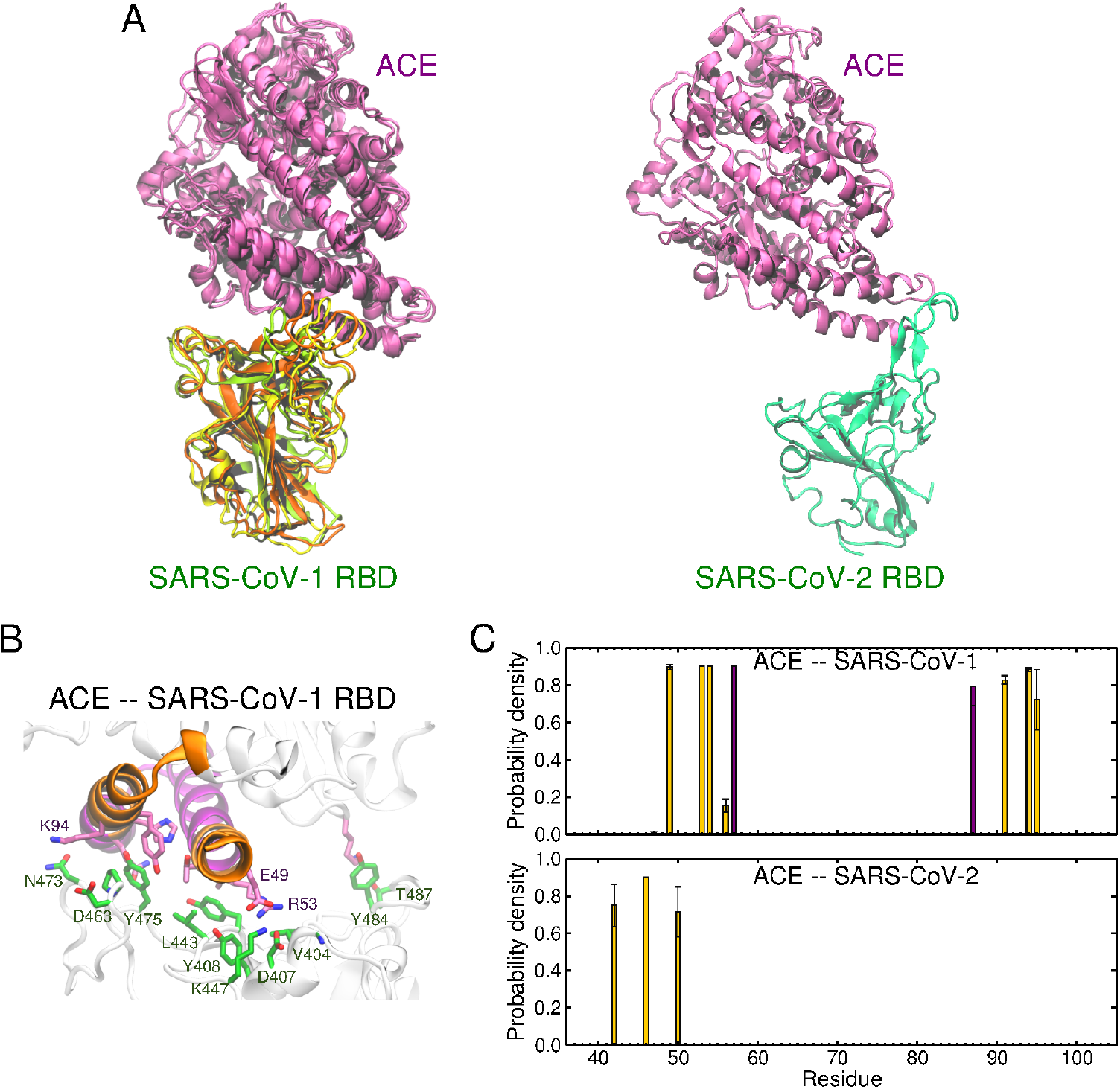
Analysis of simulations of SARS-CoV-1 and SARS-CoV-2 RBDs in complex with ACE. A) Superimposition of the last frames of the simulations of RBD1-ACE (*left*). For visual clarity, because the relative orientation of the proteins in RBD2-ACE is very flexible due to the small surface of contact, we only show the last frame of a representative simulation of RBD-ACE (*right*). RBDs and ACE are represented in *green* and *pink*, respectively. B) Residues involved in contacts formed during more than 70% of the simulation time. RBD and ACE residues are represented as licorices, in *green* and *pink*, respectively. C) Probability density of residues in ACE forming contacts with the RBDs. A maximum distance of 4 Å between any atom in a pair of residues was established. Bars with standard deviation higher than 50% of the probability density are considered transient contacts in the simulations and not included in these plots. The colors of the bars correspond to zone 1 and zone 2 of ACE, shown in B.

The RBD2-ACE also converges to a common configuration in two of the three independent simulations of RBD2-ACE, with only few residues attaching the proteins together (Fig. 7A). In these simulations, the loop ß4-5 anchors the RBD2 to the N terminal of α1 helix and the nearby region of α2 helix of ACE, mostly involving only hydrophobic contacts between Phe^456^ and Tyr^489^ of RBD2 at the N-terminal of α1 (Fig. 7C).

Despite the fact that MD simulations of hundreds of nanoseconds cannot provide reliable quantitative estimates of binding affinity, they can be effectively used to explore the relative stability of the studied complexes. Taken together, our simulations demonstrate the convergence of stable and strong interactions between ACE and SARS-CoV-1, suggesting that ACE may allow for infection in tissues with low or undetectable levels of ACE2, but high ACE expression. This is supported by previous work showing ACE’s ability to increase SARS-CoV-1 infection in some cell types (Supplementary Text). Our results emphasize that future experiments should be designed to include the complete native spike protein, since intra-spike interactions of the RBD in the closed conformation are an important element competing with the stabilization of the open conformation of the spike via interaction with the host receptor. Additionally, studies on SARS-CoV-1 and future zoonotic coronaviruses should include ACE in their analyses.

### The forgotten co-receptors of SARS-CoV-1 infection - other co-receptors that may be involved

In the study that originally identified ACE2 as the receptor for SARS-CoV-1, two additional proteins were found to bind the spike glycoprotein - major vault protein (MVP) and myosin 1B (MYO1b) - but they were not further explored at that time (*23*). More recent data has indicated that these are, in fact, important elements for viral infection. MYO1b is directly linked to SARS because it is necessary for internalization of feline coronavirus (which also enters cells via the RAS) from the cell surface (*24*). MVP is the major component of Vault ribonucleoprotein particles, which are known to be involved in multidrug resistance in cancer cells (*25*, *26*) and are a central component of the rapid innate immune response in cells exposed to pathogens (*27*).

As part of the immune response to lung infection by the bacterium *Pseudomonas aeruginosa,* MVP is recruited to lipid rafts that mediate internalization of the pathogen via endocytosis, and it is likely essential to initiate an apoptotic cascade (*28*). These lipid rafts include 127 other proteins within them (table S2, henceforth referred to as “raft-127 genes”), including angiotensin II receptor-associated protein (AGTRAP), whose function is, in part, to internalize AT1-type receptors via endocytosis (*29*) and cycling to the endoplasmic reticulum and Golgi apparatus (*30*). In addition, several antiviral response genes are present including MAVS, CD81, ITGA2, NECTIN1, TFRC, and LDRL. NECTIN1 is a receptor for many different viruses including herpes, measles, and pseudorabies virus, and LDLR is a known receptor for vesicular stomatitis virus (*31*) and facilitates production of hepatitis C virus (*32*) Given the link between MVP and SARS-CoV-1 that was previously reported (*23*), these findings indicate that other proteins are important for SARS-CoV-1 and SARS-CoV-2 entry into cells and their replication, likely by hijacking the lipid raft-mediated endocytosis that is part of a normal immune response.

Indeed, the importance of the raft-127 genes was revealed in a recent study that used affinity-purification mass spectrometry to identify host proteins that bind to those expressed by SARS-CoV-2; of the 332 proteins identified using a high-stringency filter, seven are found in the raft-127 set and four of those seven bind to nsp7, which is part of SARS-CoV-2 replication complex. Furthermore, these four proteins (RAB5C, RAB18, RHO and RALA) are all involved in the same process, endocytosis/vesicle trafficking. A relaxed filter identified another 33 raft-127 genes including Cathepsin D (CTSD), which is substantially downregulated in cells from bronchoalveolar lavage samples taken from COVID-19 patients, and here binds to the SARS-CoV-2 ORF8 protein. This protein harbors the mutation that defines the “S” and “L” lineages of the current pandemic (*33*) that have been associated with differing pathogenic behavior (Fig. S6) (*34*).

### Gene expression patterns of SARS-CoV-2 infected bronchoalveolar lavage cells

In order to gain a broader understanding of SARS-CoV-2 infection, we carried out a transcriptome-wide analysis of these samples and identified 2,057 significantly downregulated and 591 upregulated genes. The downregulated genes were highly enriched for innate and adaptive immune response and endosomal trafficking; the upregulated genes were highly enriched for non-coding RNA (https://compsysbio.ornl.gov/sars-cov-2-mechanisms-of-infection/). Notably, 57 of the 127-raft genes were significantly differentially expressed in the BAL compared to control (55 downregulated and two upregulated).

We then filtered these differentially expressed genes for those that were highly correlated or anti-correlated with ACE2 (Fig. 8) and found that ACE2-correlated upregulated genes in COVID-19 BAL samples were highly enriched for gene signatures of lung epithelial cell types including type 1 and type 2 alveolar epithelial cells, Club Cells, and ciliated cells. Thus, for at least 5 of the 9 samples, those parenchymal cells are present in an intact state in the BAL fluid. Remarkably, lymphatic endothelial cells are also present, whereas expected gene signatures for vascular endothelial cells were not. In addition to the signatures of those cell types, there is significant activation of additional genes in key functional categories for those cell types including adhesion, tight junctions, and epithelial tubular morphogenesis. This could suggest that the epithelial cells are attempting to respond to virus-driven tissue dissolution by upregulation of adhesion and tube forming genes. The sum of these cell signatures and extended categories accounts for more than 500 of the 1100 gene signatures that were observed. The upregulated genes also lacked indication of any cytokine storm or cytokine-activated status.

**Fig. 8.**
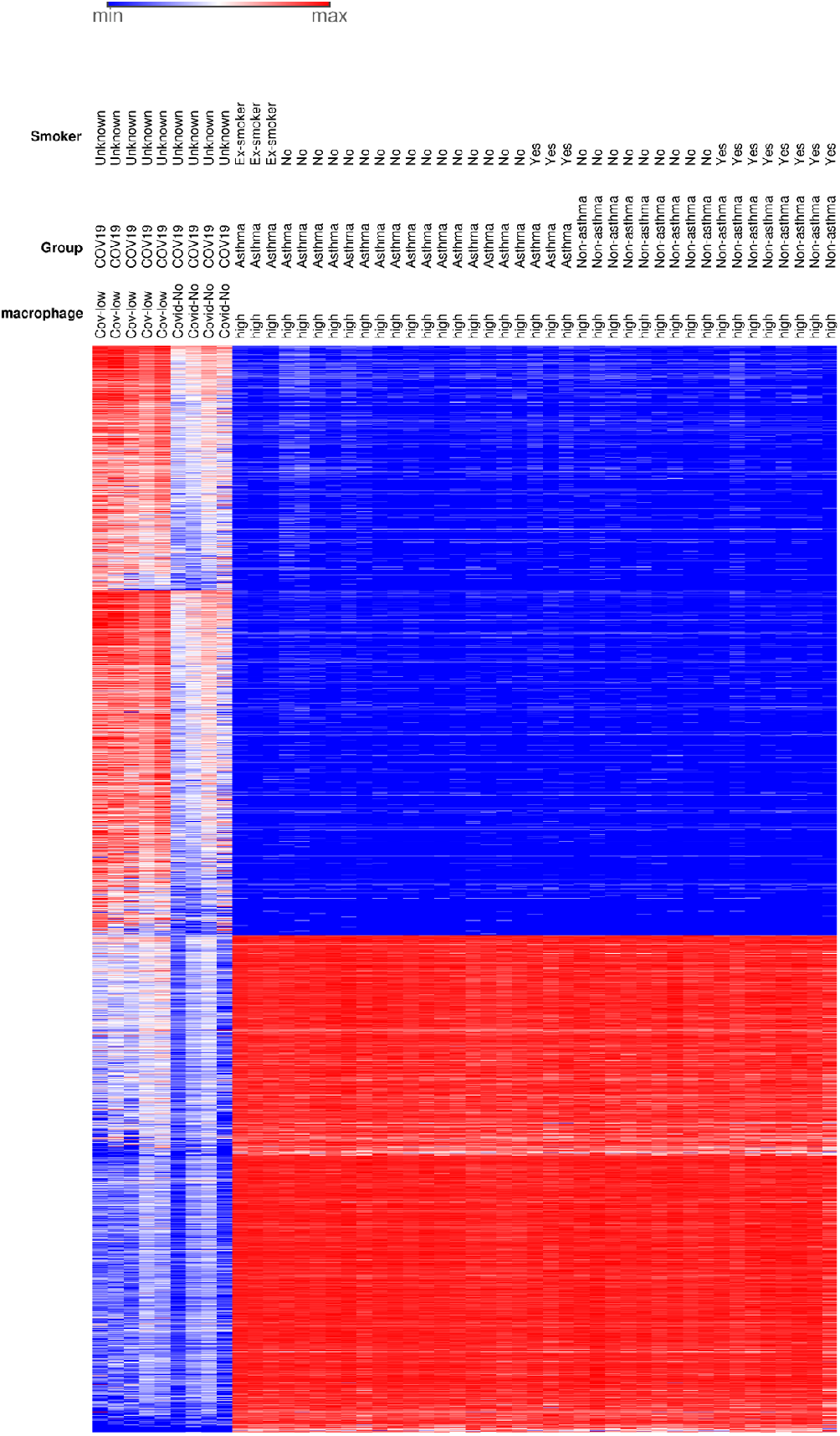
Overview of globally differentially expressed genes in COVID-19 BAL samples relative to controls. The COVID-19-overexpressed clusters are highly enriched for genes that define lung structural cell types including epithelial type I, type II, and lymphatic endothelial cells (upper right quadrant). There is also increased expression for genes related to intercellular or extracellular adhesion compared to controls, as well as additional genes in those categories not normally expressed in those cell types. The ACE2-correlated differentially expressed genes in the lower half of the heatmap define macrophage and dendritic cells, which are uniformly absent in the COVID-19 samples (lower right quadrant). None of the COVID-19 samples matched the pattern in the controls, indicating exclusion of these cells from BAL fluid in infected cells. The COVID-19-repressed signature (964 genes, https://compsysbio.ornl.gov/sars-cov-2-mechanisms-of-infection/) is a complete inventory of genes associated with macrophages and dendritic cells, suggesting that COVID-19 actively destroys these cells or reprograms them to the point of being unrecognizable.

The ACE2 anti-correlated, downregulated gene signature in the COVID-19 BAL samples is particularly noteworthy for the absence of a rich suite of genes broadly associated with multiple myeloid cell types including macrophages, dendritic cells and neutrophils, particularly CSFR1+ macrophages. As mentioned above we found that several components of RAS are upregulated in bronchoalveolar lavage (BAL) samples taken from COVID-19 patients, including MAS, which appears to be a marker for the disease (Fig. 1), and which is associated with decreased cytokine release and migration of immune cells (*6*). These results are consistent with the release of a myriad of lung parenchymal cells into the fluid phase as a result of SARS-CoV-2 infection from a distant location (Fig. 2). The absence of macrophages and the induction of a RAS-mediated cytokine suppression pathway via MAS in COVID-19 lung samples suggest that a major component of SARS-CoV-2’s virulence is its apparent ability to cause a functional immune deficiency syndrome in lung tissue.

### Structural models are a valuable resource for Systems Biology approaches

Structural analyses and molecular dynamics studies of SARS-CoV-1 and SARS-CoV-2 spike proteins revealed differences that significantly affected the interaction with putative host receptors. Next, because SARS-CoV-1 is the closest coronavirus evolutionarily to SARS-CoV-2 that can infect humans and that has been intensely studied, we extended the structural analysis to all mature proteins of SARS-CoV-2 to identify other critical elements that may enhance the spread of COVID-19 compared to SARS. We used the currently available experimentally solved structures of SARS-CoV-2 proteins, and an ensemble workflow to generate structural models and profiles of all unsolved viral proteins (Supplementary Materials).

The proteome of SARS-CoV-2 includes four structural proteins, namely, spike (S), envelope (E), membrane protein (M), and nucleocapsid (N). It also produces 15 mature non-structural proteins (nsp1-nsp10 and nsp12-nsp16), and nine accessory proteins (*35*) An in-depth comparative genome study reported that 380 amino acids that are fixed across thousands of SARS-like coronaviruses are changed, and specific to SARS-CoV-2 (*36*), suggesting that this minor portion may be essential for determining the pathogenic divergence of COVID-19. Here we verified that, in total, including the mutations that have occurred over the course of the pandemic to date, there are 1570 amino acid substitutions between SARS-CoV-2 and SARS-CoV-1 among these proteins. As shown in Fig. 9, the majority of variations are non-conservative and distributed among the mature proteins, but several nsps are highly similar to their counterparts in SARS-CoV-1. The strong conservation of these nsps would tend to suggest strong purifying selection to maintain viral fitness. This relationship and their likely long-term stability in the population makes them attractive drug targets. Except for ORF8, the most variable sequences diverge approximately 30% relative to SARS-CoV-1, which typically do not change global topologies and, consequently, the main protein function.

**Fig. 9.**
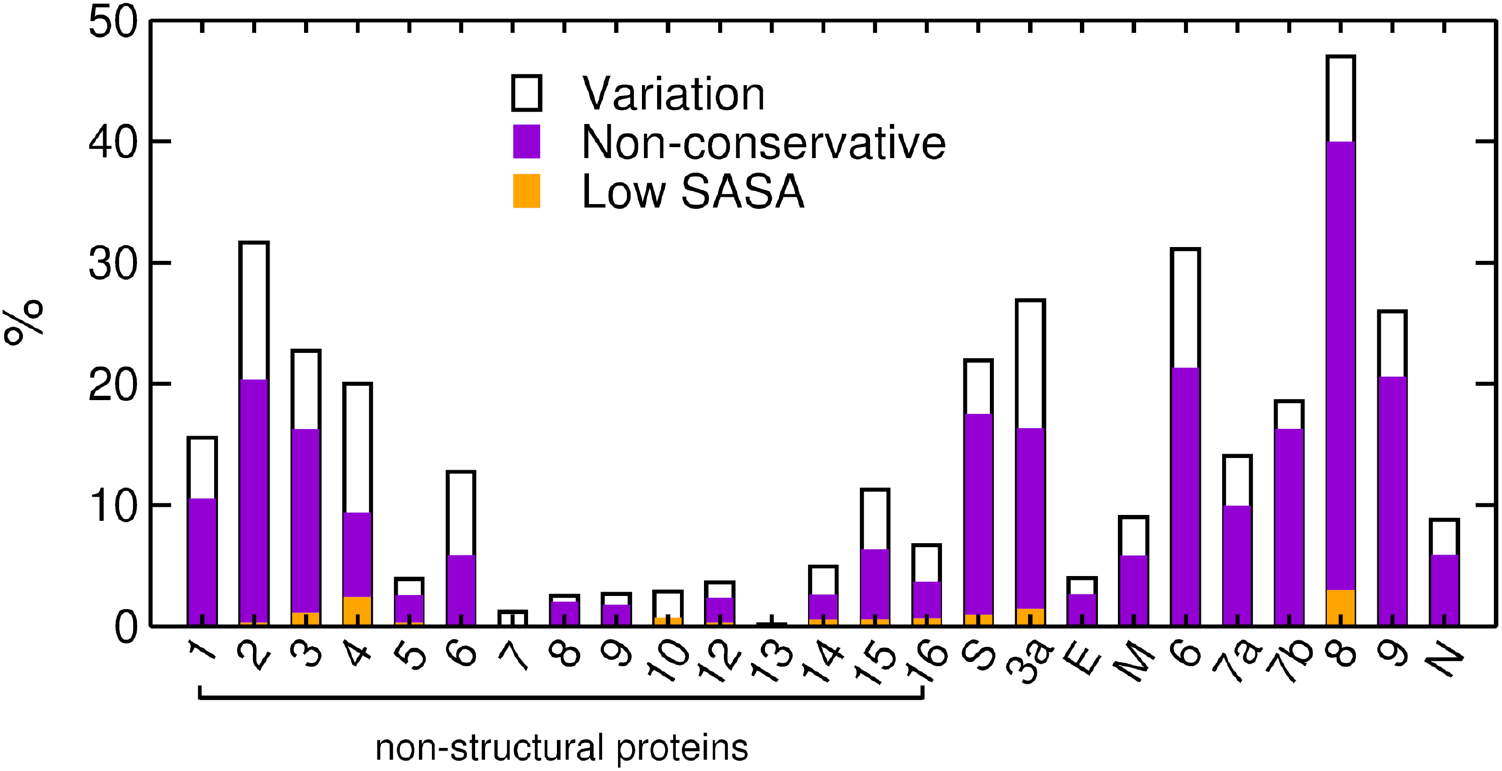
Distribution of sequence variation in SARS-CoV-2 proteins relative to SARS-CoV-1. Variations considered non-conservative, represented in *violet*, are defined in table S4. Variations occurring within protein cores (low solvent accessible surface area, SASA), are represented in *orange*. ORF10 is not included as it is not found in the SARS-CoV-1 proteome.

Additionally, visual inspection of non-conserved substitutions in the predicted structures combined with analyses of their structural profiles (i.e., predicted location of structured, intrinsically disordered and transmembrane regions - Appendix 2.5), indicates that the great majority of them are located in superficial regions of the proteins or protein domains. That is, most of the substitutions do not significantly affect protein folding, but they potentially can affect post-translational modification patterns and protein function if located in key regions for interactions with other proteins, ligands and substrates. Along with the predicted models, we provide to the scientific community a synopsis for 27 mature viral proteins and nsp11 (short peptide), in which we mapped the variation relative to SARS-CoV-1, the mutations of SARS-CoV-2 occurring globally, and analyzed their potential relevance concerning the pathobiology of COVID-19. Here, as examples, we discuss the possible effects of non-conservative variations in the nonstructural proteins, nsp1 and nsp5.

### Pathogenic-relevant substitutions in nsp1 and nsp5

The nsp1 protein is associated with the degradation and suppression of the translation of host mRNA. 3CL^pro^ (nsp5), also commonly referred to as the main viral protease, cleaves the polyproteins translated from the invading RNA(*37*–*39*) to form the mature viral proteins. We highlight molecular differences of these two proteins between SARS-CoV-1 and SARS-CoV-2 as they relate to host immune response and to pathogenicity divergence, being promising targets for drug development, drug repurposing, or vaccine production.

In addition to its role in processing the viral proteome, we propose that the highly conserved nsp5 protein may also be part of a major mechanism that suppresses the nuclear factor transcription factor kappa B (NF-κB) pathway, eliminating the host cell’s interferon-based antiviral response. In SARS-CoV-1, several proteins have been reported to be interferon antagonists, including nsp1 and nsp3 (*40*, *41*). and in COVID-19 BAL samples, the NF-κB inhibiting MAS protein is induced (Fig. 1). An additional mechanism of circumventing the interferon antiviral response is described for the porcine epidemic diarrhea virus (PEDV) as well as non-coronaviruses (*42*), in which the 3CL^pro^ cleaves the NF-kB essential modulator (NEMO) (*43*). Given that the substrate binding site of SARS-CoV-2 3CL^pro^ is very similar to PEDV 3CL^pro^, it is possible that SARS-CoV-2 3CL^pro^ is also active towards NEMO. Structural divergence is concentrated in the region corresponding to the S2 binding site of PEDV and in the peptide segment 45-51, in the catalytic entrance (Fig. 10A). As a preliminary test for this hypothesis, we conducted molecular docking of NEMO targeting SARS-CoV-2 and PEDV 3CL^pro^ proteins. The best resulting substrate conformation has the Gln^231^ reaction center of NEMO positioned very similarly to the PEDV 3CL^pro^-NEMO crystal structure. The estimated binding affinity is −6·2 kcal/mol for SARS-CoV-2 3CL^pro^-NEMO, and −7·4 kcal/mol for PEDV 3CL^pro^-NEMO. The binding site of SARS-CoV-2 3CL^pro^ is conserved relative to SARS-CoV-1 3CL^pro^, except by the substitution Ala^46^Ser in the entrance of the cleft, indicating that SARS-CoV-1 3CL^pro^ may also be active towards NEMO. This result suggests that drug development targeting this mechanism may prove fruitful as it would allow for a normal host immune response to combat the pathogen.

**Fig. 10.**
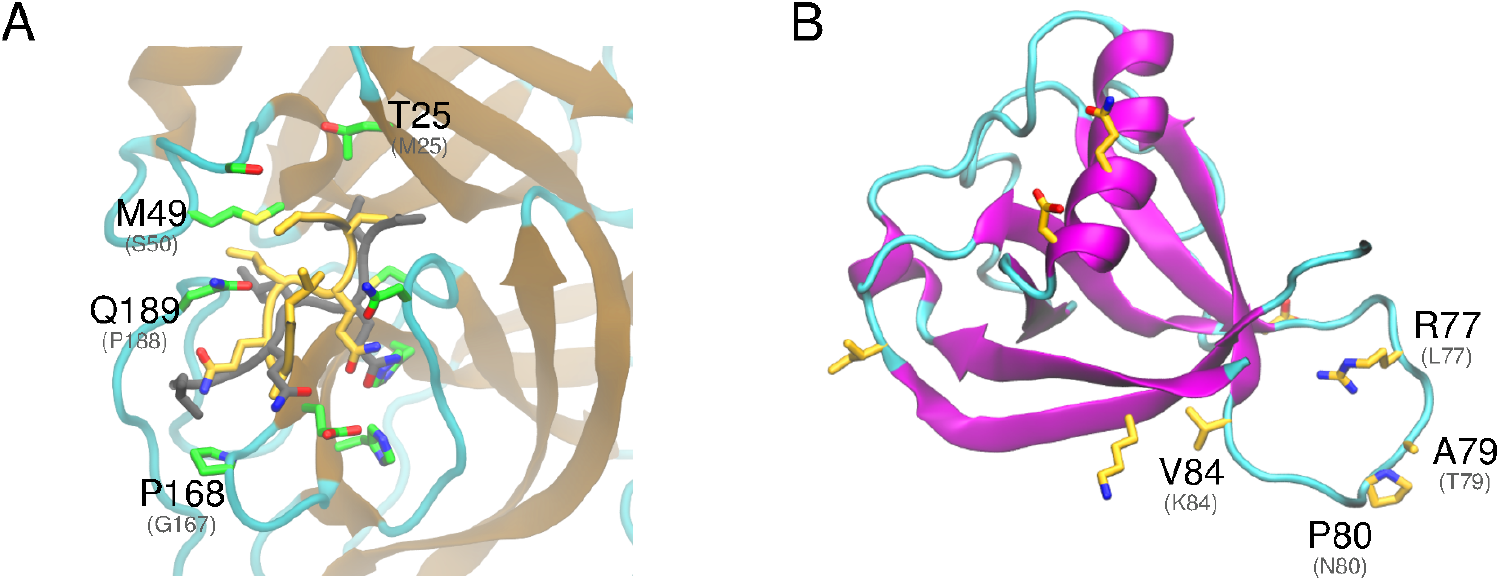
Analysis of the structural variation in SARS-CoV-2 nsp1 and nsp5 proteins. A) Close view of the catalytic site of nsp5. In *yellow*, NF-kB essential modulator (NEMO) is shown in the conformation predicted with docking. The conformation of NEMO in the PEDV 3CL^pro^ is also depicted, in *grey* (PDB id: 5zqg). B) Predicted fold of nsp1. Non-conservative substitutions relative to SARS-CoV-1 nsp1 are depicted in *yellow*. Substitutions discussed in the text are labeled, including the corresponding residue of the homologue (PEDV, in panel A and SARS-CoV-1, in B) in parentheses.

Nsp1 is highly conserved between SARS-CoV-1 and SARS-CoV-2. Notably, in SARS-CoV-2, there are four substitutions in the less conserved β3-4 loop (Fig. 10B), namely, Leu^77^Arg, Thr^79^Ala, Asn^80^Pro and Lys^84^Val. These substitutions may directly relate to pathogenicity as experimentally induced substitutions (Arg^73^Glu, Asp^75^Arg, Leu^77^Ala, Ser^78^Glu and Asn^80^Gly) in SARS-CoV-1 demonstrated increased inhibition of host gene expression, compared to the SARS-CoV-1 wild type (*44*), Subsequent experiments in mice showed that the deletion of this loop in SARS-CoV-1 resulted in increased survival rate and less severe lung damage (*45*). Given that this loop plays an essential role in the ability of nsp1 to impair host-translational activity, and the three substitutions in SARS-CoV-2 may be important elements of virulence divergence, this should be targeted in future studies that focus on disrupting infection.

### Interpretation and Concluding Remarks

A Systems Biology perspective lends itself well to combating the current COVID-19 pandemic and for future zoonotic outbreaks as they will likely require multiple solutions from different biological and epidemiological perspectives. Here, our analyses advocate for a re-evaluation of the currently accepted view that the primary means of infection for the SARS-CoV-2 virus is via association with ACE2 in lung tissue.

A non-respiratory path of infection has important implications for selecting the best methods to prevent the spread of SARS-CoV-2. If a major route of infection is nasal/oral that leads to strong intestinal colonization, and the poorer binding of SARS-CoV-2 Spike protein with ACE (compared to SARS-CoV-1), if any, results in delays in the colonization of lung tissue, this may explain why many people are considered asymptomatic as they do not display the traditional symptoms (cough, fever, pneumonia). Such individuals could thus be shedding virus through feces and poor hygiene long before they display traditional symptoms (if ever). A recent study indicates that fecal-mediated viral aerosolization from toilets may be a source of infective viral particles found in the air of bathrooms (<u>46</u>). It appears that there may be sufficient expression of ACE2 in nasal passages for SARS-CoV-2 to establish low level colonization that can lead to viral shedding via sneezing, nose blowing and lack of subsequent hand hygiene. Awareness of these routes of infection may offer ways to reduce the number and severity of COVID-19 cases.

Furthermore, the significant variation in ACE and ACE2 expression among organs is consistent with some of the symptomology being reported for COVID-19, e.g. gastrointestinal (*11*), testes (*13*), thyroid (fatigue (*14*, *15*)), brain epithelial cells (headache (*14*)), etc. In addition, there is wide variation in tissue-specific expression across the GTEx population, which may partially explain differences in susceptibility and severity of illness being observed in COVID-19. It is possible that the variation of ACE2 expression in the population is associated with some of the comorbidities found to be associated with mortality in COVID-19, including hypertension, cardiovascular disease and obesity. Furthermore, it is possible the fluid loss associated with diarrhea resulting from gastrointestinal colonization by SARS-CoV-2, may affect the RAS system as it tries to regulate blood volume in the body.

Even though we verified very weak binding of SARS-CoV-2 spike protein to ACE in our simulations, we urge that the hypothesis that ACE, which is highly expressed in lung tissue, is an alternative receptor for SARS-CoV-2 should be thoroughly tested experimentally in humans and potential zoonotic reservoirs. Our analyses suggest that ACE could be a secondary receptor for SARS-CoV-1 in the absence of ACE2, and it may have played a subtle but important role for the SARS outbreak. It could explain the lower incidence of respiratory distress in COVID-19 compared to SARS (*47*).

The expression analysis of the BAL samples indicates that COVID-19 likely induces increases in type I and type II alveolar epithelial cells, ciliated cells, Club cells, and lymphatic endothelial cells in BAL samples. There also seemed to be a significant repression of myeloid and lymphoid lineages, particularly CSFR1+ macrophages as well as their inflammatory signatures, which may be due to the induction of the Ang 1-7 receptor MAS by the virus as part of its strategy to escape detection. We posit that COVID-19 actively destroys these cells or reprograms them to the point of being unrecognizable. This apparent elimination of functional macrophages and the complete lack of activated cytokine signature in COVID-19 lung BAL samples would seem to indicate that SARS-CoV-2’s net effect of causing a **functional immune deficiency syndrome (COVID-19-FIDS)** is a considerable component of its virulence.

Along with the information of host proteins involved in infection, we provide an extensive structural analysis of the viral proteome, all of which is available as a web resource (https://compsysbio.ornl.gov/covid-19/covid-19-structome/) and in the Appendix section of Supplementary Material. The collective analysis also informs the identification of promising drug, vaccine and diagnostic targets for COVID-19. The workflow developed for this study can readily be implemented in future efforts against pathogen outbreaks.

## Acknowledgements

This research used resources of the Oak Ridge Leadership Computing Facility, which is a DOE Office of Science User Facility supported under Contract DE-AC05-00OR22725. This research used resources of the Compute and Data Environment for Science (CADES) at the Oak Ridge National Laboratory. We thank Joao Gabriel F. M. Gazolla (Biosciences Division, Oak Ridge National Laboratory) for support in the design of figures.

## Contributions

DJ conceived of the study, raised funding, supervised the study, performed data analysis and participated in writing and editing. ETP predicted protein structures, performed molecular dynamics simulations and analysis, analyzed protein structures and mutants, and participated in the writing and editing. MG collected and curated datasets, performed data analysis, analyzed protein structures and mutants, and participated in the writing and editing. PJ predicted protein structures, analyzed protein structures and mutants and participated in the writing. MS organized data, predicted protein structures, analyzed protein structures and mutants and participated in the writing. MP, OD, BKA, AG analyzed protein structures and mutants and participated in the writing. DK, CA performed data analysis and figure creation. JP participated in the early conceptual framework. BA, JK, AM, EB analyzed transcriptomic data, derived reference signatures, and developed infrastructure for comparative analysis of test signatures. BA, JK, AM participated in writing.

## Data and materials availability

Include a note explaining any restrictions on materials, such as materials transfer agreements. Note accession numbers to any data relating to the paper and deposited in a public database; include a brief description of the data set or model with the number. If all data are in the paper and supplementary materials include the sentence “All data is available in the main text or the supplementary materials.” All data, code, and materials used in the analysis must be available in some form to any researcher for purposes of reproducing or extending the analysis.

## Supplementary Materials

Materials and Methods Supplementary Text Figs. S1 to S7

Tables S1 to S5

Supplementary References (*1-42*)

Appendix: Structural Analysis of the SARS-CoV-2 Proteome

