## Supplementary Materials for "Functional Immune Deficiency Syndrome via Intestinal Infection in COVID-19"

**This PDF file includes:**

Materials and Methods
Supplementary Text
Figs. S1 to S7
Tables S1 to S5
Appendix: Structural Analysis of the SARS-CoV-2 Proteome

### Materials and Methods

Gene expression analysis

FASTQ files were downloaded from the Sequence Read Archive on NCBI and trimmed using the default parameters in CLC Genomics Workbench (20.0.3). RNA-Seq analysis was performed using the latest version of the human transcriptome (GRCh38_latest_rna.fna) with the reference genome for the SARS-CoV-2 virus appended (NC_045512). Transcripts per million (TPM) were used for comparisons across groups and the Differential Expression for RNA-Seq tool from CLC, which uses log-transformed TPM fitted to a GLM was used to identify globally differentially significantly expressed genes (*1*). Expression was significant if the Bonferroni correction was less than 0.01, the fold-change was greater than 2, and the mean TPM for either the control or the experimental groups was greater than 25. Significant enrichment of the Gene Ontology terms (geneontology.org) was determined with an FDR <0.05. GO terms and their enrichment values were analyzed with REVIGO (revigo.irb.hr) to reduce redundancy of terms. A similarity score of 0.4 was used to be conservative within the homo sapiens GO database.

For nasal brushings and the BAL samples, the TPM were log2+1 transformed, tested for differential expression and then filtered for genes that were found to be highly correlated with ACE2 expression in human lung. For those genes, we extracted the human gene expression data for lung from GTEx (N=578 individuals) and collapsed gene transcripts to genes (N=38340). We calculated Pearson correlation coefficient for all pairs of genes and used those that had correlation of +0.80 and -0.65 for ACE2 as the filter.

ToppCell carries out extensive differential gene expression analysis between cell classes, subclasses, and tissues and allows for between-dataset and between-sample of origin evaluation of gene expression patterns. This enabled the comparison of ACE vs. ACE2 expression across different tissues and datasets.

Expression datasets used are available feom the NCBI Sequence Read Archive under the following accession numbers.

Nasal: SRR7586338, SRR7586357, SRR7586364, SRR7586406, SRR7586390, SRR7586351, SRR7586412, SRR7586334, SRR7586366, SRR7586352, SRR7586344, SRR7586340, SRR7586373, SRR7586347, SRR7586402, SRR7586365, SRR7586336, SRR7586401, SRR7586407, SRR7586360, SRR7586339, SRR7586368, SRR7586349, SRR7586379, SRR7586372, SRR7586392, SRR7586374, SRR7586320, SRR7586328, SRR7586358, SRR7586388, SRR7586408, SRR7586387, SRR7586395, SRR7586337, SRR7586353, SRR7586380, SRR7586370, SRR7586342, SRR8369129A, SRR7586398, SRR7586345, SRR7586399, SRR7586405, SRR7586409, SRR7586324, SRR7586396, SRR7586327, SRR7586354, SRR7586403, SRR7586356, SRR7586371, SRR7586343, SRR7586397, SRR7586350, SRR7586333, SRR7586318, SRR7586319, SRR7586394, SRR7586378, SRR7586348, SRR7586362, SRR7586391, SRR7586330, SRR7586389, SRR7586361, SRR7586393, SRR7586322, SRR7586341, SRR7586323, SRR7586410, SRR7586325, SRR7586383, SRR7586332, SRR7586346, SRR7586376, SRR7586369, SRR7586377, SRR7586355, SRR7586386, SRR7586363, SRR7586321, SRR7586404, SRR7586329, SRR7586385, SRR7586331, SRR7586384, SRR7586367, SRR7586411, SRR7586335, SRR7586382, SRR7586359, SRR7586400, SRR7586381, SRR7586375, SRR7586326, SRR7586317

BAL: SRR10571753, SRR10571655, SRR10571733, SRR10571663, SRR10571658, SRR10571722, SRR10571743, SRR10571665, SRR10571732, SRR10571742, SRR10571659, SRR10571737, SRR10571664, SRR10571725, SRR10571660, SRR10571721, SRR10571744, SRR10571756, SRR10571731, SRR10571741, SRR10571726, SRR10571728, SRR10571757, SRR10571736, SRR10571657, SRR10571724, SRR10571755, SRR10571720, SRR10571739, SRR10571738, SRR10571730, SRR10571719, SRR10571734, SRR10571752, SRR10571735, SRR10571662, SRR10571723, SRR10571727, SRR10571656, SRR10571754

Population level SARS-CoV-2 analysis

FASTA files and meta data were downloaded from the Global Initiative on Sharing All Influenza Data (GISAID) website. Sequences were aligned with CLC Genomics Workbench (20.0.3), trimmed to include only coding sequence, and then translated to protein. A Neighbor-Joining Tree was generated using Jukes-Cantor protein distance measure.

Molecular dynamics simulations

The solved structures of the complexes of SARS-CoV-1 and SARS-CoV-2 receptor binding domains (RBDs) bound to angiotensin-converting enzyme 2 (ACE2), referred as RBD1-ACE2 and RBD2-ACE2, respectively, were used as starting configuration for atomistic molecular dynamics (MD) simulations (PDB ids: 6m17 and 2ajf) (*2, 3*). MD simulations were performed with the 2020 version of GROMACS (*4*). Each system was simulated in triplicate. The CHARMM36 protein force field (*5*) was used with TIP3P water (*6*). For glycosylated proteins, the CHARMM carbohydrate force field was used (*7*).

Proteins were solvated with a 15 Å water layer in an octahedral box under periodic boundary conditions. Sodium and chloride ions were added at a concentration of ~0.16 M to ensure electroneutrality of the system. All ions present in the original crystallographic structures were kept for the simulations.

The system was first energy minimized for 5000 steps with steepest descent. This was followed by 2 phases of equilibration. In the first phase, 6 ns passed while applying positional constraints on alpha carbons except for those at binding interfaces. The temperature was gradually raised to 298.15 K. In the second phase, 20 ns passed during which positional constraints were applied to alpha carbons of the C-terminal domains of ACE and ACE2 and a few residues in the core of the spike protein's receptor-binding domain. In each simulation of the triplicate, atomic velocities were reinitialized from a Maxwell-Boltzmann distribution at 10 ns, using a different random seed in each case. Equilibration and production were run in the NPT ensemble, and a 2 fs time step was used. Temperature was maintained using velocity rescaling via a stochastic term that properly generates constant pressure-constant temperature ensembles (*8*); and a coupling constant of 1.0 ps. The Berendsen barostat (*9*) was used with a coupling constant of 1.0 ps to maintain a pressure of 1 atm. All systems were run with a total of 300 ns of production.

Simulation analyses

*Inter-residue contacts:* Time evolution of number of contacts was computed using the mindist utility of GROMACS and a distance cutoff within alpha carbons of 8 Å. The probability density of contacts was computed using the timeline plugin of VMD (*10*).

Visual analysis and the design of figures were made using the version 1.93 of VMD.

Ensemble workflow for protein structure prediction

To date, partial or full structures of five proteins from SARS-CoV-2 have been experimentally solved. In view of the urgency to understand the molecular machinery of SARS-CoV-2, we used an ensemble workflow to generate structural models of all unsolved structural and mature nonstructural viral proteins. Due to the performance of methods for protein structure prediction varying by complexity, protein sequences were carefully analyzed to optimize the combination of the state-of-the-art methods of protein structure prediction. As such, the resulting models have the highest possible resolution and maximum information with regards to the overall shape of each protein. Here, we provide a synopsis for each of the 27 mature viral proteins along with their structural models and additional important information, such as variability relative to SARS-CoV-1 and potential functional relevance to SARS-CoV-2.

Case-by-case protocols were generated based on a profile extracted from each sequence, consisting of two main factors:

1. Primary sequence-based information. Residues within conserved domains (Pfam (*11*)) and intrinsically disordered regions were identified using IuPred2 (*12*), which relies on the composition of amino acid segments and their tendency to form stable structural motifs. TMHMM (*13*) was used to predict the helical transmembrane protein regions based on a hidden Markov model. No β-barrel transmembrane proteins are coded for in SARS-CoV-2.
2. Availability of experimentally determined structures. PSI-BLAST was used to identify homologous with partial or full structures available in the Protein Data Bank (PDB) that could be used as templates for modeling.

Several SARS-CoV-1 proteins that are highly conserved have been solved experimentally and are available for our analysis. In order to maximize accurate translation of information from these structures, amino acid substitutions were analyzed to identify those that likely impact protein conformation. Examples of changes that affect protein structure are a hydrophobic side chain being replaced by a charged amino acid at the protein core or a substitution to proline (a helix "breaker") within a helical structure. In case such substitutions are not found, and the protein has more than 70% identity to the template, loops and substitutions are locally modeled (LM) using the Rosetta remodel (*14*) and fixbb (*15, 16*) applications, respectively. The comparison of recently released crystallographic structures with the models generated using carefully analyzed protein sequences and using LM for selected regions appears to be an effective approach (Table S5). Achieving high local resolution, especially in sites of substrate/ligand binding, can considerably enhance the results of subsequent studies for small molecule candidate identification using molecular docking. Although ensemble docking approaches are often applied to contend with the conformational flexibility of the protein target, refining the binding site based on structural information from homologs in the holo form, if available, is more suitable for identifying functional complexes.

Homology-based modeling is typically the optimal approach for cases in which the identity to the template is above 30%. The fragment-based (FB) approach of the I-TASSER43 workflow was used in cases where the range of identity was 30-70%, and to provide an alternative model to LM in regions of proteins harboring substitutions that would be expected to significantly affect protein conformation. In order to predict structures for proteins that do not have a crystal structure of a homolog available, we applied the trRosetta (*17*) workflow. Based on benchmarks of the Critical Assessment of Techniques for Protein Structure Prediction (CASP13), trRosetta was designed to achieve sound performance for modeling novel folds by using a deep residual network for predicting inter-residue distance and orientation that guides energy minimization.

We use the analysis of nsp3, the largest mature protein of SARS-CoV-2, as an example for the workflow (Fig. S7): 1. The first model was generated using residues 5-111 via LM (78% identity to PDB 2GRI). This region partially overlaps with a conserved domain (DUF3655), which is predicted by ANCHOR2 to be a disordered binding region; 1b. A fragment-based model that expands to the end of this domain (segment 2-169) suggests that it undergoes disorder-to-order transition upon interaction with the N-terminal (Fig. 6B). 2. The flexible linker following DUF3655 was skipped and the next segment (205-374), corresponding to the ADP ribose phosphatase domain of nsp3 was modeled based on the alignment with a highly similar template (73% identity, PDB id 2fav, SARS-CoV-1). A comparison with a recently solved crystal structure of this region (PDB id 6w02) reveals that the generated model is good (RMSD 1.5 Å); 3. The preliminary analysis of the substitutions in the segment composed of residues 413-676 relative to the template from SARS-CoV-1 (2w2g) demonstrates that most of the non-conservative substitutions occur in superficial regions, but several suggest that conformational divergence should be considered. In order to account for local rearrangements in this region, which includes the SUD-M domain, a FB model was generated as an alternative to the LM model, since the I-TASSER workflow includes an atomistic refinement phase. Fig. 6C shows the amino acid substitutions in the LM model aligned to the template. The magnified section depicts the conformational change captured by the FB model due to the loss of the disulfide bridge present on the template. 4. Residues 680-743 correspond to PL2pro, the coronavirus polyprotein cleavage domain. Although a psi-blast search identifies a solved homolog covering this region, this domain is missing in the structure. Due to the lack of any template, this domain was modeled ab initio, with the trRosetta workflow; 5. Residues 746-1062 correspond to the papain-like viral protease, which acts in the proteolytic processing of viral replicase. It was locally modeled using the homolog from SARS-CoV-1 (PDB id 5tl6), which is 82% identical; 6. Similarly, the subsequent residues 1089-1203 correspond to the putative nucleic acid binding domain of nsp3, was modeled via LM using the homolog from SARS-CoV-1 (PDB id 2k87), which is 81% identical. No other conserved regions were identified in nsp3. The C-terminal region, which is predicted to contain transmembrane segments, was not modeled, since the soluble functional domains of this protein are of highest interest, and as there is no previous structural information to ensure the generation of a reliable model for it.

Molecular docking

Molecular docking of the nuclear factor-kappa B essential modulator (NEMO) targeting SARS-CoV-2 and PEDV 3CLpro proteins was performed using Autodock Vina (*18*). Autodocktools was used to prepare the inputs (*19*). The search space was defined as a box with dimensions 20 x 20 x 20 Å, encompassing the side chains of the full catalytic site of these enzymes. Grid space 1·0 Å was used, and exhaustiveness parameter was set 20. The N-Cα and Cα-C bonds in the segment $Gln^{229}$-$Ala^{233}$ and bonds in the side chains of $Leu^{227}$, $Leu^{230}$, $Val^{232}$ and $Ala^{233}$ were set as flexible, except for those forming $\pi$-conjugated systems. The remaining bonds were fixed in the conformation of NEMO in the crystal structure of PEDV nsp5-NEMO (PDB id 5zqg), as the results of Vina are often more accurate for a number of active bonds lower than 15 (*20*).

Tracking SARS-CoV-2 molecular evolution

Resources are available to track and infer functional consequences of mutations as the virus spreads globally. We aligned the full protein coding sequences of all available SARS-CoV-2 isolates (N=1,495 as of this publication) and identified the frequency and geographic associations of the new mutations as well as their proteome location. The most frequent mutation (72%) at

position 84 in the ORF8 protein from an ancestral serine to leucine has been proposed to increase the aggressiveness of the virus, allowing it to spread further.(*21*) This idea seems to also be consistent with the less effective spread of SARS-CoV-1, in which ORF8b has a threonine in this site (it has similar chemical properties of serine).
Another important mutation is the double substitution in the nsp13 helicase ($Pro^{504}$Leu, $Tyr^{541}$Cys), frequently found in Washington State, USA. Interestingly, nsp13 is absolutely conserved within SARS-like coronaviruses.(*22*) Both substitutions are located in the 2A (a RecA like) domain of the protein, with $Tyr^{541}$ predicted to be part of a critical region for nucleic acid binding.(*23*) In fact, in previous work, it was shown that a the double substitution $Ser^{539}$Ala/$Tyr^{541}$Ala decreased helicase unwinding activity.(*24*) Based on those results, it is likely that the loss of the tyrosine, a bulky aromatic amino acid, will impact nucleic acid binding efficiency. The binding region is also associated with the RNA triphosphatase activity of nsp13, thus the mutation may affect the viral 5' RNA capping, and thereby viral replication.(*25*)
Finally, we also find a frequent (41%) Asp to Gly substitution at position 614 in the end of S1 subunit of the spike protein. We hypothesize that $Asp^{614}$ counterbalances the attraction of $Arg^{646}$ to $Glu^{868}$ in the subunit S2 of the adjacent monomer, and, therefore, this mutation may enhance the trimer packing by releasing $Arg^{646}$ to mainly interact with $Glu^{868}$ (Fig. S5). This site is linked to a second mutation ($Pro^{323}$Leu) in nsp12. Other notable findings are a Scandanavian-specific strain defined by a Thr to Met substitution in the membrane protein and a Leu to Phe in the nsp6 gene of the viral type infecting individuals in Japan.

Single-cell RNA-seq Analysis

Mouse cell Atlas and Human Cell Landscape Single-cell RNA-seq data were downloaded from Tabula Muris and Human Cell Landscape platforms. Total UMI counts per cell were normalized to one million to compare across tissues. Cell classes and subclasses were drawn from metadata in Tabula Muris and Human Cell Landscape compendium and were amended by cell annotations of scRNA-Seq data from other well-annotated datasets (*i.e.* LungMap) for further labelling. Cells were collapsed into bins by their identities, including tissue, lineage, class and subclass to support visualization. The 75 percentile of expression values of cells in the bins was used as their expression levels. Using the annotations, a hierarchy was defined for each bin and gene lists were created to represent the hierarchical gene expression modules. Differential expression analysis was carried out by t-test and the top two hundred genes of each module were used to create a heatmap using Morpheus. Rows were collapsed to the mean expression of ACE and ACE2 across modules and cell subclasses from Mouse Cell Atlas. Gene modules with ACE or ACE2 in their gene lists were represented for Human Cell Landscape.
scRNA-seq data of respiratory tissues were downloaded from GSE12260 (including samples from nasal brushing, nasal turbinate and bronchial biopsy) and Tissue Stability Cell Atlas (including samples from parenchyma). Mouse Tongue scRNA-seq data were extracted from Mouse Cell Atlas. We used the package SCANPY(*26*) to filter out cells with less than 500 total UMI counts or more than 15% mitochondrial reads. Total UMI counts were normalized to one million per cell. The top 3,000 highly variable genes were calculated, and normalized values were scaled to unit variance. PCA was carried out on highly variable genes and the top 50 principle components were selected. Louvain clustering and UMAP visualization were used to cluster the cells and gain a better view of the cell distribution. We used known marker genes to label major cell types and refined our annotations by mapping signatures from the well-annotated scRNA-seq datasets. Gene

expression levels of ACE and ACE2 were shown on the same UMAP and compared across respiratory tissues.

### Supplementary Text

ACE2 is not highly expressed in normal lung tissue

Both the Genotype-Tissue Expression (GTEx) and Proteomics DataBase (Proteomics DB) indicate ACE2 is either expressed at very low levels or is not detectable in tissues that are currently thought to be major entry points for SARS-CoV-1 and SARS-CoV-2 such as lung. High expression of ACE2 is based on several reports that can be divided into those that use mRNA and those that use protein.(*27–33*) The two earliest reports that measured ACE2 mRNA in lung tissue showed either no expression at all or low levels (*33*, *34*) whereas ACE is found at moderate to high levels; neither were quantitative and both were based on small sample sizes. The most recent reports used data from single cell RNA-Seq experiments (and those results agree with data from the GTEx project) and show that ACE2 is expressed at very low levels in a small fraction of cells (ACE was not reported).(*31*, *32*) ACE vs ACE2 in single cell atlas datasets from human tissues.

The earliest reports of ACE2 expression in the lung were based on protein expression using a peptide-derived polyclonal antibody and immunohistochemistry.(*27*, *28*) Aside from being non-quantitative, the results are difficult to interpret and could be due to the non-specificity of these types of antibodies. The three remaining reports of ACE2 expression in lung or lung-derived epithelium were based on improved antibody immunohistochemistry and western blots.(*29*, *30*, *35*) Although still not quantitative, the reports are again consistent with GTEx and Proteome DB; ACE2 is either not expressed or found at very low levels in normal lung tissue or normal lung-derived epithelial cells. However, it is induced either by exposing cells grown in culture to air (air-liquid interface) or by infection with the SARS-CoV-1 virus, suggesting that compromised lung epithelium or cells that are already infected with virus are routes of infection in this tissue.

ACE may mediate SARS-CoV-1 infection

The involvement of ACE in the system that was used to identify ACE2 as the SARS-CoV-1 receptor is unknown because they did not overexpress it as was done with ACE2.(*36*) However, a subsequent study using pseudovirus typing demonstrated a correlation between ACE2 expression and infectivity in several cell lines and although ACE2 was much higher, overexpression of ACE was shown to increase infectivity in some cell lines.(*37*) Notably, ACE2-mediated infection was highest in kidney- and colon-derived cell lines and much less efficient in those from the lung. This same pattern in kidney-, colon-, and lung-derived cell lines were confirmed in a more recent analysis of beta-coronaviruses.(*38*) From these combined results we conclude that while ACE2 is involved in SARS-CoV-1 infection in kidney-derived cells, the involvement of ACE2 in lung-mediated viral entry and ACE in viral infection in general is inconclusive. It should be noted that the majority of positive results were garnered in kidney- and colon-derived cell lines, which express higher levels of ACE2 than ACE. In the lung and other respiratory tissues, the reverse is true; ACE is expressed at higher levels than ACE2. From *in vitro* work it appears that ACE may be a less efficient receptor for SARS-CoV-1 (and perhaps SARS-CoV-2) but, given an environment in which levels of ACE are an order of magnitude higher than ACE2, this may be a potential route of infection in respiratory tissue.

Gene Expression of BAL Samples from COVID-19 patients

Interestingly, in addition to the high representation of the raft-127 genes in those that were differentially expressed, the vast majority (98%) of the 2,057 downregulated genes were protein coding whereas only 41% of the upregulated genes coded for protein and 40% were non-coding RNAs (ncRNA). One of the most upregulated ncRNA (LOC10537114, a 3,171 fold increase) is associated with lung cancer susceptibility.(39) A Gene Ontology (GO) biological enrichment analysis identified no significant enrichment within the upregulated protein coding genes but the downregulated genes demonstrated significant enrichment of the adaptive and innate immune responses, including the NF-κB pathway, which is known to be inhibited by coronavirus during infection.(40) In addition, trafficking among the Endoplasmic Reticulum (ER), Golgi, and endosomal compartments was highly enriched for many categories. Other enriched terms implicated lipid rafts, apoptosis, protein production, and respiration.

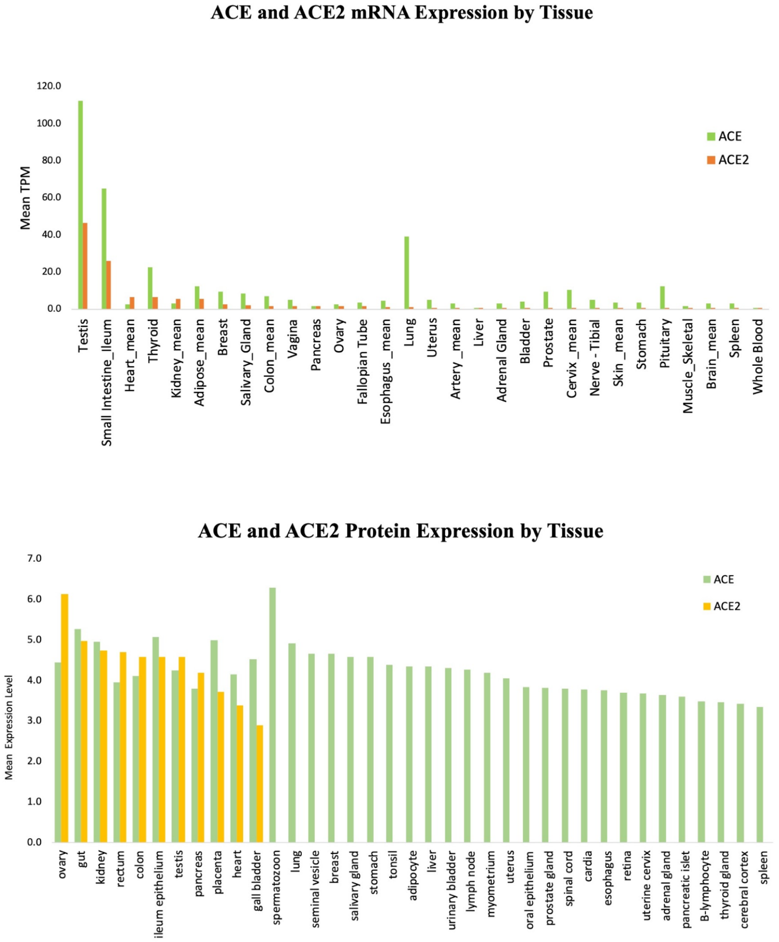


**Fig. S1**.

Expression values for ACE and ACE2 from GTEx and Proteome DB databases. mRNA are given as mean TPM (transcripts per million).

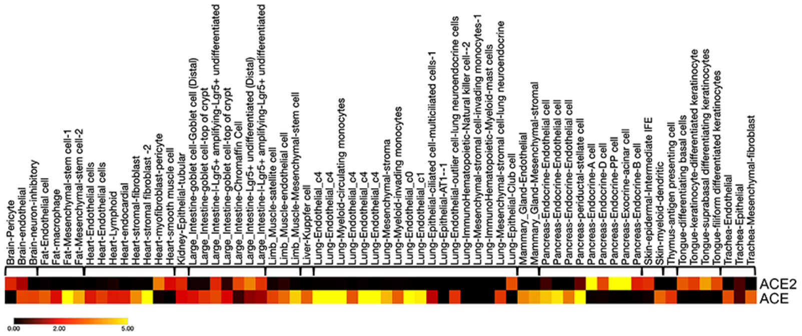


**Fig. S2.**

ACE2 vs. ACE expression in Mouse Tabula Muris. Like human tissues and cell types, ACE2 exhibits highly selective expression patterns in mice and is expressed in select epithelial cells of the tongue and GI tract, including pancreatic cells and enterocytes. In the lung, only Club cells have significant ACE2 expression, albeit at 4-fold or lower levels vs pancreatic or enterocyte subtypes. Pericytes in the brain and heart are notably positive for ACE2, but lung pericytes do not demonstrate significant expression. Similar to the nasal epithelium brushing samples, mouse tongue expression of ACE2 corresponds to developing and differentiating keratinocytes. Color scales are Log2(TPM+1).

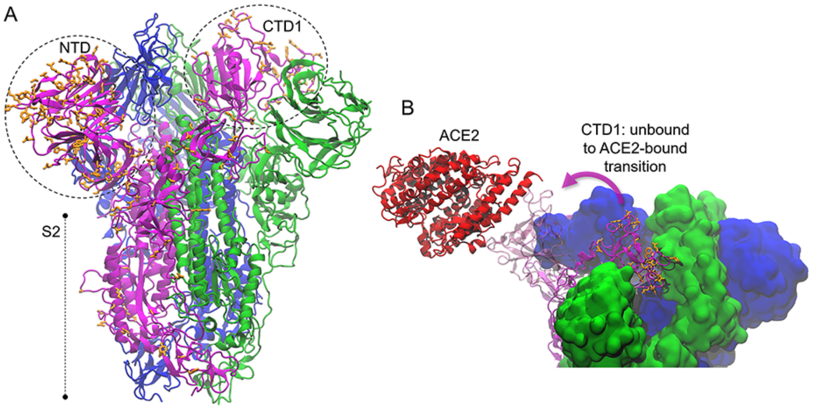


**Fig. S3.**

A) Local modeling-derived SARS-CoV-2 spike glycoprotein. B) Conformational transition of the receptor-binding domain of the S1 subunit of the spike glycoprotein and association with ACE2 receptor. Non-conservative substitutions relative to SARS-CoV-1 spike are depicted in orange. N-terminal (NTD) and C terminal domains (CTD) are identified.

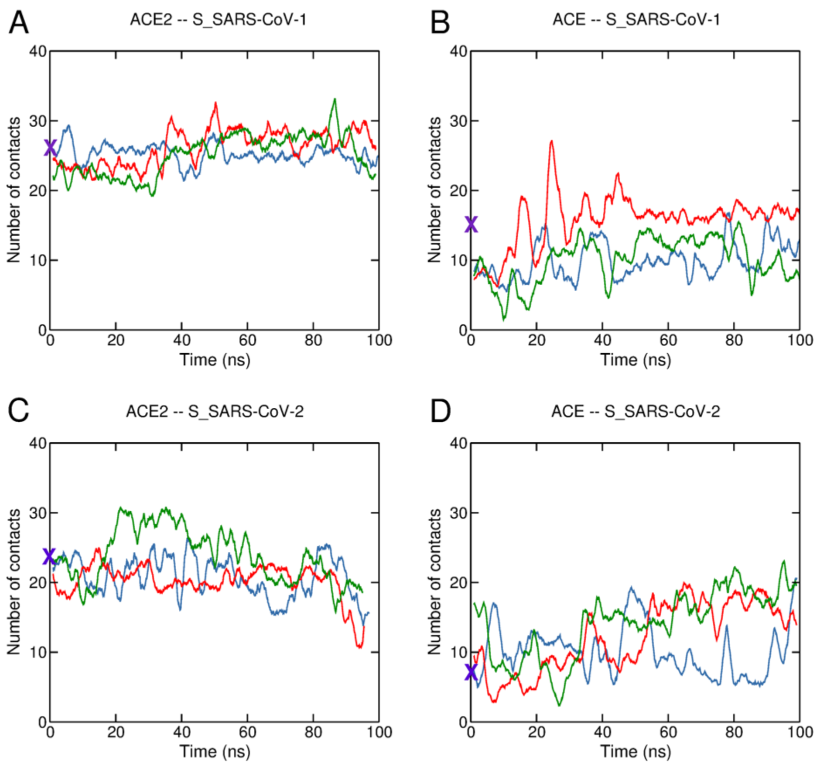


**Fig. S4.**

Time evolution of the number of contacts within the receptor binding domain (SARS-CoV-1 and SARS-CoV-2 spike glycoproteins) and the putative receptors ACE2 and ACE. A contact was considered for pairs of residues with C-alpha less than 8 Å distant.

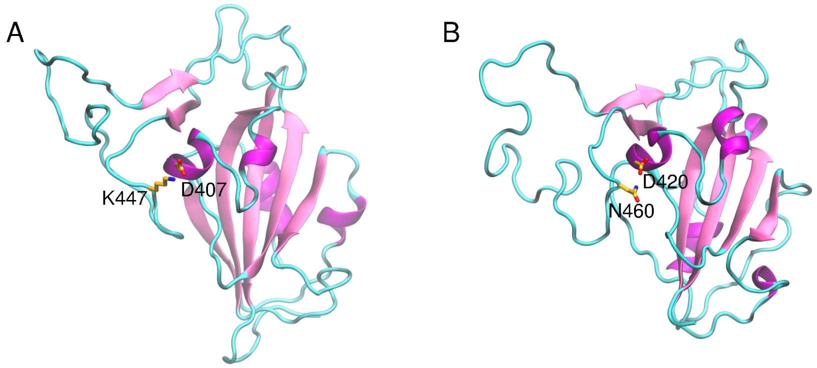


**Fig. S5.**

Substitution of Lys447 in SARS-CoV-1 S protein (A) by Asn$^{460}$ in SARS-CoV-2 S (B) protein results in loss of a salt bridge connecting a long loop to RBD core.

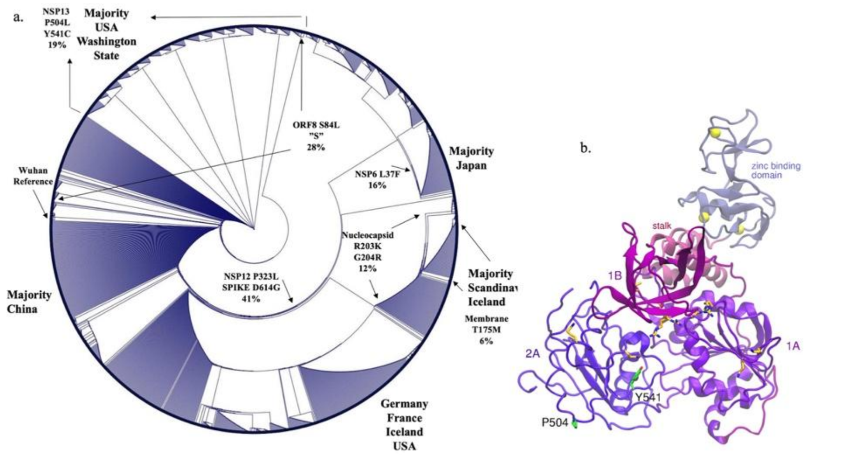


**Fig. S6**.

A) Neighbor-joining circular dendrogram of 1495 protein sequences for SARS-CoV-2 from GSAID. Frequent mutations that define branch points are given as well as some geographic-specific clades (e.g., the Thr to Meth substitution in the Membrane protein at site 175 defines a sub-group of viruses in Scandinavia). Similarly, a linked set of mutations is currently spreading in Washington State in the USA, which may have functional implications on the (B) Predicted structure of SARS-CoV-2 nsp13. Non-conservative substitutions relative to SARS-CoV-1 nsp13 are depicted in orange. Mutations observed in SARS-CoV-2 nsp13 are depicted in green. Zinc ions are represented as yellow spheres.

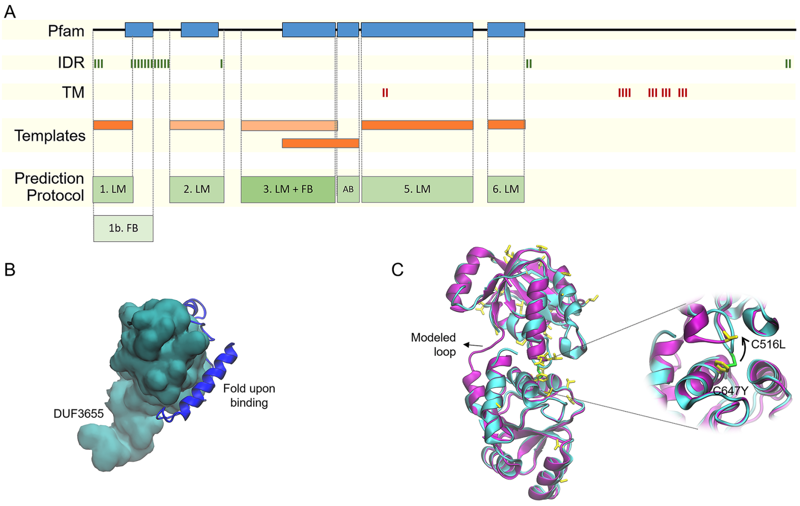


**Fig. S7.**

Ensemble workflow for structure prediction of SARS-CoV-2 nsp3. Case-by-case protocols of structure prediction are determined by finely parsing each protein sequence using information about the position of intrinsically disordered regions (IDR), transmembrane regions (TM), signal peptides, and templates (Fig. S1A). The method applied to SARS-CoV-2 nsp3 defined six regions to be modeled by the local modeling (LM), fragment-based (FB) and/or *ab initio* (AB) approaches. The first region, 2-169, of nsp3 sequence was modeled using the combination LM+FB (Fig. S1B), so that the structured region (5-111) could be determined in high resolution via LM of the side chain using a highly similar template (PDB 2gri, identity 76%). The bound conformation of intrinsically disordered segments was predicted using FB, a more flexible method. The choice of prediction method for protein regions with templates of high identity includes a thorough structural analysis of the template. For example, the region 413-676 of nsp3 is aligned with a high identity template (2w2g, 76%), so that modeling only variant side chains was considered. However, due to the predicted loss of a disulfide bridge, the FB approach was used since it allows larger conformational changes (C).

**Table S1.**

Allele frequencies for the *Alu* insertion (I) and deletion (D) in the ACE gene across 199 populations from the ALlele FREquency Database (ALFRED) at the Yale Center for Medical Informatics.

| Population | I | D |
|---|---|---|
| Africa_Kung_San | 0·35 | 0·65 |
| Africa_Nguni | 0·42 | 0·58 |
| Africa_San | 0·31 | 0·69 |
| Africa_Sotho-Tswana | 0·32 | 0·68 |
| Africa_Tsonga | 0·22 | 0·78 |
| Africa_African_Pygmies | 0·52 | 0·48 |
| Africa_Afro-Caribbeans | 0·26 | 0·74 |
| Africa_Biaka | 0·44 | 0·56 |
| Africa_Hausa | 0·41 | 0·59 |
| Africa_Ibo | 0·38 | 0·62 |
| Africa_Mbuti | 0·35 | 0·65 |
| Africa_Yoruba | 0·37 | 0·63 |
| Africa_Chagga | 0·20 | 0·80 |
| Africa_Jews_Ethiopian | 0·38 | 0·62 |
| Africa_Masai | 0·46 | 0·54 |
| Africa_African_Americans | 0·41 | 0·59 |
| Africa_Berber | 0·29 | 0·71 |
| Africa_Moroccans | 0·27 | 0·73 |
| Africa_Nigerians | 0·33 | 0·68 |
| Africa_Tunisian | 0·33 | 0·67 |
| Asia_Anatolian | 0·29 | 0·71 |

| | | |
|---|---|---|
| Asia_Arabs | 0·48 | 0·52 |
| Asia_Armenian | 0·22 | 0·78 |
| Asia_Azerbaijani | 0·35 | 0·65 |
| Asia_Bahraini | 0·50 | 0·50 |
| Asia_Druze | 0·33 | 0·67 |
| Asia_Emirati | 0·40 | 0·61 |
| Asia_Jews_Yemenite | 0·38 | 0·63 |
| Asia_Jordanian | 0·44 | 0·56 |
| Asia_Samaritans | 0·28 | 0·73 |
| Asia_Saudi | 0·00 | 1·00 |
| Europe_Abazians | 0·55 | 0·45 |
| Europe_Adygei | 0·47 | 0·53 |
| Europe_Albanian | 0·43 | 0·57 |
| Europe_Aromanian | 0·39 | 0·62 |
| Europe_Azorian | 0·52 | 0·49 |
| Europe_Bashkir | 0·39 | 0·61 |
| Europe_Basque | 0·43 | 0·57 |
| Europe_Bosniaks | 0·38 | 0·62 |
| Europe_Bretons | 0·37 | 0·63 |
| Europe_Canarian | 0·30 | 0·70 |
| Europe_Cherkess | 0·54 | 0·46 |
| Europe_Chuvash | 0·46 | 0·54 |
| Europe_Croatian | 0·37 | 0·63 |
| Europe_Cypriot_Greek | 0·35 | 0·65 |
| Europe_Cypriot_Turkish | 0·50 | 0·50 |
| Europe_Czech | 0·42 | 0·58 |

| | | |
|---|---|---|
| Europe_Danes | 0·17 | 0·83 |
| Europe_Darghinian | 0·45 | 0·55 |
| Europe_English | 0·51 | 0·49 |
| Europe_European_Americans | 0·40 | 0·60 |
| Europe_Europeans_Mixed | 0·50 | 0·50 |
| Europe_Finns | 0·42 | 0·58 |
| Europe_French | 0·46 | 0·54 |
| Europe_Gaguazes | 0·40 | 0·60 |
| Europe_Galician | 0·35 | 0·65 |
| Russia_Georgians | 0·31 | 0·69 |
| Europe_Greeks | 0·34 | 0·66 |
| Europe_Ingush | 0·43 | 0·57 |
| Europe_Irish | 0·34 | 0·66 |
| Europe_Italians | 0·34 | 0·66 |
| Europe_Jews_Ashkenazi | 0·27 | 0·73 |
| Europe_Kabardinian | 0·61 | 0·39 |
| Europe_Kalmyks | 0·53 | 0·48 |
| Europe_Karachay | 0·38 | 0·62 |
| Europe_Kumyk | 0·48 | 0·52 |
| Europe_Macedonian | 0·31 | 0·69 |
| Europe_Mari | 0·48 | 0·52 |
| Europe_Moldovan | 0·70 | 0·30 |
| Europe_Mordvin | 0·46 | 0·54 |
| Europe_Nogay | 0·37 | 0·63 |
| Europe_Portuguese | 0·45 | 0·55 |
| Europe_Roma | 0·43 | 0·57 |

| | | |
|---|---|---|
| Europe_Romanian | 0·57 | 0·43 |
| Europe_Russians | 0·28 | 0·72 |
| Europe_Sardinian | 0·39 | 0·61 |
| Europe_Serb | 0·41 | 0·59 |
| Europe_Spaniards | 0·38 | 0·62 |
| Europe_Swiss | 0·40 | 0·61 |
| Europe_Tatar | 0·54 | 0·46 |
| Europe_Udmurt | 0·40 | 0·60 |
| Europe_Ukrainian | 0·62 | 0·38 |
| Asia_Burusho | 0·57 | 0·43 |
| Asia_Kalash | 0·44 | 0·56 |
| Asia_Pakistani | 0·54 | 0·46 |
| Asia_Pashtun | 0·60 | 0·40 |
| Asia_Dungan | 0·56 | 0·44 |
| Asia_Indian_Mixed | 0·55 | 0·45 |
| Asia_Kazakh | 0·85 | 0·15 |
| Asia_Khanty | 0·59 | 0·41 |
| Asia_Komi-Zyrian | 0·63 | 0·37 |
| Asia_Kyrgyz | 0·44 | 0·56 |
| Asia_Tajik | 0·44 | 0·56 |
| Asia_Turks | 0·57 | 0·43 |
| Asia_Uzbek | 0·42 | 0·58 |
| Asia_Agharia | 0·48 | 0·52 |
| Asia_Ambalakarer | 0·46 | 0·54 |
| Asia_Badaga | 0·68 | 0·32 |
| Asia_Bagdi | 0·55 | 0·45 |

| | | |
|---|---|---|
| Asia_Brahmin | 0·70 | 0·30 |
| Asia_Chakma | 0·70 | 0·30 |
| Asia_Chamar | 0·60 | 0·40 |
| Asia_Gaud | 0·61 | 0·39 |
| Asia_Gond | 0·65 | 0·35 |
| Asia_Halba | 0·68 | 0·32 |
| Asia_Irula | 0·68 | 0·32 |
| Asia_Jamatiya | 0·64 | 0·36 |
| Asia_Kamar | 0·61 | 0·39 |
| Asia_Kapu | 0·68 | 0·32 |
| Asia_Kashmiri | 0·61 | 0·39 |
| Asia_Khonda_Dora | 0·59 | 0·41 |
| Asia_Kotas | 0·55 | 0·45 |
| Asia_Kshatriya | 0·81 | 0·19 |
| Asia_Kurumba | 0·86 | 0·14 |
| Asia_Lodha | 0·69 | 0·31 |
| Asia_Madiga | 0·56 | 0·44 |
| Asia_Mahishya | 0·69 | 0·31 |
| Asia_Mala | 0·58 | 0·42 |
| Asia_Mog | 0·59 | 0·41 |
| Asia_Moor_Sri_Lanka | 0·64 | 0·36 |
| Asia_Munda | 0·53 | 0·47 |
| Asia_Muria | 0·58 | 0·42 |
| Asia_Nepalese | 0·65 | 0·35 |
| Asia_Pallan | 0·56 | 0·45 |
| Asia_Parsi | 0·54 | 0·46 |

| | | |
|---|---|---|
| Asia_Rajput | 0·55 | 0·45 |
| Asia_Relli | 0·62 | 0·38 |
| Asia_Riang | 0·47 | 0·53 |
| Asia_Santal | 0·62 | 0·38 |
| Asia_Sri_Lankan | 0·65 | 0·35 |
| Asia_Tamil | 0·45 | 0·55 |
| Asia_Tanti | 0·60 | 0·40 |
| Asia_Tharu | 0·60 | 0·41 |
| Asia_Tipperah | 0·47 | 0·53 |
| Asia_Toda | 0·62 | 0·38 |
| Asia_Vanniyar | 0·66 | 0·34 |
| Asia_Vellala | 0·44 | 0·56 |
| Asia_Vysya | 0·53 | 0·47 |
| Asia_Yadava | 0·55 | 0·45 |
| EastAsia_Ami | 0·62 | 0·38 |
| EastAsia_Atayal | 0·70 | 0·31 |
| Asia_Ewenki | 0·61 | 0·39 |
| EastAsia_Han | 0·42 | 0·58 |
| EastAsia_Japanese | 0·58 | 0·43 |
| EastAsia_Koreans | 0·58 | 0·42 |
| EastAsia_Sibo | 0·50 | 0·50 |
| EastAsia_Taiwanese | 0·57 | 0·43 |
| EastAsia_Uyghur | 0·79 | 0·21 |
| EastAsia_Cambodians_Khmer | 0·53 | 0·47 |
| EastAsia_Filipino | 0·68 | 0·32 |
| EastAsia_Hakka | 0·62 | 0·39 |

| | | |
|---|---|---|
| EastAsia_Moluccas | 0·63 | 0·38 |
| EastAsia_Nusa_Tengarras | 0·56 | 0·44 |
| EastAsia_Vietnamese | 0·86 | 0·14 |
| EastAsia_Javanese | 0·84 | 0·16 |
| EastAsia_Malaysians | 0·48 | 0·52 |
| Oceania_New_Zealander | 0·70 | 0·30 |
| Oceania_Papuan_New_Guinean | | |
| Oceania_Australian | 1·00 | 0·00 |
| Oceania_Melanesian_Nasioi | 0·59 | 0·41 |
| Oceania_Micronesians | 0·64 | 0·36 |
| Siberia_Buryat | 0·78 | 0·22 |
| Siberia_Chukchi | 0·80 | 0·20 |
| Siberia_Siberian_Eskimo | 0·55 | 0·45 |
| Siberia_Tuva | 0·63 | 0·37 |
| Siberia_Yakut | 0·52 | 0·48 |
| NorthAmerica_Alaskan_Natives | | |
| NorthAmerica_Cheyenne | 0·49 | 0·51 |
| NorthAmerica_French_Acadians | | |
| NorthAmerica_Hispanic_American | | |
| NorthAmerica_Inuit_Canadian | 0·57 | 0·43 |
| NorthAmerica_Inuit_Greenland | | |
| NorthAmerica_Muscogee | 0·74 | 0·26 |
| NorthAmerica_Na-Dene | 0·89 | 0·11 |
| NorthAmerica_Navajo | 0·56 | 0·44 |
| NorthAmerica_Pima_Arizona | 0·54 | 0·46 |
| NorthAmerica_Pima_Mexico | 0·77 | 0·23 |

| | | |
|---|---|---|
| NorthAmerica_Zuni | 0·68 | 0·33 |
| NorthAmerica_Maya_Yucatan | 1·00 | 0·00 |
| SouthAmerica_Ache | 0·63 | 0·37 |
| SouthAmerica_Ayoreo | 0·50 | 0·50 |
| SouthAmerica_Bari | 0·42 | 0·58 |
| SouthAmerica_Brazilian | 0·73 | 0·27 |
| SouthAmerica_Chimila | 0·81 | 0·19 |
| SouthAmerica_Cinta_Larga | 0·93 | 0·07 |
| SouthAmerica_Gaviao | 0·83 | 0·17 |
| SouthAmerica_Guarani | 0·88 | 0·12 |
| SouthAmerica_Guihiba | 0·54 | 0·46 |
| SouthAmerica_Kaingang | 0·95 | 0·06 |
| SouthAmerica_Karitiana | 0·98 | 0·02 |
| SouthAmerica_Parakana | 0·73 | 0·27 |
| SouthAmerica_Quechua | 0·92 | 0·08 |
| SouthAmerica_Surui | 0·81 | 0·19 |
| SouthAmerica_Ticuna | 0·98 | 0·02 |
| SouthAmerica_Wai-Wai | 0·68 | 0·32 |
| SouthAmerica_Xavante | 0·75 | 0·25 |
| SouthAmerica_Yanomami | 0·94 | 0·06 |
| SouthAmerica_Yuco | 096 | 0·04 |
| SouthAmerica_Zoro | 0·00 | 1·00 |

**Table S2.**

Raft-127 genes identified in Kowalski et al. 2007 (*25*) and their differential expression in BAL samples.

| Symbol | Lipid rafts of cells infected with *P. aeruginosa* | DEGs |
|---|---|---|
| SPC25 | microsomal signal peptide 25 kDa subunit | UP |
| TF | serotransferrin | UP |
| AGTRAP | angiotensin II receptor-associated protein | DOWN |
| AHNAK | Ahnak | DOWN |
| ARF1 | ADP ribosylation factor 1 | DOWN |
| BANF1 | barrier to autointegration factor 1 | DOWN |
| CD81 | CD81 | DOWN |
| CLIC1 | nuclear chloride ion channel 27 | DOWN |
| CLPTM1 | cleft lip and palate associated transmembrane | DOWN |
| COX5A | cytochrome c oxidase subunit Va | DOWN |
| CTSD | cathepsin d | DOWN |
| CYCS | cytochrome c | DOWN |
| FLNA | filamin a | DOWN |
| FLOT2 | flotillin 2 | DOWN |
| GLUD1 | glutamate dehydrogenase 1 | DOWN |
| GOLGA7 | Golgi autoantigen, golgin subfamily a, 7 | DOWN |
| HINT1 | histidine triad nucleotide binding protein 1 | DOWN |
| HM13 | minor histocompatibility antigen h13 | DOWN |
| HNRNPF | hnrnp f | DOWN |

| | | |
|---|---|---|
| IMPDH2 | inosine monophosphate dehydrogenase 2 | DOWN |
| IQGAP1 | ras GTPase-activating-like protein IQGAP1 | DOWN |
| KPNB1 | importin beta 1 | DOWN |
| LAMTOR5 | hepatitis b virus x interacting protein | DOWN |
| LDHA | L-lactate dehydrogenase | DOWN |
| LMNA | lamin a | DOWN |
| MDH1 | malate dehydrogenase | DOWN |
| MGST1 | microsomal glutathione S-transferase 1 | DOWN |
| MGST3 | microsomal glutathione s-transferase 3 | DOWN |
| MIF | macrophage migration inhibitory factor | DOWN |
| MSN | Moesin | DOWN |
| MVP | major vault protein | DOWN |
| MYDGF | C19orf10 protein | DOWN |
| MYH9 | myosin heavy chain, nonmuscle, differentially spliced | DOWN |
| NDUFS3 | NADPH-ubiquinone oxidoreductase 30 kDa | DOWN |
| NME1 | nucleoside diphosphate kinase a | DOWN |
| NME2 | nucleoside diphosphate kinase b | DOWN |
| OTUB1 | ubiquitin-specific protease otubain 1 | DOWN |
| PDIA6 | protein disulfide isomerase-associated 6 | DOWN |
| PHB2 | prohibitin 2 | DOWN |
| PPIF | peptidylprolyl isomerase F | DOWN |
| PRDX1 | peroxiredoxin 1 | DOWN |
| RAB11B | rab11b | DOWN |

| | | |
|---|---|---|
| RAB5C | rab5c | DOWN |
| RAP1A | rap1a | DOWN |
| RAP1B | rap1B | DOWN |
| RER1 | Rer1 | DOWN |
| RHOA | rhoa | DOWN |
| RPS17 | 40s ribosomal protein s17 | DOWN |
| S100A8 | calgranulin B | DOWN |
| SOD1 | superoxide dismutase | DOWN |
| ST13 | hsc70-interacting protein | DOWN |
| SYPL1 | pantophysin | DOWN |
| TFRC | transferrin receptor protein 1 | DOWN |
| TMED10 | transmembrane protein tmp21 | DOWN |
| TOMM20 | mitochondrial import receptor subunit | DOWN |
| TPM3 | tropomyosin 3 | DOWN |
| VAMP2 | vesicle associated membrane protein 2 | DOWN |
| AK2 | adenylate kinase isoenzyme 2 | na |
| ALDOC | fructose bisphosphate aldolase c | na |
| AOP2 | antioxidant protein 2 | na |
| ARF2 | ADP-ribosylation factor 4 | na |
| ASPH | aspartyl/asparaginyl beta-hydroxylase | na |
| BKCO1 | transmembrane protein 32 | na |
| CARD11 | caspase recruitment domain protein 11 | na |
| CD3D | CD3e-associated protein | na |

| | | |
|---|---|---|
| CDH13 | Cadherin 13 | na |
| CISD1 | cells protein MDS029 | na |
| CLPP | putative atp-dependent clp protease proteolytic | na |
| CLTH | clathrin heavy chain 1 | na |
| CSNK1A1 | casein kinase I | na |
| CYP51 | cytochrome p450 51 | na |
| DHCR7 | 7 dehydrocholesterol reductase | na |
| DNAJC5 | DNA homologue subfamily c member 5 | na |
| DNASE1L1 | dnase I-like | na |
| EBP | emopamil binding protein | na |
| EERG28 | potential membrane protein C14orf1 | na |
| EPRS | bifunctional aminoacyl-trna synthase | na |
| FASN | fatty acid synthase | na |
| FLNB | beta filamin | na |
| GANAB | glucosidase II alpha-subunit | na |
| GCN1 | similar to yeast translation activator GCN1 | na |
| GLG1 | golgi apparatus protein 1 | na |
| HACD3 | HSPC121 | na |
| HCC1 | nuclear protein hcc-1 | na |
| HNRNPH2 | heterogeneous nuclear ribonucleoprotein H2 | na |
| HRAS | h-ras-1 | na |
| IER3 | homologue of mouse immediate early response 3 | na |
| ITGA2 | integrin alpha-2 | na |

| | | |
|---|---|---|
| ITIH5 | inter-alpha (globulin) inhibitor H5-like | na |
| KRT15 | cytokeratin 15 | na |
| LAMTOR3 | late endosomal/lysosomal mp1 interacting protein | na |
| LBR | lamin b receptor | na |
| LDLR | low-density lipoprotein receptor | na |
| LSS | lanosterol synthase | na |
| MAOA | amine oxidase a | na |
| MARCHF5 | ring finger protein 153 | na |
| MAVS | KIAA1271 | na |
| MYL2 | myosin regulatory light chain 2 | na |
| NECTIN1 | poliovirus receptor related protein 1 | na |
| NUP155 | nuclear pore complex protein nup155 | na |
| PAG1 | phosphoprotein associated with glycosphingolipidenriched | na |
| PDIA4 | protein disulfide isomerase a4 | na |
| PEX13 | peroxisomal membrane protein 20 | na |
| PFK6 | 6-phosphofructokinase | na |
| PSE1 | importin beta 3 | na |
| PSEN1 | presenilin 1 | na |
| RAB18 | rab18 | na |
| RAB33B | rab33b | na |
| RAB35 | rab35 | na |
| RAC1C | isoform Rac1c | na |
| RAC1C | ras-related C3 botulinum toxin substrate 1 | na |

| | | |
|---|---|---|
| RALA | Ras-related protein ral-a | na |
| RRAS | r-ras | na |
| RTN4 | reticulon 4 | na |
| SC2 | synaptic glycoprotein sc2 | na |
| SDH | succinate dehydrogenase | na |
| SLC39A10 | SLC39A10 protein | na |
| SPCS1 | microsomal signal peptidase 12kDa subunit | na |
| SPTB | beta spectrin | na |
| TENM4 | transmembrane protein 4 | na |
| TIMM50 | homologue of yeast Tim50 | na |
| TMEM33 | db83 | na |
| TPD52L2 | tumor protein d54 | na |
| TRAP1 | TRAP-1 | na |
| UQCR | ubiquinol cytochrome c reductase complex | na |
| VMA5 | vacuolar atp synthase subunit c | na |
| XPO1 | exportin 1 | na |

**Table S3.**
Enrichment analysis in Gene Ontology (GO) for differentially expressed genes in BAL samples from COVID-19 patients and asthma controls.

| GO term ID | Description | Fold-enriched | Biological Category |
|---|---|---|---|
| GO:1990928 | response to amino acid starvation | 3.75 | Apoptosis |
| GO:2001242 | regulation of intrinsic apoptotic signaling pathway | 3.51 | Apoptosis |
| GO:0038066 | p38MAPK cascade | 7.66 | Cellular signaling |
| GO:0060071 | Wnt signaling pathway, planar cell polarity pathway | 4.21 | Cellular signaling |
| GO:0048010 | vascular endothelial growth factor receptor signaling pathway | 3.7 | Cellular signaling |
| GO:0006890 | retrograde vesicle-mediated transport, Golgi to ER | 3.4 | ER/Golgi/Endosome |
| GO:1901998 | toxin transport | 3.67 | ER/Golgi/Endosome |
| GO:0006984 | ER-nucleus signaling pathway | 3.76 | ER/Golgi/Endosome |
| GO:0071985 | multivesicular body sorting pathway | 4.81 | ER/Golgi/Endosome |
| GO:0045056 | transcytosis | 4.86 | ER/Golgi/Endosome |
| GO:1904380 | endoplasmic reticulum mannose trimming | 5.11 | ER/Golgi/Endosome |
| GO:0061738 | late endosomal microautophagy | 5.37 | ER/Golgi/Endosome |
| GO:1903513 | endoplasmic reticulum to cytosol transport | 5.67 | ER/Golgi/Endosome |
| GO:0036258 | multivesicular body assembly | 5.79 | ER/Golgi/Endosome |
| GO:0045324 | late endosome to vacuole transport | 6.81 | ER/Golgi/Endosome |
| GO:1904903 | ESCRT III complex disassembly | 7.15 | ER/Golgi/Endosome |
| GO:0006907 | pinocytosis | 7.43 | ER/Golgi/Endosome |
| GO:0007042 | lysosomal lumen acidification | 7.66 | ER/Golgi/Endosome |
| GO:0006614 | SRP-dependent targeting to membrane | 9.25 | ER/Golgi/Endosome |
| GO:0016601 | Rac protein signal transduction | 3.89 | Immune |
| GO:0050860 | negative regulation of T cell receptor signaling pathway | 3.89 | Immune |
| GO:0042535 | positive regulation of tumor necrosis factor biosynthetic process | 4.08 | Immune |
| GO:0098761 | cellular response to interleukin-7 | 4.08 | Immune |

| | | | |
|---|---|---|---|
| GO:0090382 | phagosome maturation | 4.27 | Immune |
| GO:0048143 | astrocyte activation | 4.3 | Immune |
| GO:0010759 | positive regulation of macrophage chemotaxis | 4.77 | Immune |
| GO:0038061 | NIK/NF-kappaB signaling | 4.86 | Immune |
| GO:0019882 | antigen processing and presentation | 4.92 | Immune |
| GO:0097178 | ruffle assembly | 5.11 | Immune |
| GO:0045657 | positive regulation of monocyte differentiation | 5.57 | Immune |
| GO:0032930 | positive regulation of superoxide anion generation | 6.13 | Immune |
| GO:0035722 | interleukin-12-mediated signaling pathway | 6.22 | Immune |
| GO:0002479 | antigen processing via MHC class I, TAP-dependent | 6.99 | Immune |
| GO:0019076 | viral release from host cell | 8.51 | Immune |
| GO:2001188 | regulation of T cell activation via MHC - antigen presenting cell | 8.51 | Immune |
| GO:0002399 | MHC class II protein complex assembly | 10.21 | Immune |
| GO:0071226 | cellular response to molecule of fungal origin | 10.21 | Immune |
| GO:0071826 | ribonucleoprotein complex subunit organization | 3.46 | LipidRaft |
| GO:0034381 | plasma lipoprotein particle clearance | 3.57 | LipidRaft |
| GO:1990776 | response to angiotensin | 4.16 | LipidRaft |
| GO:0046479 | glycosphingolipid catabolic process | 5.5 | LipidRaft |
| GO:0055094 | response to lipoprotein particle | 4.16 | Lipoprotein |
| GO:0071402 | cellular response to lipoprotein particle stimulus | 4.58 | Lipoprotein |
| GO:0010288 | response to lead ion | 3.68 | Metal ion |
| GO:0033572 | transferrin transport | 5.11 | Metal ion |
| GO:0000715 | nucleotide-excision repair, DNA damage recognition | 4.44 | Nuclease |
| GO:0032069 | regulation of nuclease activity | 6.03 | Nuclease |
| GO:0032075 | positive regulation of nuclease activity | 8.93 | Nuclease |
| GO:0000084 | mitotic S phase | 5.34 | Proliferation |

| | | | |
|---|---|---|---|
| GO:1902410 | mitotic cytokinetic process | 5.83 | Proliferation |
| GO:0006457 | protein folding | 3.42 | Protein Folding |
| GO:0042982 | amyloid precursor protein metabolic process | 4.3 | Protein Folding |
| GO:1903332 | regulation of protein folding | 5.11 | Protein Folding |
| GO:0097242 | beta-amyloid clearance | 5.77 | Protein Folding |
| GO:0042987 | amyloid precursor protein catabolic process | 5.96 | Protein Folding |
| GO:1900223 | positive regulation of beta-amyloid clearance | 7.29 | Protein Folding |
| GO:0018279 | protein N-linked glycosylation via asparagine | 4.02 | Protein Metabolism |
| GO:0006517 | protein deglycosylation | 4.08 | Protein Metabolism |
| GO:0051444 | negative regulation of ubiquitin-protein transferase activity | 4.2 | Protein Metabolism |
| GO:0006521 | regulation of cellular amino acid metabolic process | 6.06 | Protein Metabolism |
| GO:0006413 | translational initiation | 7.71 | Protein Metabolism |
| GO:0006091 | generation of precursor metabolites and energy | 3.48 | Respiration |
| GO:2000191 | regulation of fatty acid transport | 3.65 | Respiration |
| GO:0098780 | response to mitochondrial depolarisation | 4.08 | Respiration |
| GO:0045730 | respiratory burst | 5.35 | Respiration |
| GO:0019646 | aerobic electron transport chain | 6.13 | Respiration |
| GO:0019322 | pentose biosynthetic process | 10.21 | Respiration |
| GO:0043620 | regulation of DNA-templated transcription in response to stress | 4.7 | RNA metabolism |
| GO:0061418 | regulation of transcription in response to hypoxia | 5.89 | RNA metabolism |
| GO:1902415 | regulation of mRNA binding | 6.13 | RNA metabolism |
| GO:1904874 | positive regulation of telomerase RNA localization to Cajal body | 6.81 | RNA metabolism |
| GO:0000184 | mRNA catabolic process, nonsense-mediated decay | 7.66 | RNA metabolism |
| GO:0021762 | substantia nigra development/14-3-3 family of proteins | 4.22 | Serine phosphorylation |
| GO:0070262 | peptidyl-serine dephosphorylation | 4.77 | Serine phosphorylation |

**Table S4.**

Classification of conservative substitutions considered in this study. Amino acids in brackets can be considered to be part of a group after structural analysis.

| Group | Amino acid |
|---|---|
| Aromatic | Y, W, [H], [F] |
| Non-polar | A, I, V, L, M, [F], [P] |
| Polar, hydrogen bond interacting | S, T, [C] |
| Amidic, hydrogen bond interacting | N, Q |
| Acidic | D, E |
| Basic | R, H, K |
| Glycine | G |

**Table S5.** Average root mean-square deviation (RMSD) of models built with the workflow relative to experimentally determined structures (*Workflow*). In parenthesis, the region modeled is defined. *Exp.*: Resolution of experimentally determined structure; *Other*: average RMSD of models generated with C-I-TASSER pipeline (*41*). Structure alignment was conducted using Lovoalign (*42*).

| Protein | Exp. / Å | Workflow / Å | Other / Å |
|---|---|---|---|
| nsp5 (full) | 2.0 | 1.5 | 2.8 |
| nsp3 (205-374) | 1.5 | 1.5 | - |
| nsp15 (27-332) | 2.2 | 0.7 | 0.5 |

### Appendix: Structural Analysis of the SARS-CoV-2 Proteome

#### Nonstructural protein 1 (nsp1)

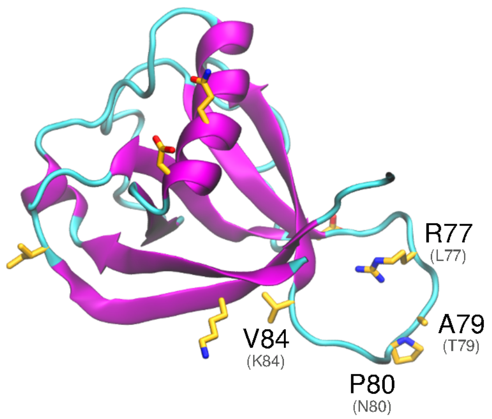


**Fig. S8.** Predicted structure of SARS-CoV-2. Non-conservative substitutions relative to SARS-CoV-1 are depicted in *orange*.

The nonstructural protein 1 (nsp1) is the first nonstructural protein in the ORF1a/ORF1ab gene. Nsp1 is found within α and β coronaviruses (Fig. S8). Experiments *in vitro* suggest that SARS-CoV-1 nsp1 disrupts the host interferon defense response by potentially affecting the downstream defense signaling (*43*–*45*). The nsp1 protein binds the 40S Ribosomal subunit, which has been associated with degradation of host mRNA and suppression of host mRNA translation, leaving the viral RNA unaffected (*45*, *46*). The resultant complex cleaves the 5' UTR of host mRNAs, inhibiting translation.

#### Nonstructural protein 2 (nsp2)

Positive RNA viruses create viroplasms, subcellular compartments that are thought to protect the viral machinery against host-defense strategies, and even facilitate replication (*47*). They often contain double stranded RNA, as well as the replication complex of the virus. For SARS-CoV-1, the replicase proteins assemble at the cytosol side of the ER, after which invagination occurs. There is no evidence of a pore maintaining the connection of the invaginated space to the cytosol, thus no maintained sphererule has been observed, though vesicles do form. A network of double-membrane-vesicles (DMVs) is eventually formed, anchored to the ER by a convoluted-membrane (CM) compartment. Replication appears to happen in the CM, while DMVs directly connected to the CM contain replicase proteins and viral proteins. Those DMVs not directly connected to CM form vesicle packets containing viral genomes (*47*, *48*).

In Hagemeijer *et al.* (*49*), it was shown that the nonstructural protein 2 (nsp2) localized to the membranes of CM and DMV, and it is exposed to the cytoplasm. It appears that during formation of the replication complex, nsp2 is recruited, but once formed no further exchange of nsp2 occurs. Mutants of SARS-CoV-1 with the deleted nsp2 coding sequence are still capable of viral replication, although with decreased growth (*50*) The specific role of nsp2 has not been established; it may be that nsp2 assists other viral proteins in performing their function, such as regulating the autophagy defense response or promoting mitochondrial dysfunction, thereby helping viral replication or effecting disease severity. It has been shown that nsp2 interacts directly with prohibitin 1 (PHB1) and PHB2 (*51*) host proteins with a wide variety of functions. Kathiria *et al*. (*52*) have shown that PHB knockdown resulted in increased ROS, mitochondrial depolarization, and induced autophagy. In (*53*) several phenotypes of PHB are reviewed, including the role of PHB in mitochondrial stability. PHB has also been implicated in inflammatory response in both the lung and gut (*53*, *54*). In von Brunn *et al*. (*55*), it was shown that nsp2 displayed co-immunoprecipitation (CoIP) interaction with N-terminal side of nsp3, nsp6, nsp8, nsp11, nsp16 and ORF3a, and co-localization with nsp8 and nsp3, where nsp8 almost always co-localized with the microtubule protein, LC3, an autophagy marker protein (*56*). These results suggest that nsp2 may be involved in mitochondrial dysfunction through PHB1 and 2, and association with LC3 may be worth investigating. Another potential for nsp2's association with mitochondrial dysfunction may lie with its CoIP interaction with ORF3a, as the SARS protein ORF3a is known to disrupt mitochondrial stability (*57*).

A

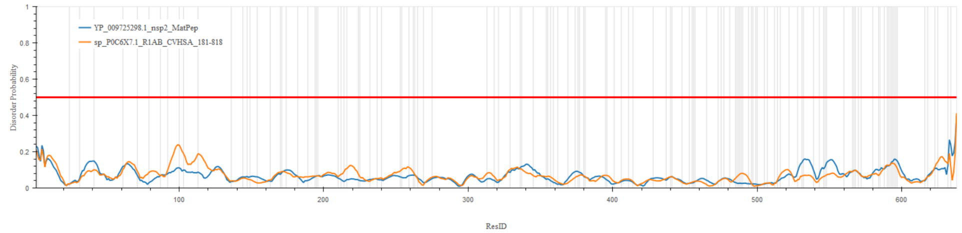


B

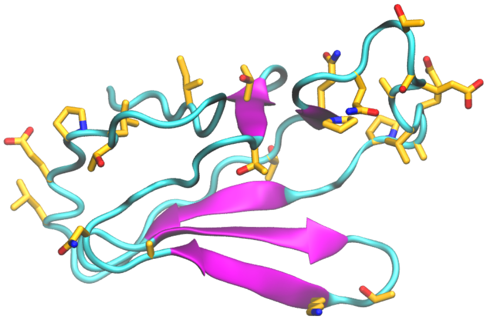

**Fig. S9.** A) Profile of disordered/structured regions of SARS-CoV-2 nsp2, predicted using DisEMBL (*58*), using the *hot loop* definition. The red line indicates the 50% threshold for disorder, comparing nsp2 SARS-CoV-1 (*yellow* line), to nps2 SARS-CoV-2 (*blue* line). The grey vertical lines indicate the location of the non-conserved substitutions in SARS-CoV-1 nsp2 relative to SARS-CoV-2 nsp2. B) C-terminal domain of SARS-CoV-2 nsp2. Non-conservative substitutions relative to SARS-CoV-1 are depicted in *orange*.

*Structural analysis and comparison with SARS-CoV-1 nsp2* - Structure information about nsp2 is scarce. SARS-CoV-2 nsp2 is 68% identical to SARS-CoV-1 nsp2. Among the substitutions, 130 are non-conservative and highly concentrated in the C-terminal end of the protein (Fig. S9A). An *ab initio* model was generated for this region (556-633, Fig. S9B). Fewer mutations appear at the N-terminal side, and a highly conserved region appears in the middle of the protein sequence.

**Nonstructural protein 3 (nsp3) - Papain-like proteinase**

The nonstructural protein 3 (nsp3) acts as a phosphatase and its catalytic domain is conserved, sharing homology with the Ymx7 protein in yeast, AF1521 in the archaea (*Archeoglobus fulgidus*), and Er58 in *E. coli*. (*59*) The protein comprises at least six domains: 1) an N-terminal ubiquitin-like (Ubl1) domain followed by glu-rich acidic domain; 2) an X domain with a predicted Appr-100-p processing activity; 3) a SUD domain (SARS-specific unique domain); 4) a peptidase C-16 domain that contains the papain-like protease (abbreviated PLnc); 5) a transmembrane domain; and 6) the Y domain.

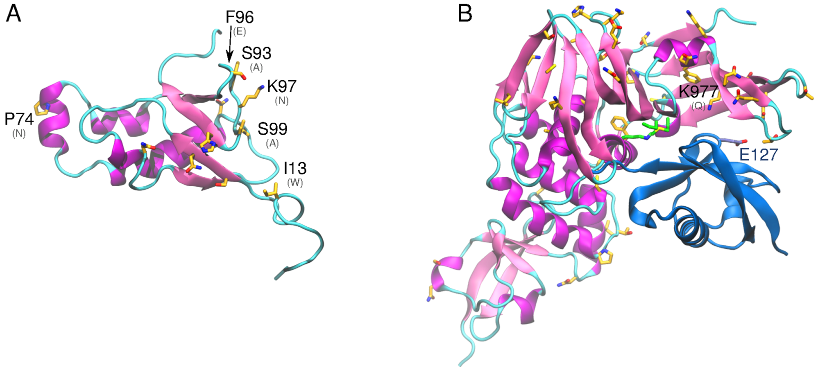


**Fig. S10.** A) Predicted structure of N-terminal ubiquitin-like domain, and (B) papain-like protease domain of SARS-CoV-2 nsp3 bound to human interferon-stimulated gene product 15 (ISG15). Non-conservative substitutions relative to SARS-CoV-1 are depicted in *orange*. Key residues identified to bind ISG15 are conserved (in *green*). The complex with ISG15 was built based on PDB id 5tl6.

*Structural analysis and comparison with SARS-CoV-1 nsp3* - Ub1 binds the nucleocapsid (N) protein and the interaction involves acidic residues of Ubl1 helix α2 and the serine and arginine rich region of the N protein (*60*). Ub1 also binds to the 5' untranslated region of coronavirus RNA. Ub1 has significant structural homology with Ras effector proteins, and it is thought that Ub1 may interact with and modulate the activity of Ras in the host. This interaction potentially affects growth and cell cycle cascades and could be a potential mechanism for cell mortality.(*61*) Significantly, there are differences in residue identity between SARS-CoV-1 and SARS-CoV-2 in residues 41 to 63, 83 to 87, 88 to 94, 95 to 98 of the Ub1 region (Fig. S10A). These regions correspond to structural homology with Ras effectors and these residue changes may affect interactions with host Ras and contribute to pathogenicity divergence (*61*).

The glu-rich acidic region is also known as the hypervariable region (HVR). It is intrinsically disordered and its interaction partners are not entirely clear. One study by yeast-2-hybrid demonstrates interaction with nsp6, but a GST pull-down assay also revealed interactions with nsp8, nsp 9, and 3 regions of nsp3 itself (nucleic-acid binding domain, betacoronavirus-specific marker domain, and transmembrane region 1) (*60*). This region is significantly elongated, relative to the SARS-CoV-1 counterpart, with 16 additional amino acids, including several potential sites of glycosylation.

The X domain binds ADP-ribose, the structure of this complex was recently solved for SARS-CoV-2 (PDB id 6w02), and there are several studies and structures available in the Protein Data Bank of homolog complexes (PDB id: 5hol, 5dus, and 2fav). ADP-ribosylation is a type of post-translational modification (PTM), which may be implicated in inhibiting the immune response of the host through PTM of proteins related to expression of interleukin-6 and interferon-beta (*60*). A GST-pulldown assay showed interaction of the X domain of nsp3 with nsp12. Several non-conservative substitutions occur in this domain relative to SARS-CoV-1, but those are located in superficial regions and do not significantly affect protein conformation (RMSD 1.5 Å, relative to

2fav). The papain-like protease domain (papain-like protease 2; PL2pro) (*60*) contains two ubiquitin binding sites. It is implicated in suppression of the host immune response, but its targets and, more generally, which immune-related signal transduction cascades are affected is unclear. It is implicated in inhibiting components of NF-κB, interferon-beta, and p53. In a structural study, $PL2^{pro}$ was found to bind ubiquitin-like interferon-stimulated gene product 15 (ISG15) (*62*), the latter an important post-translational modification of host proteins, including cytokines like interferon. It is believed that cleaving these posttranslational modifications of cytokine proteins by $PL2^{pro}$ disrupts the host immune response (*62*). Importantly, ISG15 has significant interspecies variability, potentially contributing to its very different virulence patterns among species. This region is mostly conserved within SARS-CoV-1 and SARS-CoV-2, including residues that were identified as critical for the interaction with ISG15, namely $Arg^{911}$, $Met^{953}$ and $Pro^{992}$. However, the substitution $Gln^{977}Lys$ likely intensifies the interaction with ISG15 by forming a salt bridge with $Glu^{127}$, suggesting an important mechanism for variable virulence (Fig. S10B).

**Nonstructural protein 4 (nsp4)**

Working in coordination with nsp3 and nonstructural protein 6 (nsp6), the nonstructural protein 4 (nsp4) of SARS-CoV-1 is essential for membrane rearrangements during viral replication (*63*, *64*). Data suggests that coexpression of nsp3 with nsp4 results in host membrane rearrangement and the formation of double membrane vesicles and convoluted membranes (*65*). Nsp4 in mouse hepatitis virus, another coronavirus, is a glycosylated protein and is demonstrated to be involved in virally induced membrane rearrangement, replication complex assembly, and assembly of double membrane vesicles (*66–70*). Aberrant double membrane vesicles and impaired RNA replication can be observed when mouse hepatitis virus nsp4 is lacking a glycosylation site (*67*, *71*). Prevention of interaction between nsp4 and nsp3 eliminated viral replication (*63*).

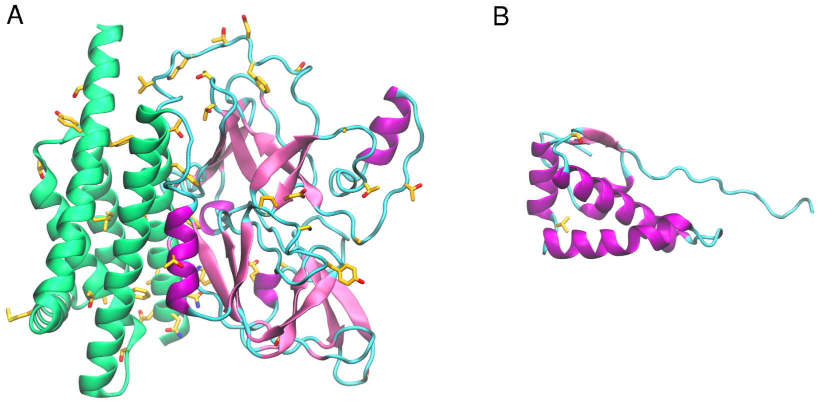


**Fig. S11**. A) *Ab initio* model of region 1-402 of SARS-CoV-2 nsp4. B) Predicted structure of the conserved C-terminal domain of SARS-CoV-2. Transmembrane regions are shown in *green*, soluble domains in *pink* and non-conservative substitutions relative to SARS-CoV-1 nsp4 in *orange*.

*Structural analysis and comparison with SARS-CoV-1 nsp4* - Nsp4 of SARS-CoV-1 is a nonstructural protein derived from the replicase polyprotein and is thought to be a tetra spanning transmembrane protein (*68*). Disruption in glycosylation sites within the luminal loop between transmembrane domains 1 and 2 give rise to aberrant double membrane vesicles (*67*). Key amino acid residues His$^{120}$ and Phe$^{121}$ of nsp4 of SARS-CoV-1, which are conserved in SARS-CoV-2, are essential for interaction and binding of nsp4 to nsp3 as well as viral propagation (*63*).

Structural information about nsp4 is scarce. A low-resolution model was generated for the full nsp4 using the *ab initio* protocol (Fig. S11A). The soluble C-terminal domain is solved for a homologue (3vcb, MHV nsp4), enabling the generation of a higher resolution model for this region using the fragment-based modeling protocol (Fig. S11B). Despite the likely inaccurate orientation of domains in the full protein model, the predicted secondary structures are consistent with the expected positions of the transmembrane and soluble domains. This model is a useful resource for general structural analysis.

The sequence of SARS-CoV-2 nsp4 is 80% identical to SARS-CoV-1 nsp4. Among the non-conservative substitutions, the predicted model reveals the substitution of four cysteines in the predicted TM helices by the bulky amino acids, Trp$^{6}$, Phe$^{7}$, Phe$^{24}$ and Phe$^{390}$. These substitutions may affect the arrangement and packing of transmembrane segments. Several substitutions are predicted to occur on the surface of the modeled soluble region spanning amino acids 33-275. The cytoplasmic C-terminal domain is highly conserved.

**Nonstructural protein 5 (nsp5 or 3CL$^{pro}$) - 3C-like proteinase**

The nonstructural protein 5 (nsp5, also known as 3CL$^{pro}$), is the main protease of the coronavirus genome, exhibiting a main role in the cleavage of the polyproteins translated from the viral RNA (*72–74*). This protein is highly conserved relative to SARS-CoV-1 (96% identity) and among RNA+ viruses (Nidovirales) in general, making it an attractive target for pan-antiviral drugs (*75–77*). In addition, it has been shown that loss of nsp3 and nsp10 substantially reduces 3CL$^{pro}$ activity (*78*,*79*) and therefore therapeutics that target these proteins could indirectly inhibit 3CL$^{pro}$.

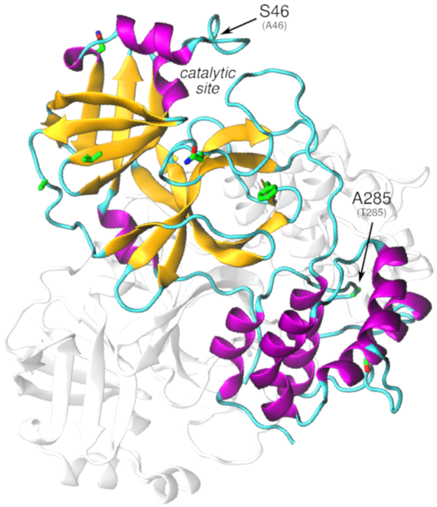


**Fig. S12.** SARS-CoV-2 nsp5 dimer modeled using the crystal structure of the monomer and PDB 3m3v as dimer template. Non-conservative substitutions relative to SARS-CoV-1 nsp5 are depicted in *green*.

*Structural analysis and comparison with SARS-CoV-1 nsp5* - Studies with SARS-CoV-1 show that dimerization is essential to stabilize the productive conformation of 3CL$^{pro}$ catalytic site. The recently solved structure of 3CLpro of SARS-CoV-2 (PDB id: 6y2e) confirms the dimer as its biological state (Fig. S12). The dimer interface is highly conserved within SARS-CoV-1 and -

CoV-2, as well as other residues that indirectly affect dimerization, such as Ser$^{144}$, Ser $^{147}$ and Asn$^{28}$ (*80*). However, a relevant substitution Thr$^{285}$Ala is found on the interface. Based on previous studies with SARS-CoV-1 3CL$^{pro}$, this replacement is thought to favor the hydrophobic pack within monomers, and it was recently associated with a slightly higher catalytic efficiency of SARS-CoV-2 compared to SARS-CoV-2 (*81*). The analysis of the evolutionary tree from the aligned sequences of coronavirus from all available species reveals that alanine at site 285 defines the SARS-CoV-2 clade and three bat coronaviruses from mainland China (Fig. S13). In contrast, the many of the beta coronaviruses that infect mammals have a cysteine at this location. Given the proximity with the cysteine in the opposite monomer, it is possible that a disulfide bridge is formed in these proteases, which may result in a more tightly bound dimer and increased catalytic efficiency. Further exploration of this site is warranted.

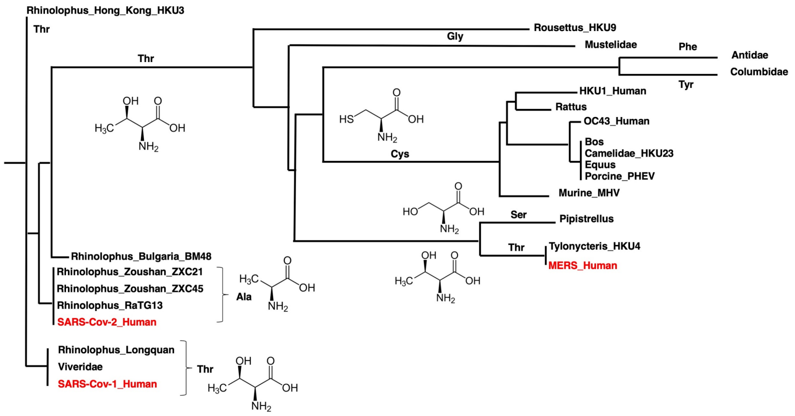


**Fig. S13.** Phylogenetic tree of all known beta coronaviruses using the full protein sequence for 3CL$^{pro}$. *Red* text indicates viruses of concern for human health. The amino acid changes for the site at 285, which may affect dimerization are shown along branches. An alanine at this site defines the SARS-CoV-2 clade with the horseshoe bats from mainland China.

There are 7 non-conservative amino acid substitutions between SARS-CoV-1 and SARS-CoV-2, besides Thr$^{285}$Ala. Most of those are located in regions that are not clearly associated with protease function, except by Ala$^{46}$Ser, positioned at the catalytic cleft entrance, which may affect substrate affinity/selectivity. Moreover, the phosphorylation of serines in 3CL$^{pro}$ of rotaviruses was shown to be essential for protease activity. The substitutions Ala$^{46}$Ser, Ser$^{65}$Asn, Ser$^{94}$Ala, Ala$^{267}$Ser and Thr$^{285}$Ala may also affect the phosphorylation pattern of 3CL$^{pro}$ in SARS-CoV-2.

**Nonstructural protein 6 (nsp6)**

Upon infection, corona viruses create signature membrane rearrangements (e.g. DMVs) so that the viral RNA replication complex may anchor to host machinery (*82*). Nsp6, along with nsp3 and nsp4, plays a critical role in this membrane rearrangement (*64*). In SARS-CoV-1 nsp6 is known to activate autophagy (*83*) by inducing perinuclear vesicles localized around the microtubule organization center (*64*).

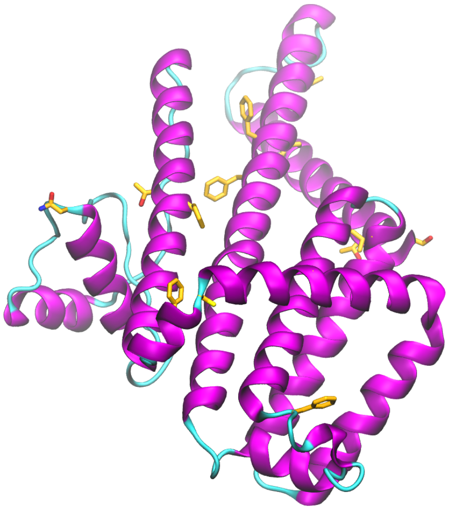

**Fig. S14.** *Ab initio* predicted structure of SARS-CoV-2 nsp6. Non-conservative substitutions relative to SARS-CoV-1 nsp6 are depicted in *orange*.

*Structural analysis and comparison with SARS-CoV-1 nsp6* - SARS-CoV-2 nsp6 is 87% identical to SARS-CoV-1 nsp6. Non-conservative mutations are located on the protein surface (Fig. S14).

**Nonstructural proteins 7, 8 and 12 (nsp7, nsp8, nsp12) - Replication complex**

After infection the virus assembles a multi-subunit RNA-synthesis complex consisting of several nonstructural proteins (nsps), namely nsp7, 8, 9, 10, and 12.(*84*) This complex is responsible for the replication and transcription of the viral genome. The complex of nsp12, 7, and 8 (total 160 kDa) constitutes the minimum set of nsps required for nucleotide polymerization.(*85*) Nsp12 encodes the RNA-dependent RNA polymerase (RdRp) domain. Nsp7 and nsp8 are nsp12 cofactors. Nsp7 forms a hexadecameric complex with nsp8 and together these may act as a processivity clamp for the RNA polymerase.(*86*) In the replication complex there are two well defined regions of nsp8: nsp8(I), forming the heterodimer with nsp7, and nsp8(II), binding to nsp12.(*85*)

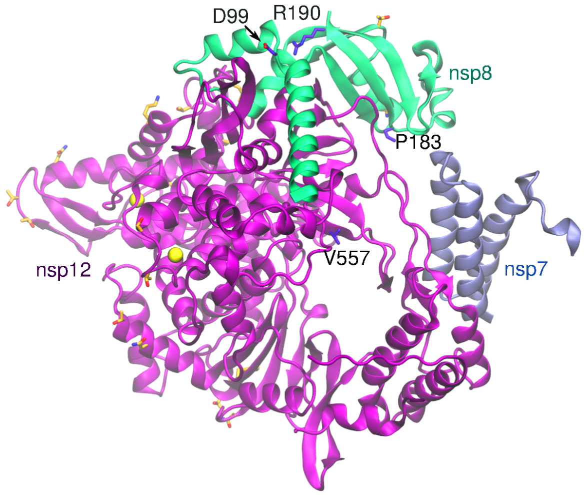


**Fig. S15.** Local modeling-derived structure of SARS-CoV-2 nsp7-nsp8-nsp12 complex. Non-conservative substitutions are depicted in *orange*. Some key functional residues identified are depicted in *purple*. Zinc ions are represented as *yellow* spheres. PDB 6nur was used as the template to assemble the complex.

*Structural analysis and comparison with SARS-CoV-1 nsp7-nsp8-nsp12 complex*- A study conducted by Kirchdoerfer and Ward(*85*) showed that the polymerase domain of SARS-CoV-1 nsp12 **(**a.a 398-919**)** consists of the fingers domain (a.a 398–581, 628–687), palm domain (a.a. 582–627, 688–815) and a thumb domain (a.a. 816–919). The Nidovirus-unique N-terminal extension (aka NiRAN) is situated between a.a. 1–397. There are two metal (Zn) binding sites in nsp12: The first is in the NiRAN extension and is coordinated by residues $His^{295}$, $Cys^{301}$, $Cys^{306}$, and $Cys^{310}$. The second site is in the fingers domain and is coordinated by $Cys^{487}$, $His^{642}$,$Cys^{645}$, and $Cys^{646}$. The binding site for the nsp7-nsp8 heterodimer overlaps with the conserved regions of polymerase functional domains (fingers (a.a 398–581, 628–687) and thumb domains (a.a. 816–919). The binding site between nsp8(II) and nsp12 is on the N-terminal region (77–126) of nsp8(II).
Several studies have pointed to functional residues in nsp7, 8, and 12 sequences. The study conducted by Lehmann et al.(*87*), shows conserved sequence motifs in the NiRAN domain named $A_N$, $B_N$, and $C_N$. These motifs are conserved throughout all members of the order *Nidovirales*, of which CoVs are members.(*87*) Another example is covalent modification of the $Lys^{73}$ residue in nsp12 (using GTP or UTP) that reduced viral growth and recovery in the equine arterivirus.(*87*) *In vitro* polymerase activity assays further enabled the identification of other key functional sites, $Lys^{7}$, $His^{36}$, and $Asn^{37}$, which, being replaced by alanine, are associated with decreased polymerase activity.(*88*) Four other mutations in nsp8 were detrimental to polymerase activity namely, $Pro^{183}Ala$, $Asp^{99}Ala$, $Pro^{116}Ala$, and $Arg^{190}Ala$. These mutations are associated with a defective fold of nsp8 and disruption of nsp8(II)-nsp12 binding.(*88*)
The nucleoside analog GS-5734 is capable of impairing CoV RNA synthesis by targeting the viral RNA synthesis machinery. The side chain in a motif of the fingers domain in nsp12 is involved in GS-5734 interaction through $Val^{557}$ residue.(*89*, *90*)
All the mentioned residues are fully conserved in SARS-CoV-2. All non-conservative substitutions are located in the complex surface, mostly in nsp12 (Fig. S15). Several of these substitutions involve residue that are potential sites for post-translational modifications (e.g., cysteines, serines, threonines, asparagines and tyrosines), indicating a variation in post-translational patterns relative to the SARS-CoV-1 RNA polymerase complex.

**Nonstructural protein 9 (nsp9)**
The nonstructural protein 9 (nsp9) is cleaved from the viral polyproteins by the $3CL^{pro}$ protease.(*84*, *91*) SARS-CoV-1 nsp9 is able to form dimers (Fig. S16) that bind either ssDNA and ssRNA(*92*) and is thought to protect the coronavirus genome from degradation during replication.(*91*, *93*) Deletion of nsp9 in the mouse β-coronavirus (hepatitis virus , MHV), impairs viral RNA synthesis and viral infection, where dimerization is particularly important.(*94*, *95*)

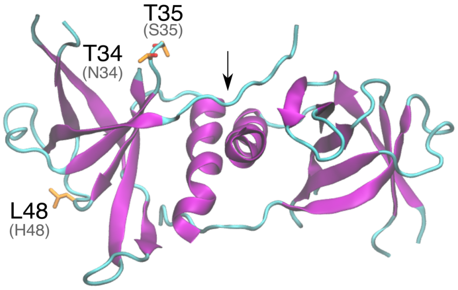


**Fig. S16.** Cartoon representation of SARS-CoV-2 nsp9 dimer. Arrow indicates dimer interface and mutations are highlighted in orange**.**

*Structural analysis and comparison with SARS-CoV-1 nsp9* - Overall, the structure of nsp9 is very well conserved among β-coronaviruses.(*91*, *96*) In SARS-CoV-1, nsp9 consists of a small globular protein (113 amino acid residues) with seven antiparallel β-sheets and one α-helix.(*91*) Parallel dimers occur between two nsp9 monomers interacting in their C-terminal α-helixes and their N-fingers.(*96*) In SARS-CoV-1, $Gly^{100}$ and $Gly^{104}$ residues are shown to be in the core of the dimer interface.(*91*) Mutation of these conserved glycines in the nsp9 α-helix impairs ssDNA binding, suggesting that dimerization is essential for nucleic acid binding.(*96*) Site-specific mutagenesis studies in the alpha-coronavirus Porcine Epidemic Diarrhea (PEDV) nsp9 identified other key residues for dimerization. The substitutions $Lys^{10}$Ala, $Arg^{68}$Ala, $Lys^{69}$Ala and $Arg^{106}$Ala decreases binding affinity to form the dimer 7·2-fold relative to wild-type nsp9, while $Tyr^{82}$Ala enhance dimer stability, by a 8·0-fold increase in binding affinity.(*96*) Additionally, Zeng et al. (2018), studied Porcine Epidemic Diarrhea Virus (PEDV) nsp9 residues involved in ssDNA binding activity and they identified five mutations that affected this process, namely, $Lys^{10}$Ala, $Arg^{68}$Ala, $Lys^{69}$Ala, $Arg^{106}$Ala and $Tyr^{82}$Ala. The equivalents in SARS-CoV-2 identified from a sequence alignment are $Arg^{10}$Ala, $Arg^{74}$Ala, $Arg^{11}$1Ala, and $Tyr^{87}$Ala, which are conserved in SARS-CoV-1.

Because SARS-CoV-1 and SARS-CoV-2 sequences are highly conserved (sequence identity is 97%), much structural information from SARS-CoV-1 is transferable to SARS-CoV-2. The three substitutions in SARS-CoV-2 nsp2 relative to SARS-CoV-1, $Asn^{34}$Thr, $Ser^{35}$Thr and $His^{48}$Leu, are located on the nsp9 surface, distant from the dimerization region (Fig. S16). Particularly, substitution $Asn^{34}$Thr can result in an additional phosphorylation site, but phosphorylation of nsp9 has not yet been reported. As $Thr^{34}$ and $Thr^{35}$ are close to a potential ubiquitination site ($Lys^{36}$), we hypothesize that their phosphorylation may prevent nsp9 ubiquitination.

**Nonstructural protein 10 (nsp10)**

Nsp10 is one of sixteen nonstructural proteins of the SARS-CoV-2 proteome. It interacts with nsp14 and nsp16 to perform 3’-5’ exoribonuclease and 2'-O-methyltransferase activities, respectively.(*97*, *98*) Disturbance in the interaction between nsp10 and nsp16 has been shown to be crippling to the virus, although still functional.(*99*) Nsp14, nsp16, and nsp10 have been associated with the coronavirus wide replication and transcription complex.(*100*, *101*) Crystal structures of SARS-CoV-1 nsp10-nsp16 have been reported.(*102*)

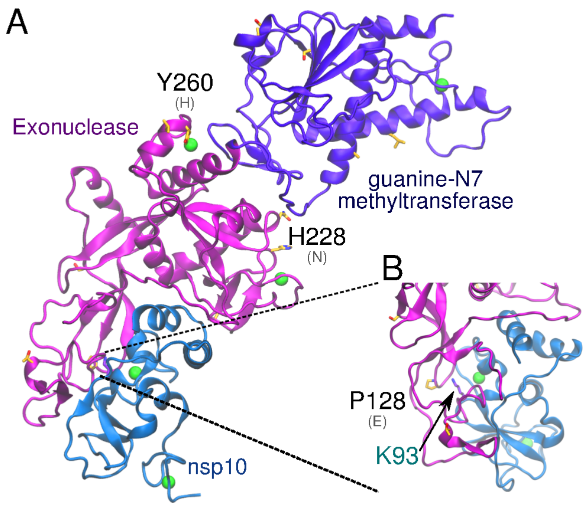


**Fig. S17.** A) Local modeling-derived structure of SARS-CoV-2 nsp10-nsp14 complex. B) Closer view of residues $Pro^{128}$ and $Lys^{93}$ in nsp14 and nsp10 respectively. Non-conservative substitutions relative to SARS-CoV-1 nsp10 and nsp14 are depicted in *orange*. Zinc ions are represented as *green* spheres. PDB 5c8s was used as the template to assemble the complex.

*Structural analysis and comparison with SARS-CoV-1 nsp9* - SARS-CoV-1 and SARS-CoV-2 nsp10 proteins are highly conserved (identity 97%), with no non-conservative substitutions between them. These proteins are composed of 139 a.a. residues and fold forming two zinc fingers.(*99*) This tight interaction with nsp14, involving multiple hydrogen bonds, salt bridges and hydrophobic packing, suggests that nsp10 might be necessary to maintain the integrity of nsp14 exoribonuclease domain.(*103*, *104*)

**Nonstructural protein 11 (nsp11)**

Nsp11 is a short protein only 13 a.a. in length and is translated from both polyproteins ORF1a and ORF1ab. In SARS-CoV-1 nsp11 has been implicated in RNA synthesis.(*105*) SARS-CoV-2 nsp11 has 85% sequence identity to SARS-CoV-1 nsp11, with two substitutions between them ($Ser^5$Gln and $Thr^6$Ser).

**Nonstructural protein 13 (nsp13) - helicase**

The nonstructural protein 13 (nsp13), encoded from ORF1ab as part of the pp1ab polyprotein, is involved in a number of functions, such as NTPase, dNTPase, RTpase, RNA helicase, and DNA helicase activity.(*25*) It also interacts with nsp12, which is thought to enhance its helicase activity.(*25*, *106*)

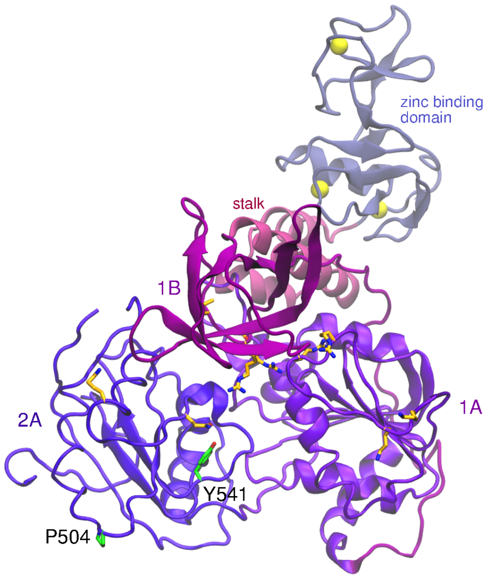


**Fig. S18**. Predicted structure of SARS-CoV-2 nsp13. Non-conservative substitutions relative to SARS-CoV-1 nsp13 are depicted in orange. Mutations observed in SARS-CoV-2 nsp13 are depicted in green. Zinc ions are represented as yellow spheres.

*Structural analysis and comparison with SARS-CoV-1 nsp13* - The helicase nsp13 contains 5 domains, an N-terminal zinc-binding domain, stalk domain, RecA-like domain 1A, 1B, and RecA-like domain 2A, all together consisting of 603 residues. It forms a triangular structure with the 2A and 1A as the base.(*24*)
The zinc-binding domain consists of conserved cysteine residues, which have shown to be essential for the various enzymatic activities of nsp13.(*107*) The stalk domain links the zinc-binding domain to the rest of the helicase, where the stalk domain has been argued to serve as an essential signal transduction mechanism.(*24*) A single mutation in the stalk domain of SARS-CoV-1 nsp13, $Arg^{132}$Pro, resulted in a dramatic decrease in viral infectivity *in vitro*.(*108*) The essential nature of nsp13 in the viral replication cycle, and its multifunctional nature has made it an attractive target for vaccine research.(*109*)(*110*, *111*) SARS-CoV-2 nsp13 (Fig. S18) is fully conserved relative to SARS-CoV-1.

**Nonstructural protein 14 (nsp14) - 3' to 5'- exonuclease**
The nonstructural protein nsp14 plays a critical role in coronaviruses replication and transcription. The exonuclease domain of nsp14 is imperative for replication fidelity within RNA viruses and has been shown to function as a proofreading exoribonuclease. Nsp14 associates with other nonstructural proteins from ORF1a and ORF1ab, including nsp7, nsp8, nsp12, and nsp10.(*88*, *103*, *112–114*) The association with nsp10 provides the ability to excise mismatched nucleotides(*104*), and disruption of this heterodimer was shown to decrease replication fidelity.(*115*)
*Structural analysis and comparison with SARS-CoV-1 nsp14* - Similar to SARS-CoV-1, SARS-CoV-2 nsp14 protein contains two catalytic domains: (1) The N-terminal exonuclease domain and (2) the C-terminal guanine-N7 methyltransferase domain. These domains, N-terminal and C-terminal, span a.a. sequences 1-287 and 288-527, respectively. The exonuclease domain also has conserved residues Asp, Glu, Asp and Asp (DEDD) that are key indicators of the DEDDh superfamily of DNA and RNA exonucleases.(*84*, *116*) The guanine-N7 methyltransferase domain

has been shown to be necessary for viral mRNA capping(*103*), hiding viral mRNA from host mRNA degradation machinery.

Nsp14 contains two zinc fingers within the N-terminal domain. The first is composed of Cys$^{207}$, Cys$^{210}$, Cys$^{226}$ and His$^{229}$, while the second is composed of His$^{257}$, Cys$^{261}$, His$^{264}$, and Cys$^{279}$(*103*)**.** Ala$^{1}$-Arg$^{76}$ and Ala$^{119}$-Asp$^{145}$ of nsp14 interact with nsp10. The third zinc finger of nsp14 is within the C-terminal domain and is formed by Cys$^{452}$, Cys$^{477}$, Cys$^{484}$, and His$^{487}$.(*103*)

SARS-CoV-2 nsp14 is 73% identical to the SARS-CoV-1 counterpart. Among the 14 non-conservative substitutions, Glu$^{128}$Pro is the only substitution located close to the interface with nsp10 (Fig. S17B). The proximity with the positively charged Lys$^{93}$ of nsp10 in the heterodimer model suggests that this substitution may slightly affect binding of nsp10-nsp14 (Fig. S17B). The substitutions His$^{260}$Tyr and Asn$^{228}$His may also be relevant as they are located in close proximity to the zinc fingers in the N-terminal exonuclease domain (Fig. S17A).

**Nonstructural protein 15 (nsp15) - endoRNAse**

Nsp15 is a Nidoviral RNA uridylate-specific endoribonuclease (NendoU) and its C-terminal is a catalytic domain belonging to the EndoU family of enzymes.(*117–119*) EndoU enzymes have RNA endonuclease activity producing 2'-3' cyclic phosphodiester and 5'-hydroxyl termini.(*120*) A study in 2017 proposed that the NendoU activity interferes with the innate immune response,(*121*) however this finding was disputed in a 2019 study(*122*) that showed that the interference was independent of the endonuclease activity.

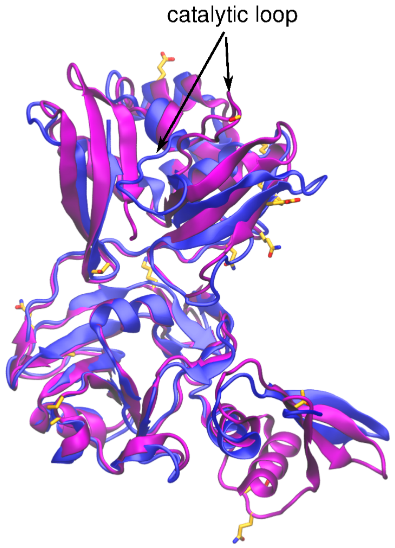


**Fig. S19.** Superimposed crystal structures of SARS-CoV-1 (*blue*) and SARS-CoV-2 (*purple*) nsp15. Non-conservative residues in SARS-CoV-2 nsp15 relative to SARS-CoV-1 are depicted in *orange*.

*Structural analysis and comparison with SARS-CoV-1 nsp15* - The nsp15 monomer (346 a.a.) forms hexamers. Each monomer consists of three domains, namely the N-terminal, middle domain, and catalytic NendoU domain at the C-terminal.(*117*) Within SARS-CoV-1 and SARS-CoV-2, nsp15 is very conserved (89% identity). However, the recently solved crystal structure of the nsp15 monomer, shows a significant conformational variation in the region of the catalytic site. Although several non-conservative substitutions occur around this region, the conformation of the catalytic loop (a.a. 234-249) is known to greatly change upon protein oligomerization (Fig. S19).

**Nonstructural protein 16 (nsp16) - 2'-O-ribose methyltransferase**

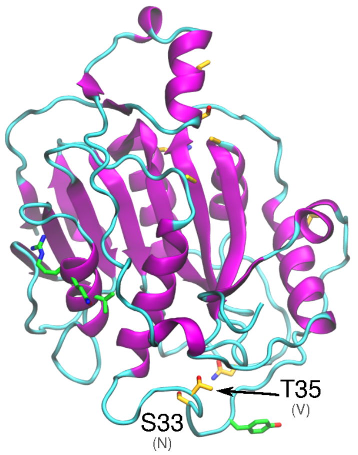


**Fig. S20.** Local modeling-derived structure of SARS-CoV-2 nsp16. Non-conservative substitutions relative to SARS-CoV-1 nsp16 and depicted in *orange* and hey functional residues are depicted in *green*.

The nonstructural protein nsp16 is involved in capping of viral mRNA to protect it from host degradation, and it has been demonstrated that it has to be associated with nsp10 to be active. One potential method of ablating nsp16 activity is to disrupt the binding interface between nsp16 and nsp10. In SARS-CoV-1, the following interfacial mutations in nsp16 were shown to ablate nsp16 2'-O-methyltransferase activity: Ile$^{40}$Ala, Met$^{41}$Ala, Val$^{44}$Ala, Val$^{78}$Ala, Arg$^{86}$Ala, Val$^{104}$Gly, Leu$^{244}$Ala and Met$^{247}$Ala.(*102*) In a separate study, a double mutant (His$^{83}$Ala/Pro$^{84}$Ala) and a triple mutant (Tyr$^{76}$Ala/Cys$^{77}$Ala/Arg$^{78}$Ala) of nsp10 abolished SAM and m7GppA-RNA binding to nsp16(*123*), further supporting the essential role of nsp10 in activating nsp16. In terms of the physicochemical interfacial properties that are essential for activating nsp16, it appears that certain hydrophobic interactions play a critical role and that enhancement of these may increase nsp16. This is exemplified by the Tyr$^{96}$Phe mutation in nsp10 that increases methyltransferase activity by increasing hydrophobic interaction between nsp10 and the hydrophobic pocket of nsp16, which includes the sidechain of Val$^{84}$ and the mainchain portions of Gln$^{87}$ and Arg$^{86}$.(*123*) In SARS-Cov-1, residue Tyr$^{30}$ in the RNA binding site of nsp16, when mutated to either alanine or phenylalanine, abolished methlyltransferase activity, suggesting the critical role of this residue in RNA binding and, consequently, the enzymatic function.(*102*) All these key amino acids are conserved in SARS-CoV-2 nsp16, but non-conservative substitutions in the vicinity of Tyr$^{30}$, namely Asn$^{33}$Ser and Val$^{35}$Thr may affect RNA binding, as well as conservative substitutions (Glu$^{32}$Adp and Ile$^{36}$Leu) that may have a steric effect (Fig. S20).

**Spike glycoprotein** (discussed along the main text)

### ORF3a

ORF3a is the largest of all SARS-CoV accessory proteins and it is expressed in SARS-CoV-1 infected cells, incorporated in viral particles.(*124*) ORF3a is an integral transmembrane protein, localized mostly in the Golgi complex, cytoplasm, and cell surface.(*125–127*) Deletion of several accessory proteins including ORF3a do not dramatically decrease viral replication, suggesting they are not essential for viral growth.(*128*) However, the most significant decrease in viral growth was observed when ORF3a was deleted.(*128*) ORF3a is able to form homodimers via disulfide bridges, which, in turn, form homotetramers through non-covalent interactions.(*127*) The homotetramers have ion channel properties able to transport $Na^+$, $K^+$, and $Ca^{+2}$ ions.(*129*) Although ion channel activity of ORF3a was shown to not be required for viral replication in mouse lungs(*129*), if ORF3a is silenced, viral release decays proportionally to the amount of siRNA used.(*127*)

ORF3a has been demonstrated to interact with M, N and S viral proteins.(*130*) For instance, it forms disulfide bonds with S protein via the cysteine rich motif.(*131*) A number of mutations in ORF3a correlates with mutation in the S protein in different SARS-CoV-1 isolates, suggesting that its function is linked to the S protein.(*131*) Additionally, ORF3a and the E protein are both required for maximum viral growth.(*129*) Deletion of both ORF3a and E protein binding modules yielded no viable viruses, while the presence of only one of them was sufficient to revert the phenotype.(*129*)

Regarding pathogenic effects, ORF3a can trigger inflammatory responses by inducing the expression of NB-κB and IL-8.(*132*) Besides, ORF3a has been shown to be involved in inflammatory responses by promoting activation of NLRP3 inflammasome.(*133*) NLRP3 activation is achieved by the ubiquitination of apoptosis-associated speck-like protein containing a caspase recruitment domain (ASC) facilitated by TNF receptor-associated factor 3 (TRAF3) and ORF3a direct interaction.(*133*) Furthermore, ORF3a triggers unfolded protein response (UPR) in the endoplasmic reticulum via PKR-like ER kinase (PERK) pathway, thereby increasing IFN alpha-receptor subunit 1 (IFNAR1) phosphorylation, ubiquitination and lysosomal degradation.(*134*) Thus, ORF3a may weaken INF responses and innate immunity responses. ORF3a has apoptotic and cell cycle regulation functions via caspase pathways and by arresting cells at G1 stage.(*57*, *135*) Further, if either tyrosine (YXXφ), di-acidic (EXD), or cysteine rich motifs are mutated the apoptotic effects of ORF3a are reduced.(*136*) Furthermore, ORF3a potassium binding activity is blocked, apoptosis is also blocked.(*136*) From a clinical perspective, some patients recovered from SARS showed joint pain and bone damage. Consequently, the study conducted by Obitsu et al.(*137*) showed that ORF3a is able to induce osteoclastogenesis and thus bone abnormalities observed in SARS survivors. Further, some patients developed pulmonary thrombosis during SARS-CoV-1 infection(*138*), while ORF3a was reported to increase expression of all subunits of fibrinogen.(*139*) Taken together, ORF3a may contribute to the formation of blood clots in some patients.

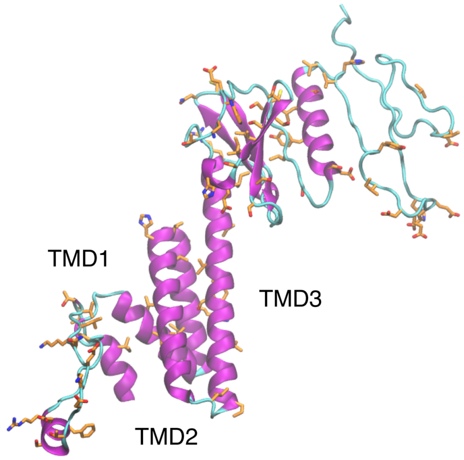

**Fig. S21.** *Ab initio* predicted structure of SARS-CoV-2 ORF3a. Non-conservative substitutions relative to SARS-CoV-1 ORF3a are depicted in *orange*.

*Structural analysis and comparison with SARS-CoV-1 ORF3a* - ORF3a protein is the biggest of all SARS-CoV-1 accessory proteins with 274 a.a. Structurally, ORF3a possess a signal peptide (1 - 15) and three transmembrane domains TMD1 (34 - 56), TMD2 (77 - 99), and TMD3 (103 - 125) and a cytoplasmic domain at its C-terminal of 149 residues.(*131*, *140*, *141*) Several important residues have been identified for ORF3a function. At ORF3a cytoplasmic domain, there is a predicted calcium binding domain at residues 209 - 264 that goes through conformational changes upon binding calcium *in vitro*.(*140*) Besides, a cysteine rich motif (Cys-Trp-Leu-Cys-Trp-Lys-Cys) from residues 127 - 133, located at the end of TMD3 region is essential for ORF3a and S protein interaction.(*131*) The tyrosine motif Tyr-Asn-Ser-Val (160 - 163) was found essential for a proper ORF3a export to the cell surface.(*126*) Mutations at Tyr-Asn-Ser-Val motif, increase ORF3a retention at Golgi complex and lysosome degradation.(*126*) Further, the motifs Tyr-Asn-Ser-Val, di-acidic (Glu-Gly-Asp, 172 - 173) and cysteine rich motifs, have been associated with apoptotic function of ORF3a.(*136*) Mutations at 176 - 274 residues pointed this region to be important to induce cell cycle arrest at G1 stage.(*135*) Regarding subcellular transport, yeast two-hybrid assays showed that caveoline-1 interacts directly to ORF3a N-terminal domain.(*142*) ORF3a contains three caveolin binding sites (Trp-Gln-Leu-Ala-Leu-Tyr-Lys-Gly-Phe, Tyr-Leu-Tyr-Ala-Leu-Ile-Tyr-Phe, and Tyr-Asp-Ala-Asn-Tyr-Phe-Val-Cys-Trp).(*142*) if one or two are mutated ORF3a colocalize still with caveolin-1, suggesting that only one site is enough for this interaction to take place.(*142*) Like M protein, ORF3a is glycosylated and substitutions $Ser^{27}$Gly, $Thr^{28}$Ala, $Thr^{32}$Ala and $Thr^{34}$Ala prevents its glycosylation.(*143*) Further substitutions at TMD2 $Tyr^{91}$Ala and $His^{93}$Ala, and TMD3 $Tyr^{109}$Ala eliminated ORF3a channel activity.(*129*) Besides, residues 125 - 200 are predicted to be β-sheet and are able to bind at the 5'-UTR the viral genomic RNA, which may in turn help with virion assembly.(*144*) Finally, ORF3a contains a PDZ domain-binding motif (PBM, Ser-Val-Pro-Leu, 271 - 274) required for maximum viral yield.(*129*)
SARS-CoV-2 ORF3a is a protein of 275 residues with three transmembrane domains (TMD) and a C-terminal cytoplasmic domain (Fig. S21). Equal to SARS-CoV-1, the predicted transmembrane positions for SARS-CoV-2 ORF3a are located at 34 - 56 (TMD1), 77 - 99 (TMD2) and 103 - 125 (TMD3). In total 76 mutations created a sequence identity of 72·36% between SARS-CoV-1 and SARS-CoV-2. The mutations are distributed along the whole ORF3a sequence, but some conservated zones were observed at TMD1, TMD2, TMD3, cysteine-rich domain and the last portion of C-terminal domain (244 - 275). TMD2 $Tyr^{91}$ and $His^{93}$ as well as TMD3 $Tyr^{109}$ were conserved, reinforcing the idea that these residues are important for ORF3a ion channel activity. One mutation was observed in the cysteine rich motif $Cys^{127}$Leu (Leu-Trp-Leu-Cys-Trp-Lys-Cys). This substitution may affect ORF3a and S protein interaction, but it is likely that the other two cysteines are enough to establish the interaction. The tyrosine motif Tyr-Asn-Ser-Val was fully conserved likely due to its importance for ORF3a function. Besides, the $Glu^{171}$Ser was found in the di-acidic domain (Ser-Gly-Asp). This mutation may have a reductive effect in ORF3a apoptotic feature.(*136*) Truncation of residues 176 - 274 impacts the role of ORF3a in cell cycle arrest.(*135*) Ten mutations were detected at this zone in SARS-CoV-2 ORF3a, but also a conserved motif Asp-Tyr-Gln-Ile-Gly-Gly-Tyr-Thr-Glu with only $Ser^{190}$Thr substitution was observed. Thus, this conserved motif might be enough to carry out cell cycle arrest, but this hypothesis requires proper testing.

The three caveoline-1 binding sites were highly similar (Trp-Gln-Leu-Ala-Leu-Ser-Lys-Gly-Val, Tyr-Leu-Tyr-Ala-Leu-Val-Tyr-Phe and Tyr-Asp-Ala-Asn-Tyr-Phe-Leu-Cys-Trp) with only one non-conservative substitution Tyr$^{74}$Ser and two conservative substitutions, Ile$^{112}$Val and Val$^{147}$Leu. The Tyr$^{74}$Ser substitution led us to hypothesize that residue could be a potential phosphorylation site. Although *in silico* analysis of phosphorylation yielded 29 putative sites, phosphorylation of ORF3a has not been demonstrated yet. Non-conservative substitutions were found in two out of four putative glycosylation sites (Ser$^{27}$Asp and Thr$^{28}$Phe), meanwhile Thr$^{32}$ and Thr$^{34}$ were conserved. This observation suggests that glycosylation of ORF3a may occur in these residues. Furthermore, Thr$^{32}$ and Thr$^{34}$ are found in a conserved motif (SARS-CoV-1 Val-His-Ala-Thr-Ala-Thr-Ile-Pro and SARS-CoV-2 Val-Arg-Ala-Thr-Ala-Thr-Ile-Pro) where only one conserved substitution Arg$^{30}$His was found.
The ORF3a area from 125 to 200 is required for binding to the 5'-UTR of viral genomic RNA.(*144*) Although residues 125 - 200 is a well conserved region, several mutations were observed in this area. To address the question if any of these mutations may have an effect on ORF3a RNA binding, we paid attention to know nucleic acid binding residues arginine, lysine and histidine. Thus, one conservative Lys$^{134}$Arg and non-conservative His$^{152}$Asn, Lys$^{179}$Ile, Lys$^{181}$Glu, Arg$^{193}$Trp and His$^{194}$Glu were found. However, close to these mutations were found also others in their proximity that could replace their function. For instance, close to His$^{152}$Asn there is a conserved histidine at position 150, next to Lys$^{179}$Ile and Lys$^{181}$Glu there is Glu$^{182}$His, and next to Arg$^{193}$Trp and His$^{194}$Glu a Asp$^{192}$Lys substitution was observed. These new substitutions suggest that ORF3a could retain its RNA binding properties. However, this hypothesis requires empirical validation. Finally, the PBM was also found in SARS-CoV-2 ORF3a with identical sequence to SARS-CoV-1.

**Envelope protein**

The envelope (E) protein is one of the four structural proteins of coronaviruses. The other structural proteins are the membrane (M), nucleocapsid (N) and spike (S) proteins. The E protein is a type 1 transmembrane protein that is able to form pentamers by associating with other E proteins.(*145*) This pentamer forms a membrane pore that is able to transport ions.(*145*) The pore structure is called viroporin and it is present in many common viruses. The ion channel activity can be inhibited in SARS-CoV-1 E protein by the drug HMA .(*145*) Only a few E proteins are present in each viral particle, but it is highly expressed in the host cells.(*146*, *147*) The E protein has been proposed to initiate membrane curvature together with the M protein via the interaction between their C-terminal domains, but this mechanism remains largely unknown.(*148*, *149*) The M protein alone seems to not trigger a proper membrane curvature for virion production. The M protein alone can produce virions, but the absence of the E protein cripples virion production, morphology, and plaque shape.(*147*, *149–151*) The E protein localizes mainly in the ER and Golgi apparatus, where it participates in assembly, budding, and intracellular trafficking of newly formed virions.(*147*, *149*)
The E protein has been shown to interact with other viral proteins. Tandem affinity purification assays established interaction between the S and the E proteins, but the mechanism of how this happens was not pursued further.(*152*) This study also shows that the E protein interacts with the nsp3 and they suggest that nsp3 mediates E protein ubiquitination. Besides, the E protein co-immunoprecipitates with the N protein, but the function of this interaction remains unclear.(*153*) Furthermore, a yeast two-hybrid system and an *in vitro* pull down assay showed interaction

between E proteins and ORF7a, but its importance is yet to be identified.(*154*, *155*) On the other hand, the E protein PDZ-Binding Motif (PBM) interacts with PDZ domains of host proteins. For instance, the interaction of the E protein and a protein associated with *Caenorhabditis elegans,* lin-7 protein 1 (PALS1), a PDZ-containing protein, disrupts tight junctions in the lungs to reach the alveolar wall and develops into a systemic infection.(*156*) Further, the interaction with syntenin caused it to concentrate in the cytoplasm, triggering an overexpression of inflammatory cytokines, which activates an exaggerated immune response, resulting in lung tissue damage, edema accumulation, and leading to acute respiratory distress syndrome.(*157*) Interaction between the E protein and the Bcl-xL protein caused lymphopenia.(*158*)

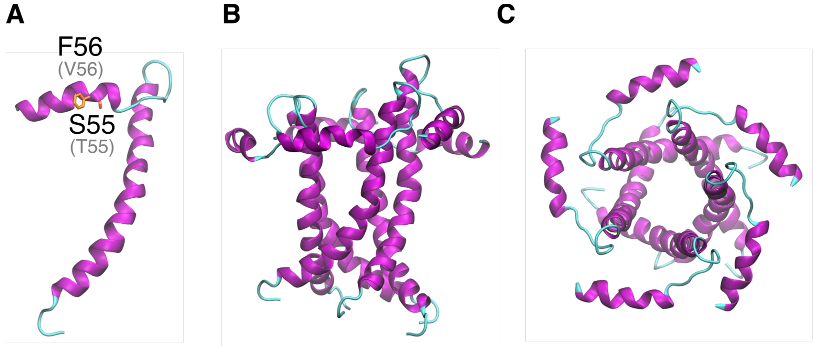


**Fig. S22.** Cartoon representation of SARS-CoV-2 E protein. E protein monomer and substitutions in the C-terminal relative to SARS-CoV-1s (A). E protein pentamer front (B) and top views (C).

*Structural analysis and comparison with SARS-CoV-1 envelope protein* - Several functionally important residues in E protein have been identified in SARS-CoV-1. For instance, there is a conserved proline residue in the C-terminal in the β-coil-β motif, that if mutated changes localization of the E protein from the Golgi complex to the plasma membrane.(*145*, *159*) Besides, the mutations $Asn^{15}Ala$ and $Val^{25}Phe$ inhibit ion channel activity of SARS-CoV-1 E protein. After several passages through cell cultures, this function is restored by the addition of new mutations, suggesting that the E protein confers a selective advantage to the virus.(*160*) It is also important to note that the C-terminal of the E protein interacts with the C-terminal of the M protein in the cytoplasmic side, and this is required to virus envelope formation.(*149*) As being highly conserved relative to SARS-CoV-1 (96% identical), this information is likely applicable to SARS-CoV-2 E protein.

SARS-CoV-2 E protein has a predicted short 11 aa N-terminal tail, a 25 aa transmembrane region and a 37 aa C-terminal cytoplasmic region, including PBM (DLLV). Four variations relative to SARS-CoV-1 E protein were verified, all located in the C-terminal end (Fig. S22A), including in one conservative substitution, two non-conservative substitutions ($Val^{56}Phe$, $Glu^{69}Arg$), and a deletion. Similar to SARS-CoV-1, SARS-CoV-2 E protein is likely to assemble as a pentamer to form a viroporin (Fig. S22B-C), and the three C-terminal substitutions are likely exposed in the pentamer to the cytoplasm, thereby they could be involved in modulating E protein interaction with other proteins.

### Membrane protein

Membrane (M) proteins are the most abundant of all coronavirus structural proteins, and their presence and conformation settle the virion shape.(*161*, *162*) M protein has been shown to interact with viral proteins N, S, ORF3a and ORF7a.(*130*, *155*, *163*, *164*) The general structure of M proteins are that they contain a short N-terminal ectodomain, three transmembrane domains (TMD), and a C-terminal endodomain.(*165*) M proteins may work as homodimers that can take compact and long conformations.(*166*) The long conformation is associated with high S protein density and efficient virus budding, while compact conformation is associated with patchy and low S protein presence and inefficient budding.(*166*) Only M and E proteins are required to form virus-like particles (VLP), but absence of S proteins makes VLPs appear larger, suggesting they may contribute to virus formation.(*166*) SARS-CoV-1 M protein has been shown to interact with N protein in mammalian two-hybrid assays.(*167*) Only RNAs interacting with M protein can be found in viral particles in mouse hepatitis virus (MHV).(*161*)

The M protein has been demonstrated to be a pathogenic factor in coronaviruses. For instance, overexpression of M protein *in vitro* and *in vivo* induces apoptosis in human cell culture and *Drosophila*, respectively.(*168*) Further, SARS-CoV-1 M protein was found to interact with IKKβ, impairing NF-κB signaling and reducing the expression of cyclooxygenase-2 (COX-2).(*169*) COX-2 is responsible for the production of prostaglandins, which in turn trigger inflammatory responses. Furthermore, interferon (INF) are a group of antiviral proteins produced by cells in response to a viral infection. M protein impairs the production of type-I INF by interfering with TRAF3-TANK-TBK1/IKKε complex formation, thus preventing IRF3 phosphorylation.(*170*) Nonetheless, antigenic regions of M protein are detected and production of INF is triggered.(*171*) These antigenic properties were observed in patients after a year of recovery from SARS-CoV-1 infection, showing persistent IFNγ release from CD4+ and CD8+ lymphocytes when cocultured with M peptides.(*172*) Thus, SARS-CoV-1 infection triggered the production of memory cells and M protein could be an interesting vaccine candidate.

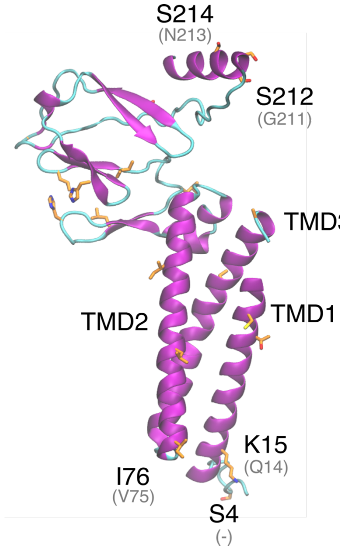


**Fig. S23.** *Ab initio* predicted structure of the SARS-CoV- 2 M protein. Conserved predicted regions and non-conservative substitutions (*black*) relative to SARS-CoV-1 (*grey*) are depicted in *orange*.

*Structural analysis and comparison with SARS-CoV-1 membrane protein* - Several key functional residues have been identified for the M protein. Transmissible gastroenteritis virus (TEGV) showed reduced IFN induction when mutations occur in the N-terminal residues 6 - 22.(*171*) Alteration of the Asn-Ser-Thr motif in this region prevented glycosylation of the M protein, suggesting its importance in INF induction.(*171*) In another study, after several subculture

passages in immortalized proximal tubular epithelial cells (PTEC), the SARS-CoV-1 M protein generated a glutamic acid to alanine point mutation at position 11.(*173*) This mutation enhances virus replication and persistence in PTEC.(*173*) Tseng *et al.*(*174*) identified other key residues in the SARS-CoV-1 M protein with different functions in replication. The Leu-Leu motif (Leu$^{218}$ and Leu$^{219}$) is required to incorporate the N protein into VLPs. Substitution Cys$^{158}$Ser reduces secretion of the M protein, but not Cys$^{63}$Ser or Cys$^{85}$Ser, and Cys$^{158}$ is involved in the interaction with the N protein. Non conservative substitutions in the motif Ser-Trp-Trp-Ser-Phe-Asn-Pro-Glu reduced the production of VLPs. Conversely, substitutions of the aromatic residues in this motif by analogs did not affect the VLPs production, suggesting that the conservation of these aromatic amino acids is critical for M protein function.
The SARS-CoV-2 M protein is predicted to have three transmembrane domains, a cytoplasmic and a non-cytoplasmic domain. This is similar to the SARS-CoV-1 M protein and other coronavirus M proteins.(*165*, *175*) InterProScan(*176*) defined these regions as follows: non-cytoplasmic region (1 - 19), TMD1 (20 - 40), cytoplasmic region (41 - 51), TMD2 (52 - 73), non-cytoplasmic region (74 - 78), TMD3 (79 - 103) and cytoplasmic region (103 - 222). The SARS-CoV and SARS-CoV-2 M proteins are very similar, with 90·54% identity. Among 21 variations, 12 are non-conservative substitutions and one is an insertion. Within the substitutions, Gln$^{15}$Lys may have an effect in antigenic properties of the M protein, by modifying its ectodomain (Fig. S23). Besides, Ser$^{4}$ insertion is very close to a predicted N-glycosylation site at the N-terminal, and can constitute an additional glycosylation site. Modifications to this glycosylation site have been associated with induction of INF production.(*171*) Most of the non-conservative mutations occur in the cytoplasmic domain at the C-terminal domain. These substitutions involve several sites that can potentially undergo post-translational modification (Ser$^{155}$His, Thr$^{189}$Gly, Asn$^{197}$Ser, Ala$^{211}$Ser, Gly$^{212}$Ser and Asn$^{214}$Ser).

### ORF6

ORF6 is an auxiliary protein in SARS coronaviruses that is not required for virus replication.(*128*, *177*) However, it can increase virus replication when expressed in a heterologous system or at low multiplicity of infection.(*178*) ORF6 localizes in the perinuclear and ER zones, associated with membranes and colocalized with M , S, and N structural proteins.(*179*, *180*) ORF6 is known to interact with the viral proteins ORF9b and nsp8.(*181*, *182*) Furthermore, ORF6 and nsp8 colocalize in cell culture, suggesting ORF6 may play a role in virus replication.(*181*)

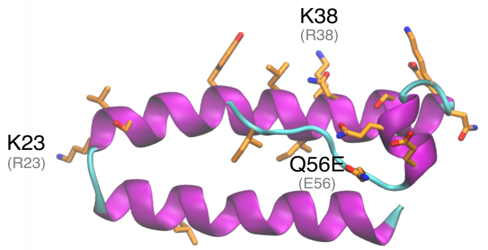


**Fig. S24.** Cartoon representation of SARS-CoV-2 ORF6. Substitutions (*balck*) relative to SARS-CoV-1 (*grey*) are shown in *orange*.

*Structural analysis and comparison with SARS-CoV-1 ORF6* - SARS-CoV-1 ORF6 is a small protein of 63 amino acids and 7·53 kDa. Several important residues have been related to SARS-CoV-1 ORF6 structure, function, and location. For instance, ORF6 sequence suggests a membrane

association in residues 7 - 37.(*180*) Even though some residues in this region are charged (Glu$^{13}$, Arg$^{20}$ and Lys$^{23}$, in SARS-CoV-2 ORF6), this was enough for membrane association.(*180*)
ORF6 prevents import of the host signal transducer and activator of transcription 1-alpha/beta (STAT1) to the nucleus by tethering karyopherin subunit alpha-2 (KPNA2) and, therefore, KPNA1 to the ER and golgi apparatus membrane, thereby impairing the activation of INF-induced genes.(*183*) Tests with specific mutations in the C-terminal region of ORF6 revealed specific residues that affect disruption of KPNA2 ER and Golgi arrest in ORF6 transfected cells.(*183*) Further, the ORF6 C-terminal was shown to interact with Nmi protein, mediating Nmi ubiquitination and proteasome degradation, thus suppressing INF signaling.(*184*) ORF6 N-terminal, in turn, was shown to induce membrane rearrangements typically observed in virus-infected cells and to have critical importance to prevent STAT1 translocation to the nucleus.(*185*)
SARS-CoV-1 and SARS-CoV-2 ORF6 proteins are 68·85% identical. Most of the substitutions are located in the C-terminal helix (Fig. S24). Lysine substitutions Arg$^{23}$Lys, Arg$^{38}$Lys and Asn$^{38}$Lys in SARS-CoV-2 ORF6 suggest the introduction of new putative ubiquitination sites. ORF6 has been shown to be involved in proteasomal degradation of Nmi, but if ORF6 itself is regulated by the proteasome system is unknown.
Two studies pointed to residues 53 - 63 to be important for SARS-CoV-1 ORF6 function.(*183*, *185*) In this region, one non-conservative substitution (Glu$^{56}$Gln) and two deletions (Tyr$^{62}$ and Pro$^{63}$) are verified.

**ORF7a**

ORF7a is an accessory protein of coronaviruses and it is not essential for viral replication *in vitro.(186)* ORF7a is a type I transmembrane protein, localized mainly in Golgi apparatus and in the cell surface.(*187*, *188*) Besides, ORF7a colocalizes with calnexin (ER marker), showing that it is also localized at ER.(*187*)
ORF7a is an antagonist of bone marrow stromal antigen 2 (BST-2/CD317/tetherin).(*188*) BST-2 is a pre-B-cell growth promoter that inhibits virus release by tethering budding virions to the host cell membrane.(*189*) A greater virion tethering to the cell membrane is observed when ORF7a is not present.(*188*) ORF7a is usually located in the Golgi apparatus and it relocates to the plasma membrane when BST-2 is expressed, colocalizing it.(*188*) ORF7a binds to BST-2 and reduces restriction activity of BTS2 by preventing its glycosylation.(*188*) Further *in vitro* experiments showed that ORF7a is able to induce apoptosis in a caspase dependent manner.(*186*)

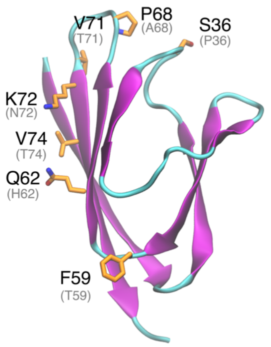


**Fig. S25.** Cartoon representation of SARS-CoV-2 ORF7a luminal domain. Substitutions (*black*) relative to SARS-CoV-1 (*grey*) are shown in *orange*.

*Structural analysis and comparison with SARS-CoV-1 ORF7a* - SARS-CoV-2 ORF7a protein structure has a 15 amino acid (a.a.) N-terminal signal peptide, a 80-a.a. luminal domain, a 21-a.a. transmembrane domain, and a 5-a.a. cytoplasmic tail.(*187*) Residues $Lys^{117}$, $Asn^{118}$ and $Lys^{119}$ form a motif that has been described as being recognized by COPII vesicular system implicated in the transport of proteins from endoplasmic reticulum (ER) to Golgi and they are required to exit the ER.(*190*)
ORF7a structure is composed of seven antiparallel β-sheets that altogether make a β-sandwich (Fig. S25). The ORF7a is very conserved between SARS-CoV-1 and SARS-CoV-2 with 85% sequence identity. A total of 18 variations are verified, with 11 non-conservative substitutions and a deletion. Substitutions in the luminal domain are located in the protein surface (Fig. S25). Therefore, the ORF7a fold is conserved, but these substitutions may affect ORF7a interaction with other viral or host proteins. Mostly conservative mutations are verified at ORF7a C-terminal (83 - 121), except by $Ile^{111}Thr$.

**ORF7b**
ORF7b is a small accessory protein expressed in SARS-CoV-1 and SARS-CoV-2 infected cells with no sequence homology with other viral proteins.(*191*) It is translated from a bicistronic open reading frame and encoded in the subgenomic RNA 7 and translated by ribosome leaky scanning.(*191*) Like other accessory proteins, ORF7b is incorporated in viral particles and detected in purified virions.(*191*) ORF7b is an integral transmembrane protein and it localizes in the cis- and trans- Golgi.(*192*) Deletion of gene 7 does not affect replication kinetics in vitro, suggesting that ORF7b is not essential for virus replication.(*193*)

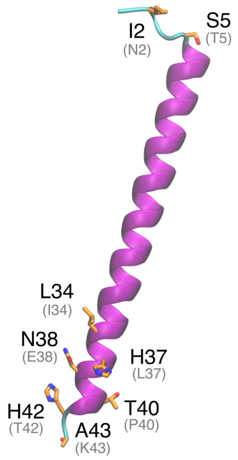


**Fig. S26.** Cartoon representation of SARS-CoV-2 ORF7b protein. Substitutions (*balck*) relative to SARS-CoV-1 (*grey*) are shown in *orange*.

*Structural analysis and comparison with SARS-CoV-1 ORF7b* - SARS-CoV-1 ORF7b is a small protein of 44 amino acids with a transmembrane domain (TMD) of 22 residues.(*192*) Mutations in the TMD affect ORF7b cellular localization.(*192*) Alanine scanning experiments, identified that residues 13-15 and 19-22 are critical for ORF7b retention in the Golgi complex. SARS-CoV-2 ORF7b is 81% identical to SARS-CoV-1 ORF7b. All substitutions are found in the terminals, meaning the TMD is fully conserved (Fig. S26).

### ORF8

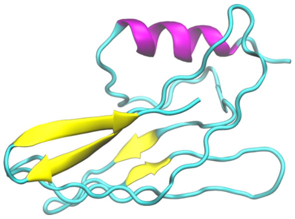

**Fig. S27.** *Ab initio* predicted structure of SARS-CoV-2 ORF8.

SARS-CoV-2 ORF8 is most similar to sequences from Bat-SARS-like coronavirus and a bat coronavirus RaTG13.(*194*) Remarkably, removal of a N-terminal 29 nucleotide sequence from SARS-CoV-1 ORF8 in early tests showed a decrease in viral replication by up to 23-fold.(*195*) This sequence was observed to be removed from the virus in human cases later in the outbreak along with additional and different mutations of ORF8.(*196*, *197*) The exact purpose of this mutation and ORF8's function in regards to viral benefit within human hosts remains unknown. With the deletion, SARS-CoV-1 encodes ORF8a and ORF8b, while SARS-CoV-2 ORF8 is intact.(*198–201*) SARS-CoV-2 ORF8 (Fig. S27) is 127 a.a. long and 45% identical to SARS-CoV-1 ORF8b.

### Nucleocapsid protein

Highly abundant in infected cells, SAR-CoV-1 nucleocapsid (N) packages the viral RNA into a helical ribonucleocapsid and plays a key role in viral assembly.(*202*) The N protein is assembled and organized in a modular fashion and has been shown to bind to viral RNA at multiple sites in an allosteric manner.(*203*) The modularity of the N protein is thought to increase selectivity through coupled allosteric binding of individual nucleocapsid domains, aid in regulation and functional expression, and independently evolve binding sites.(*203*)

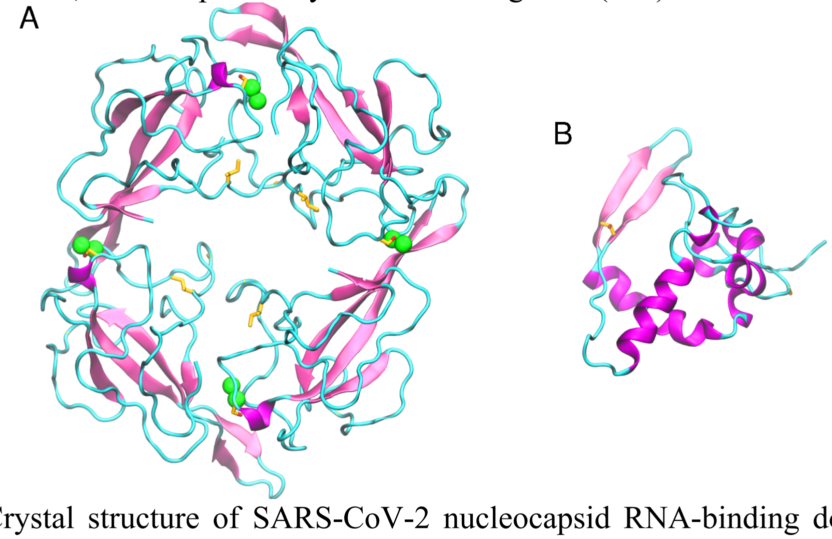


**Fig. S28.** A) Crystal structure of SARS-CoV-2 nucleocapsid RNA-binding domain. B) Local modeling-derived structure of SARS-CoV-2 nucleocapsid C-terminal. Non-conservative substitutions relative to SARS-CoV-1 nucleocapsid are depicted in *orange*.

*Structural analysis and comparison with SARS-CoV-1 nucleocapsid* - SARS-CoV-2 N protein is a 419 a.a. long protein and is 89% identical to SARS-CoV-1 nucleocapsid. In SARS-CoV-1, a Ser-Arg rich motif (Ser-Ser-Arg-Ser-Ser-Ser-Arg-Ser-Arg-Gly-Asn-Ser-Arg) was found to be important for the oligomerization of N proteins.(*164*) This motif is mostly conserved in SARS-CoV-2, except by the substitutions $Gly^{192}Asn$ and $Asn^{193}Ser$. The RNA binding domain of the N protein is located at the N-terminal, within residues 47 and 180 of SARS-CoV-2. The structure of this domain recently solved (PDB 6vyo) (Fig. S28A). Residues within the N-terminal of the SARS-CoV-1 nucleocapsid, namely, $Tyr^{87}$, $Tyr^{110}$, $Tyr^{112}$, $Y^{113}$, $Leu^{122}$, and $Ala^{13}$, are hypothesized to play a role in ribonucleocapsid packaging.(*203*) All of these residues are fully conserved in SARS-CoV-2 N. A long intrinsically disordered region follows these residues (181-246), and is predicted to fold upon binding (ANCHOR2 prediction(*12*)).
The oligomerization domain of nucleocapsids is situated in the C-terminal of SARS-CoV-1 nucleocapsid. Residues $Trp^{302}$, $Ile^{305}$, $Pro^{310}$, $Phe^{315}$, and $Phe^{316}$ are thought to be associated with highly hydrophobic interactions between two helices within the nucleocapsid(*203*), and they are fully conserved in SARS-CoV-2 N (Fig. S28B).

**ORF9**

ORF9 is an accessory protein synthesized from an alternative reading frame in the N gene. This accessory protein is integrated in viral particles and therefore it can be considered a structural protein. This incorporation occurs in presence of E and M proteins, suggesting a potential interaction among these proteins.(*97*) ORF9 localizes in mitochondria outer membrane, cytoplasm, nucleus, endoplasmic reticulum and lipid vesicles.(*182*, *204–206*) At mitochondrial level, ORF9 promotes the ubiquitination and degradation of dynamin-like protein (DRP1), thereby causing mitochondria to show an elongated phenotype.(*204*) Here, ORF9 also interacts with mitochondrial antiviral signaling protein (MAVS) and poly(rC) binding protein 2 (PCBP2).(*204*) Thus, PCBP2 facilitates the ubiquitination of MAVS by AIP4, suppressing the activation of INF regulatory factors and NF-κB.(*204*) Besides, interaction and colocalization between ORF9 and ORF6 have been suggested(*182*), but the relevance of this interaction remains unknown.

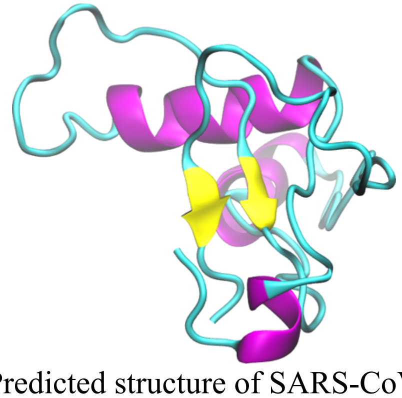

**Fig. S29.** Predicted structure of SARS-CoV-2 ORF9.

*Structural analysis and comparison with SARS-CoV-1 ORF9* - SARS-CoV-1 ORF9 is a small protein of 98 residues length. Meier et al.(*206*) showed that ORF9 forms a symmetric dimer where the monomer interactions resemble a handshake. Dimer assembling creates a hydrophobic tunnel that can accommodate a long fatty acid chain. The authors suggest that ORF9 could anchor itself to a lipidic membrane by internalizing one or more lipidic tails. SARS-CoV-2 ORF9 (Fig. S29) shares high homology with SARS-CoV-1 (72·45% protein identity).

**ORF10**

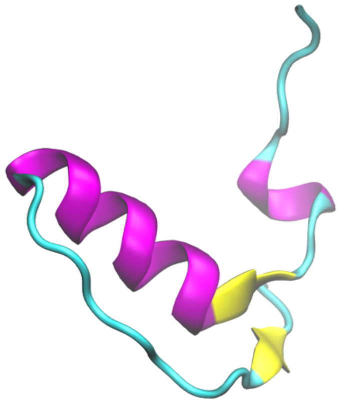

**Fig. S30.** Predicted structure of ORF10.

Within SARS-CoV-2, ORF10 is thought to not have any functional protein purpose as it was not found within the sub genomic mRNA sequencing.(*207*) It's hypothesized that this protein may act itself or as a precursor of additional RNAs in roles concerning gene expression, controlling cellular antiviral pathways, or within viral replication.(*207*) Another hypothesis is that this protein, if not an artifact, could be involved with regulating molecular function or contributing to response stimulus based on DeepGOPlus Gene Ontology.(*208*) In another study, based on the SARS-CoV-2 interactome, ORF10 is suggested to interact with a Cullin 2 RING E3 ligase complex.(*201*) Potentially, ORF10 might bind specifically to Cullin 2 ZYG11B complex and hijack this complex for ubiquitination and degradation. It appears to localize only within the extracellular region of the host.(*208*)

*Structural analysis of SARS-CoV-2 ORF10* - ORF10 of SARS-CoV-2 is the last predicted coding sequence upstream of the poly-A tail and is the shortest predicted coding sequence, composed of 38 a.a.(*207*) ORF10 is predicted to harbor a long helix and a pair of б-strands. ORF10 is not found within the SARS-CoV-1 proteome.(*207*) It appears to be unique to SARS-CoV-2 (Fig. S30).

201. D. E. Gordon, G. M. Jang, M. Bouhaddou, J. Xu, K. Obernier, M. J. O'Meara, J. Z. Guo, D. L. Swaney, T. A. Tummino, R. Huttenhain, R. M. Kaake, A. L. Richards, B. Tutuncuoglu, H. Foussard, J. Batra, K. Haas, M. Modak, M. Kim, P. Haas, B. J. Polacco, H. Braberg, J. M. Fabius, M. Eckhardt, M. Soucheray, M. J. Bennett, M. Cakir, M. J. McGregor, Q. Li, Z. Z. C. Naing, Y.

Zhou, S. Peng, I. T. Kirby, J. E. Melnyk, J. S. Chorba, K. Lou, S. A. Dai, W. Shen, Y. Shi, Z. Zhang, I. Barrio-Hernandez, D. Memon, C. Hernandez-Armenta, C. J. P. Mathy, T. Perica, K. B. Pilla, S. J. Ganesan, D. J. Saltzberg, R. Ramachandran, X. Liu, S. B. Rosenthal, L. Calviello, S. Venkataramanan, J. Liboy-Lugo, Y. Lin, S. A. Wankowicz, M. Bohn, P. P. Sharp, R. Trenker, J. M. Young, D. A. Cavero, J. Hiatt, T. Roth, U. Rathore, A. Subramanian, J. Noack, M. Hubert, F. Roesch, T. Vallet, B. Meyer, K. M. White, L. Miorin, O. S. Rosenberg, K. A. Verba, D. Agard, M. Ott, M. Emerman, D. Ruggero, A. Garcí-Sastre, N. Jura, M. von Zastrow, J. Taunton, A. Ashworth, O. Schwartz, M. Vignuzzi, C. d'Enfert, S. Mukherjee, M. Jacobson, H. S. Malik, D. G. Fujimori, T. Ideker, C. S. Craik, S. Floor, J. S. Fraser, J. Gross, A. Sali, T. Kortemme, P. Beltrao, K. Shokat, B. K. Shoichet, N. J. Krogan, A SARS-CoV-2-Human Protein-Protein Interaction Map Reveals Drug Targets and Potential Drug-Repurposing. *bioRxiv* (2020), p. 2020.03.22.002386.
